# A Spatial Framework for Understanding Population Structure and Admixture

**DOI:** 10.1101/013474

**Authors:** Gideon S. Bradburd, Peter L. Ralph, Graham M. Coop

## Abstract

Geographic patterns of genetic variation within modern populations, produced by complex histories of migration, can be difficult to infer and visually summarize. A general consequence of geographically limited dispersal is that samples from nearby locations tend to be more closely related than samples from distant locations, and so genetic covariance often recapitulates geographic proximity. We use genome-wide polymorphism data to build “geogenetic maps,” which, when applied to stationary populations, produces a map of the geographic positions of the populations, but with distances distorted to reflect historical rates of gene flow. In the underlying model, allele frequency covariance is a decreasing function of geogenetic distance, and nonlocal gene flow such as admixture can be identified as anomalously strong covariance over long distances. This admixture is explicitly co-estimated and depicted as arrows, from the source of admixture to the recipient, on the geogenetic map. We demonstrate the utility of this method on a circum-Tibetan sampling of the greenish warbler (*Phylloscopus trochiloides*), in which we find evidence for gene flow between the adjacent, terminal populations of the ring species. We also analyze a global sampling of human populations, for which we largely recover the geography of the sampling, with support for significant histories of admixture in many samples. This new tool for understanding and visualizing patterns of population structure is implemented in a Bayesian framework in the program SpaceMix.

**Author Summary:** In this paper, we introduce a statistical method for inferring, for a set of sequenced samples, a map in which the distances between population locations reflect genetic, rather than geographic, proximity. Two populations that are sampled at distant locations but that are genetically similar (perhaps one was recently founded by a colonization event from the other) may have inferred locations that are nearby, while two populations that are sampled close together, but that are genetically dissimilar (e.g., are separated by a barrier), may have inferred locations that are farther apart. The result is a “geogenetic” map in which the distances between populations are effective distances, indicative of the way that populations perceive the distances between themselves: the “organism’s-eye view” of the world. Added to this, “admixture” can be thought of as the outcome of unusually long-distance gene flow; it results in relatedness between populations that is anomalously high given the distance that separates them. We depict the effect of admixture using arrows, from a source of admixture to its target, on the inferred map. The inferred geogenetic map is an intuitive and information-rich visual summary of patterns of population structure.

## Introduction

There are many different methods to learn how population structure and demographic processes have left their mark on patterns of genetic variation within and between populations. Model-based approaches focus on developing a detailed view of the migrational history of a small number of populations, often assuming one or a small number of large, randomly mating populations (i.e. little or no geographic structure). There has been considerable recent progress in this area, using a variety of summaries such as the allele frequency spectrum [Gutenkunst et al., 2009, Bhaskar et al., 2014, Excoffier et al., 2013], or approximations to the coalescent applied to sequence data [Paul et al., 2011, Li and Durbin, 2011, Schiffels and Durbin, 2014].

Other approaches are designed only to visualize patterns of genetic relatedness and population structure, without using a particular population genetic model. Such methods can deal with many populations or individuals as the unit of analysis. Examples of this second set of methods include clustering methods [Pritchard et al., 2000, Alexander et al., 2009, Lawson et al., 2012] and reduced dimensionality representations of the data, such as Principal Components Analysis (PCA; e.g. [Cavalli-Sforza et al., 1994, Patterson et al., 2006, Price et al., 2006]).

A third set of methods that describe relatedness between populations by constructing a “population phylogeny” was pioneered by Cavalli-Sforza and Edwards [1967], as were methods to test whether a tree is a good model of population history [Cavalli-Sforza and Piazza, 1975] (see Felsenstein [1982] for a review). Tree-based approaches are appealing because trees are easy to visualize and explain, but the underlying assumptions (unstructured populations that split at discrete points in time) rarely hold true.

Recently, there has been a resurgence of interest in these tree-based methods. Some use population trees as a null model to test and quantify the signal of admixture between samples [Reich et al., 2009]. Others, such as TreeMix [Pickrell and Pritchard, 2012] and MixMapper [Lipson et al., 2013], visualize population relationships using a directed acyclic graph; for instance, TreeMix connects branches in a population tree with additional edges to explain excess covariance between groups of populations.

There has also been renewed interest in methods for dimensionality reduction for the visualization of patterns of genetic variation [Patterson et al., 2006], especially especially Principal Components Analysis (PCA; also pioneered by Cavalli-Sforza [Menozzi et al., 1978]). Examining such low-dimensional visual summaries has become an indispensable step in the analysis of modern genomic datasets of thousands of loci typed in tens or hundreds of samples. Generally, these visualizations are constructed by plotting the first few eigenvectors of the covariance matrix of normalized allele frequencies against each other.

Both PCA and tree-based methods are valuable as genetic inference and visualization tools, but both also suffer from serious limitations. Because gene flow is frequently pervasive, patterns of relatedness between samples may often be only poorly represented by a tree-based model. PCA is more flexible, as it assumes no explicit model of population-genetic processes, simply describing the axes of greatest variance in the average coalescent times between pairs of samples [McVean, 2009]. This allows PCA to describe more geographically continuous relationships: applied to human populations within continents, it often shows a close correspondence to geographic locations [e.g. Novembre et al., 2008, Wang et al., 2012]. However, the interpretation of PCA is more difficult, as the results can be strongly affected by the size and design of sampling, and the linearity and orthogonality requirements of the PC axes can lead to counterintuitive results [Novembre and Stephens, 2008, Francois et al., 2010, Frichot et al., 2012].

What is desired, then, is a method for inferring and visualizing patterns of population differentiation that can recapitulate complex, non-hierarchical structures, while also admitting simple and intuitive interpretation. Since gene flow and population movements are often constrained by geography, it is natural to base such a method in a geographic framework. There is a rich history of population genetics theory for populations distributed in continuous space [Malécot, 1975, Nagylaki, 1978, Felsenstein, 1975, Barton et al., 2002], as well as exciting new developments in the field [e.g. Petkova et al., 2014]. The pattern of increasing genetic differentiation with geographic distance was termed “Isolation by Distance” by Wright [1943], and is ubiquitous in natural populations [Meirmans, 2012]. Descriptive models of such patterns rely only on the weak assumption that an individual’s mating opportunities are spatially limited by dispersal; a large set of models, ranging from equilibrium migration-drift models to non-equilibrium models, such as recent spatial expansions of populations, give rise to the empirical pattern of isolation by distance.

In this paper, we present a statistical framework for studying the spatial distribution of genetic variation and genetic admixture based on a flexible parameterization of the relationship between genetic and geographic distances. Within this framework, the pattern of genetic relatedness between the samples is represented by a map, in which inferred distances between samples are proportional to their genetic differentiation, and long distance relatedness (in excess of that predicted by the map) is modeled as genetic admixture. These ‘geogenetic’ maps are simple, intuitive, low-dimensional summaries of population structure, and provide a natural framework for the inference and visualization of spatial patterns of genetic variation and the signature of genetic admixture. The implementation of this method, SpaceMix, is available at https://github.com/gbradburd/SpaceMix.

## Results

### Data

The genetic data we model consist of allele counts at *L* unlinked, bi-allelic single nucleotide polymorphisms (SNPs), sampled across *K* populations. After arbitrarily choosing an allele to count at each locus, denote the number of counted alleles at locus *l* in population *k* as *C*_*k,f*_, and the total number of alleles observed as *S*_*k,l*_. The sample frequency at locus *l* in population *k* is 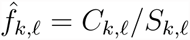. Although we will refer to “populations”, each could consist of a single individual (*S*_*k,l*_ = 2 for a diploid). We will depict results as coordinates on a map; however, the method does not require user-specified sampling locations.

We first compute standardized sample allele frequencies at locus *l* in population *k*, by

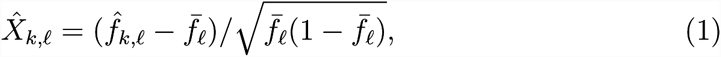

where 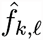 is the sample allele frequency at locus *l* in population *k*, and 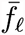 is the average of the *K* sample allele frequencies, weighted by mean population size. This normalization is widely used [e.g. Nicholson et al., 2002, Patterson et al., 2006]; j mean-centering makes the result invariant to choice of which allele to count at each locus, and dividing by 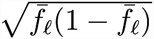 makes each locus have roughly unit variance if the amount of drift since a common ancestor is small.

We work with the empirical covariance matrix of these standardized sample allele frequencies, calculated across loci, namely, 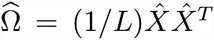. Using the sample mean to mean-center 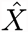 has implications on their covariance structure, discussed in the Methods (“The standardized sample covariance”). For clarity, here we proceed as if 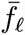 were instead an unobserved, global mean allele frequency at locus *l*.

### Spatial Covariance Model

We wish to model the distribution of alleles among populations as the result of a spatial process, in which migration moves genes locally on an unobserved landsape. Migration homogenizes those differences between populations that arise through genetic drift; populations with higher levels of historical or ongoing migration share more of their demographic history, and so have more strongly correlated allele frequencies.

We assume that the standardized sample frequencies are generated independently at each locus by a spatial process, and so have mean zero and a covariance matrix determined by the pairwise geographic distances between samples. To build the geogenetic map, we arbitrarily choose a simple and flexible parametric form for the covariance matrix in which covariance between allele frequencies decays exponentially with a power of their distance [Diggle et al., 1998, Wasser et al., 2004, Bradburd et al., 2013]: the covariance between standardized population allele frequencies (i.e. 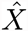 values) between populations *i* and *j* is assumed to be, for *i* ≠ *j*,

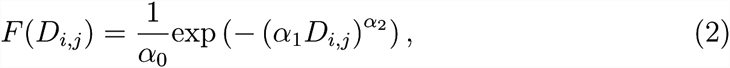

where *D*_*i,j*_ is the geogenetc distance between populations *i* and *j*, *α*_0_ controls the within-population variance (or the covariance when distance between points is 0, known as a “sill” in the geospatial literature), *α*_1_ controls the rate of the decay of covariance per unit pairwise distance, and *α*_2_ determines the shape of that decay. Within-population variance may vary across samples due to either noise from a finite sample size or demographic history unique to that sample (e.g., bottlenecks or endogamy). To accommodate this heterogeneity we introduce population-specific variance terms, resulting in the covariance matrix for standardized sample frequencies

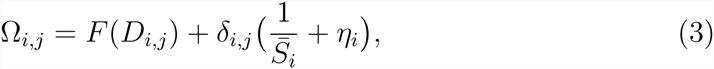

where *δ*_*i,j*_ = 1 if *i* = *j* and is 0 otherwise, *η*_*k*_ is a nonnegative sample-specific variance term (nugget) to account for variance specific to population *k* that is not accounted for by the spatial model, and 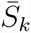 is the mean sample size across all loci in population *k*, so that 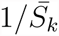 accounts for the variance introduced by sampling within the population.

The distribution of the sample covariance matrix 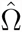 is not known in general, but the central limit theorem implies that if the number of loci is large, it will be close to Wishart. Therefore, we assume that 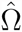is Wishart distributed with degrees of freedom equal to the number of loci (*L*) used and mean equal to the parametric form Ω given in equation (3). We denote this by

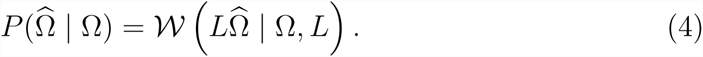

Note that if the standardized sample frequencies are Gaussian, then the sample covariance matrix is a sufficient statistic, so that calculating the likelihood of 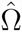 is the same as calculating the probability of the data up to a constant. Handily, it also means that once the sample covariance matrix has been calculated, all other computations do not scale with the number of loci, making the method scalable to genome size datasets. This modeling approach rests on the assumption that the loci in the dataset are independent, that is, not in linkage disequilibrium (LD). Linkage disequilibrium between loci included in the dataset will have the effect of decreasing the true number of degrees of freedom, effectively making this likelihood calculation a composite likelihood and artificially increasing confidence in parameter estimation. We discuss possible ways to accommodate linkage disequilibrium further in the Discussion.

### Location Inference

Non-equilibrium processes like long distance admixture, colonization, or population expansion events will distort the relationship between covariance and distance across the range, as will barriers to dispersal on the landscape. To accommodate these heterogeneous processes we infer the locations of populations on a map that reflects genetic, rather than geographic, proximity. To generate this map, we treat populations’ locations (i.e. coordinates in the geogenetic map) as parameters that we estimate with a Bayesian inference procedure (described in the Methods). These location parameters for each population are denoted by *G*, and determine the matrix of pairwise geogenetic distances between populations, *D*(*G*), which together with the parameters 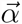 and *η* determine the parametric covariance matrix Ω (given by equation (3)). We acknowledge this dependence by writing 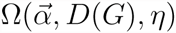.

The prior distributions on the parameters that control the shape and scale of the decay of covariance with distance (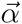 and *η*) are given in the Methods. The priors on the geogenetic locations, *G*, are independent across populations; because the observed locations naturally inform the prior for populations locations, we use a very weak prior on population *k*’s location parameter (*G*_*k*_) that is centered around the observed location. This prior on geogenetic locations also encourages the resulting inferred geogenetic map to be anchored in the observed locations and to represent (informally) the minimum distortion to geographic space necessary to satisfy the constraints placed by genetic similarities of populations. In practice, we also compare results to those produced using random locations as the “observed” locations, and can change the variance on the spatial priors to ascertain the effect of the prior on inference.

We then write the posterior probability of the parameters as

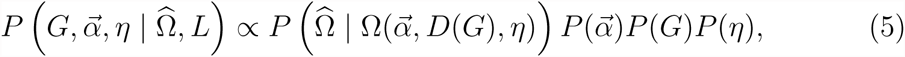

where *P* () denotes the various priors, and the constant of proportionality is the normalization constant.

We then use a Markov chain Monte Carlo algorithm to estimate the posterior distribution on the parameters as described in more detail in the Methods.

**Simulations.** We first apply the method to several scenarios simulated using the coalescent simulator *ms* [Hudson, 2002]. Each scenario is simulated using a stepping stone model in which populations are arranged on a grid with symmetric migration to nearest neighbors (eight neigbors, including diagonals) with 10 haploid individuals sampled from every other population at 10,000 unlinked loci (for details on all simulations, see Methods and Supplementary Materials). The basic scenario is shown in Fig. 1a, which is then embellished in various ways. In the SpaceMix analysis of each simulated dataset, we treat population locations as unknown parameters to be estimated as part of the model, and center the priors on each population’s location at a random point. The resulting geogenetic maps are produced from the parameters having maximum posterior probability. Since overall translation, rotation, and scale are nuisance parameters, we present inferred locations after a Procrustes transformation (best-fit rotation, translation, and dilation) to match the coordinates used to simulate the data. The axes of the resultant maps are presented as Northings and Eastings, as population locations in this geogenetic space no longer conform to the latitude or longitude of the original sampling locations. In Fig. S1, we show the relationship between genetic covariance, geographic distance, and inferred geogenetic distance for these simulations.

**Figure 1:**
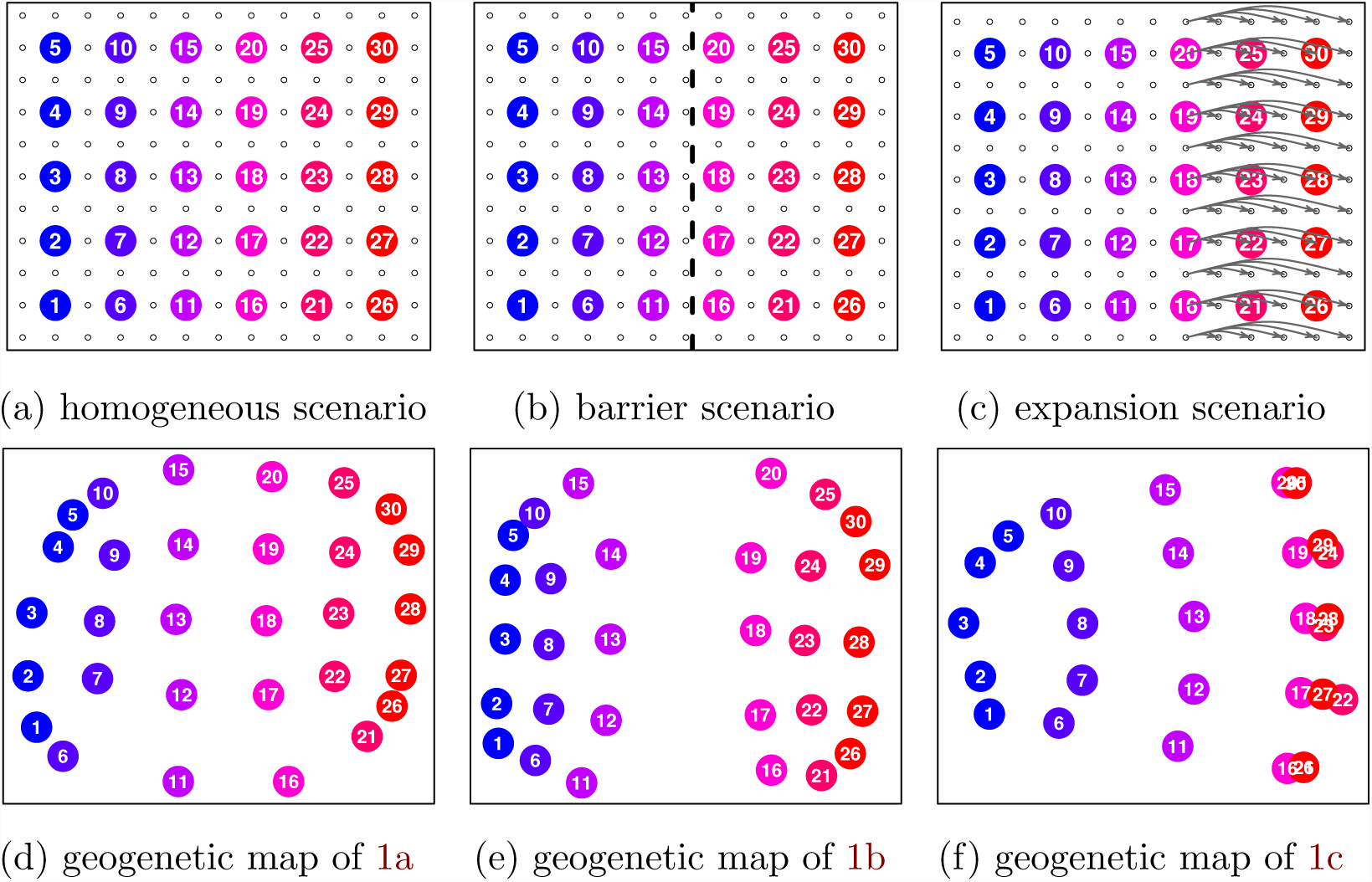
Simulation scenarios and and their corresponding geogenetic maps estimated with SpaceMix. The smaller circles in the simulation scenarios represent unsampled populations. a) configuration of simulated populations on a simple lattice with spatially homogeneous migration rates; b) a lattice with a barrier along the center line of longitude, across which migration rates are reduced by a factor of 5; c) a lattice with recent expansion on the eastern margin; d) the maximum *a posteriori* (MAP) estimate from the posterior distribution of population locations under the scenario in 1a; e) MAP estimate of population locations under the scenario in 1b; f) MAP estimate of population locations under the scenario in 1c;

The lattice scenarios, illustrated in Figs. 1 and 2, are: homogeneous migration rates across the grid; a longitudinal barrier across the center of the grid; a series of recent expansion events; and an admixture event between opposite corners of the lattice. In the simple lattice scenario with homogeneous migration rates (Figs. 1a and 1d), SpaceMix recovers the lattice structure used to simulate the data (i.e., populations correctly find their nearest neighbors). After adding a longitudinal barrier to dispersal across which migration rates are reduced by a factor of 5 (Fig. 1b), the two halves of the map are pushed farther away from one another, reflecting the decreased gene flow between them.

**Figure 2:**
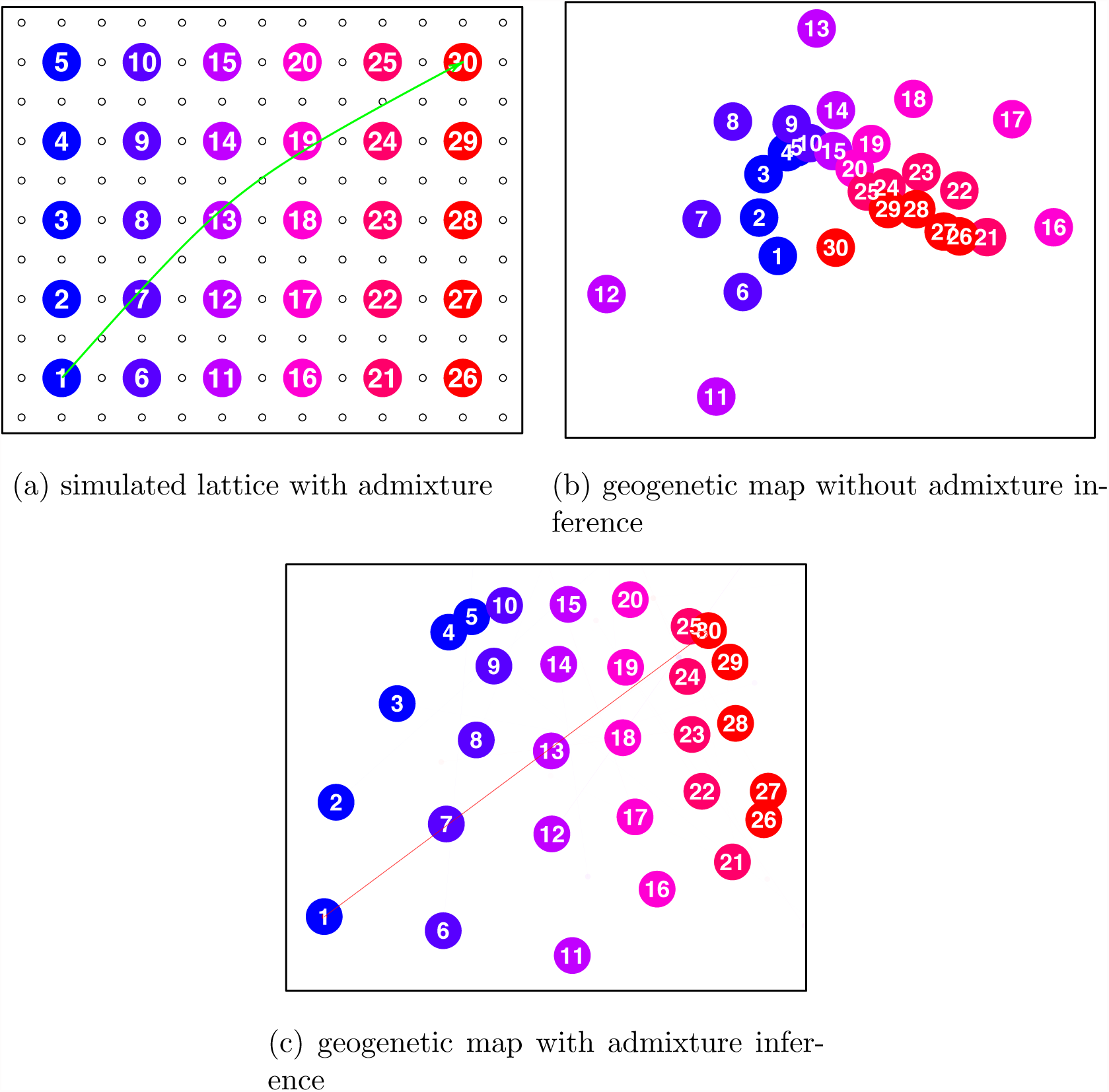
Simulation scenarios and SpaceMix inference. a) a lattice with recent admixture event between population 1 in the southwest corner and population 30 in the northeast corner, so that population 30 is drawing half of its ancestry from population 1; b) the estimate of population locations under this scenario; c) the estimate of population locations and their sources of admixture under this scenario. The 95% credible interval on *w*_30_ is 0.36–0.40. In panel (c), the width and opacity of the admixture arrows are drawn proportional the admixture proportions.

In the expansion scenario, in which all populations in the last five columns of the grid have expanded simultaneously in the immediate past from the nearest population in their row (Fig. 1c), the daughter populations of the expansion event cluster with their parent populations, reflecting the higher relatedness (per unit of geographic separation) between them.

In all scenarios, populations at the corners of the lattice are pulled in somewhat because these have the least amount of data informing their relative placements, and because, without nearest-neighbor migration from farther outside the lattice, they are in fact more closely related to their neighbors. We also examined the effects of uneven sampling on inference by sampling random subsets of the simple lattice and barrier scenarios shown in Figs. 1a and 1b. The results of these analyses are given in Figs. S3a and S3b.

We next simulated a long-distance admixture event on the same grid, by sampling half of the alleles of each individual in the northeast corner population from the southwest corner population (Fig. 2a). We then ran a SpaceMix analysis in which the locations of these populations were estimated (Fig. 2b). The admixture creates excess covariance over anomalously long distances, which is clearly difficult to accommodate with a two-dimensional geogenetic map. Fig. 2b shows the torturous lengths to which the method goes to fit a good geogenetic map: the admixed population 30 is between population 1, the source of its admixture, and populations 24, 25, and 29, the nearest neighbors to the location of its non-admixed portion. However, this warping of space is difficult to interpret, and would be even more so in empirical data for which a researcher does not know the true demographic history.

### Inference of Spatial Admixture

To incorporate recent admixture, we allow each allele sampled in population *k* to have a probability *w*_*k*_ (0 *≤ w*_*k*_ *≤* 0.5) of being sampled from location 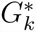,which we refer to as population *k*’s source of admixture, and a probability 1 - *w*_*k*_ of being sampled from location *G*_*k*_. With no nugget, each allele would be sampled independently, but the nugget introduces correlations between the alleles sampled in each population.

With this addition, the parametric covariance matrix before given by (3) becomes a function of all the pairwise spatial covariances between the locations of populations *i* and *j* and the points from which they draw admixture (illustrated in Fig. 3); now, we model the covariance between 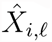 and 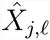 for each *l*, as

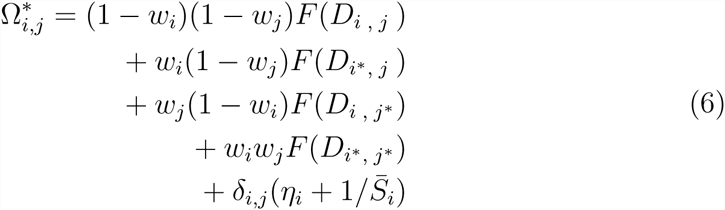

where *D* is the 2*k* × 2*k* matrix of pairwise distances between all inferred locations and sources of admixture, and for readability, we denote, e.g., *F* (*D*(*G*_*i*_, *G*^*^_*j*_)), as *F* (*D*_*i,j*^*^_). The spatial covariance, *F* (*D*), is as given in equation (2), and we reintroduce the nugget, *η*_*k*_, and the sample size effect, 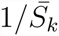 for each population as above in Eqn. (3).

**Figure 3:**
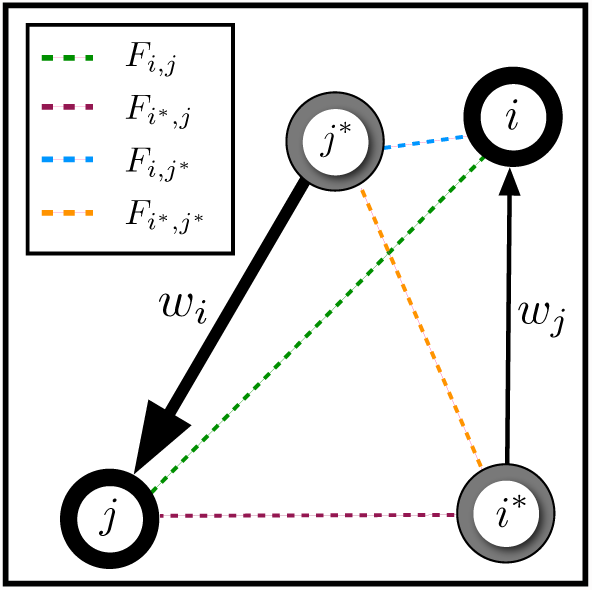
An illustration of the form of the admixed covariance given in Equation (6). Populations *i* and *j* are drawing admixture in proportions *w*_*i*_ and *w*_*j*_ from their respective sources of admixture, *i*^*^ and *j*^*^, and all pairwise spatial covariances (the *F* ’s) are shown. In this cartoon example, population *j* is drawing more admixture from its source *j*^*^ than *i* is from its source *i*^*^ (i.e., *w*_*j*_ *> w*_*i*_).

We proceed in our inference procedure as before, but now with the locations of the sources of admixture and the admixture proportions to infer. The likelihood of the sample covariance matrix is exactly as before in (4), except with Ω replaced by Ω^*^. The posterior probability of these parameters can be expressed as a function of this parametric admixed covariance, Ω^*^,

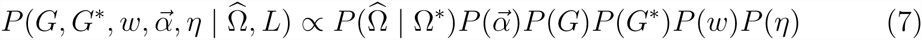

as specified by the parameters *w*, *G*^*^, 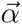, and *η*, and the inferred locations, *G*. We place a weak spatial prior on the sources of admixture, *G*^*^ around the centroid of the observed locations. The admixture proportions, *w*, are capped at 0.5, to ensure identifiability, and are heavily weighted towards small values to be conservative with respect to admixture inference. These priors are detailed in Table 2.

The models described above may be used in various combinations. In the simplest model, locations are not estimated for populations, nor do they draw admixture; the only parameters to be estimated are those of the spatial covariance function given in equation (2), and the population-specific variance terms (*η*_*i*_). In the most complex model, population locations, the locations of their sources of admixture, and the proportions of admixture are all estimated jointly in addition to the parameters of the spatial covariance function and the population specific variances. Users may wish to employ the more constrained models (e.g., fixing the locations or admixture proportions for some or all samples) in a model selection framework to test specific hypotheses. We discuss the utility of these different models in the Appendix.

Allowing admixture gives sensible results for the scenario of Fig. 2a; in the resulting map, the only population that draws substantial admixture is the one that is actually admixed, and it draws admixture (95% CI: 0.36 - 0.40) from the correct location (Fig. 2c).

A more subtle simulated admixture scenario, with admixture proportion of 10% across a geographic barrier, is shown Fig. 4a. The resulting SpaceMix map (Fig. 4b), separates the east and west sides of the grid to accommodate the effect of the barrier, and the admixed population (population 23) draws admixture from very close to its true source (population 13), and in close to the correct amount 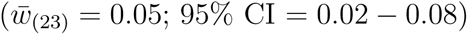.

**Figure 4:**
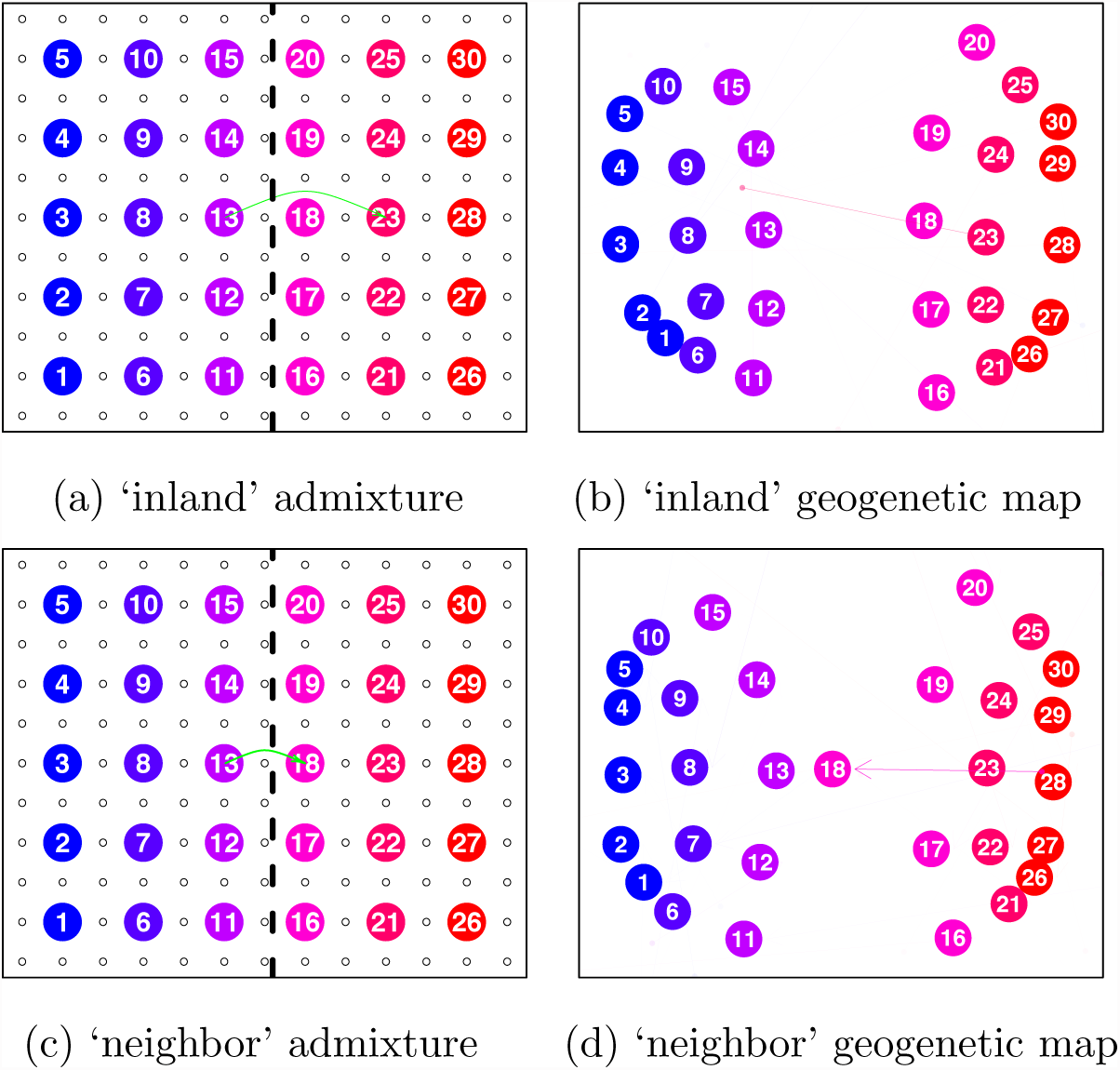
Simulation scenarios and inferred population maps for two different admixture scenarios: a) lattice with a barrier and an admixture event (10%) across the barrier to an ‘inland’ population; b) the inferred population map for the scenario in (a), where the admixed population 23 is the only population drawing non-negligible admixture (95% CI: 0.02-0.08); c) lattice with a barrier and an admixture event (40%) across the barrier to a ‘neighbor’ population on the border of the barrier; (d) the inferred population map for the scenario in (c), where the admixed population 18 is the only population drawing non-negligible admixture (95% CI: 0.04–0.14).

Another difficult scenario is shown in Fig. 4c, where 40% admixture has occurred between two populations immediately adjacent to each other on either side of a barrier. Here, the admixed population 18 is correctly identified as admixed; however, its intermediate genetic relationships are explained through an estimated location close to its true admixture source (population 13) and source of admixture (95% CI: 0.04–0.14) on the far margin of the half of the grid on its own side of the barrier. Because there is no sampled intervening population between admixed population 18 and its source of admixture 13, the model is able to explain population 18’s its higher covariance with population 13 via its estimated location *G*_(18)_, rather than via that of its source of admixture *G*^*^_(18)_. In each of these scenarios, the estimated admixture proportion is less than that used to simulate the data. This is due to the stringent prior we place against admixture. We discuss these examples further in the Methods.

## Empirical Applications

To demonstrate the applications of this novel method, we analyzed population genomic data from two systems: the greenish warbler ring species complex, and a global sampling of contemporary human populations. Maps showing our sampling in these two systems are given in Fig. 5, and information on the specific samples included is given in the Supplementary Materials, Tables S1 and S2. For all analyses presented below, we centered the priors on location parameters at randomly chosen locations rather than at the observed geographical locations. Each geogenetic map shown here is the maximum a posteriori estimate (over all parameters), transformed by rotation, translation, and scaling to best fit inferred locations (*G*) to the observed latitude and longitudes (a full Procrustes transformation). As with the simulations described above, the axes of the geogenetic maps are presented in Eastings and Northings.

**Figure 5:**
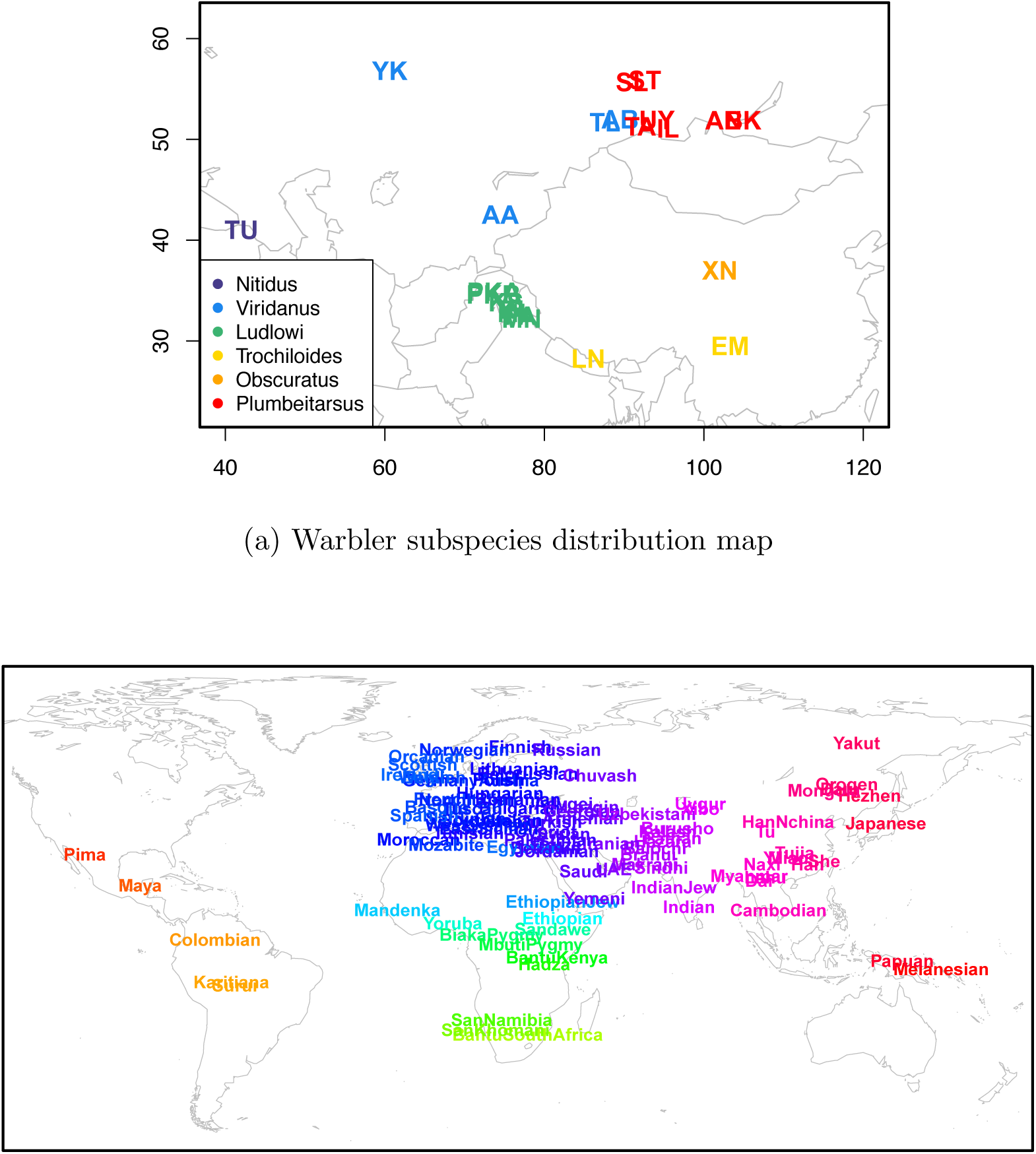
Sampling maps of both empirical systems analyzed. (a) greenish warbler subspecies distributions of all 22 sampled populations (breeding grounds), consisting of 95 individuals and colored by subspecies; (b) sampling map for human dataset, consisting of 1,490 individuals from 95 population samples.

### Greenish Warblers

The greenish warbler (*Phylloscopus trochiloides*) species complex is broadly distributed in their breeding habitat around the Tibetan plateau, and exhibits gradients around the ring in a range of phenotypes including song, as well as in allele frequencies [Ticehurst, 1938, Irwin et al., 2001, 2005]. At the northern end of the ring in central Siberia, where the eastern and western arms of population expansion meet, there are discontinuities in call and morphology, as well as reproductive isolation and a genetic discontinuity [Irwin et al., 2001, 2008]. It is proposed that the species complex represents a ring species, in which selection and/or drift, acting in the populations as they spread northward on either side of the Tibetan plateau, have led to the evolution of reproductive isolation between the terminal forms.

The question of whether it fits the most strict definition of a ring species focuses on whether gene flow around the plateau has truly been continuous throughout the history of the expansion or if, alternatively, discontinuities in migration around the species complex’s range have facilitated periods of differentiation in genotype or phenotype without gene flow [Mayr, 1942, 1970, Coyne, 2004] (see Wake and Schneider [1998] for discussion). Alcaide et al. [2014] have suggested that the greenish warbler species complex constitutes a ‘broken’ ring species, in which historical discontinuities in gene flow have facilitated the evolution of reproductive isolation between adjacent forms.

To investigate this question, we applied SpaceMix to the dataset from Alcaide et al. [2014], consisting of 95 individuals sampled at 22 distinct locations and sequenced at 2,334 SNPs, of which 2,247 were bi-allelic and retained for SpaceMix runs. These loci were treated as independent (i.e., un-linked). We discuss other ways to accommodate linkage disequilibrium further in the Discussion.

We first ran SpaceMix on the population dataset, with no admixture. The resulting inferred map (Fig. 6a) largely recapitulates the geography of the sampled populations around the ring. The Turkish population (**TU**, *Phylloscopus trochiloides* ssp. *nitidus*) clusters with the populations in the subspecies *ludlowi*, due to its recent expansion, but also has a relatively high nugget parameter (see Fig. S4a), reflecting the population history it does not share with its *ludlowi* neighbors. In the north, where the twin waves of expansion around the Tibetan Plateau are hypothesized to meet, the inferred geogenetic distance between populations from opposite sides of the ring was much greater than their observed geographic separation, reflecting the reproductive isolation between these adjacent forms (see Fig. S5).

**Figure 6:**
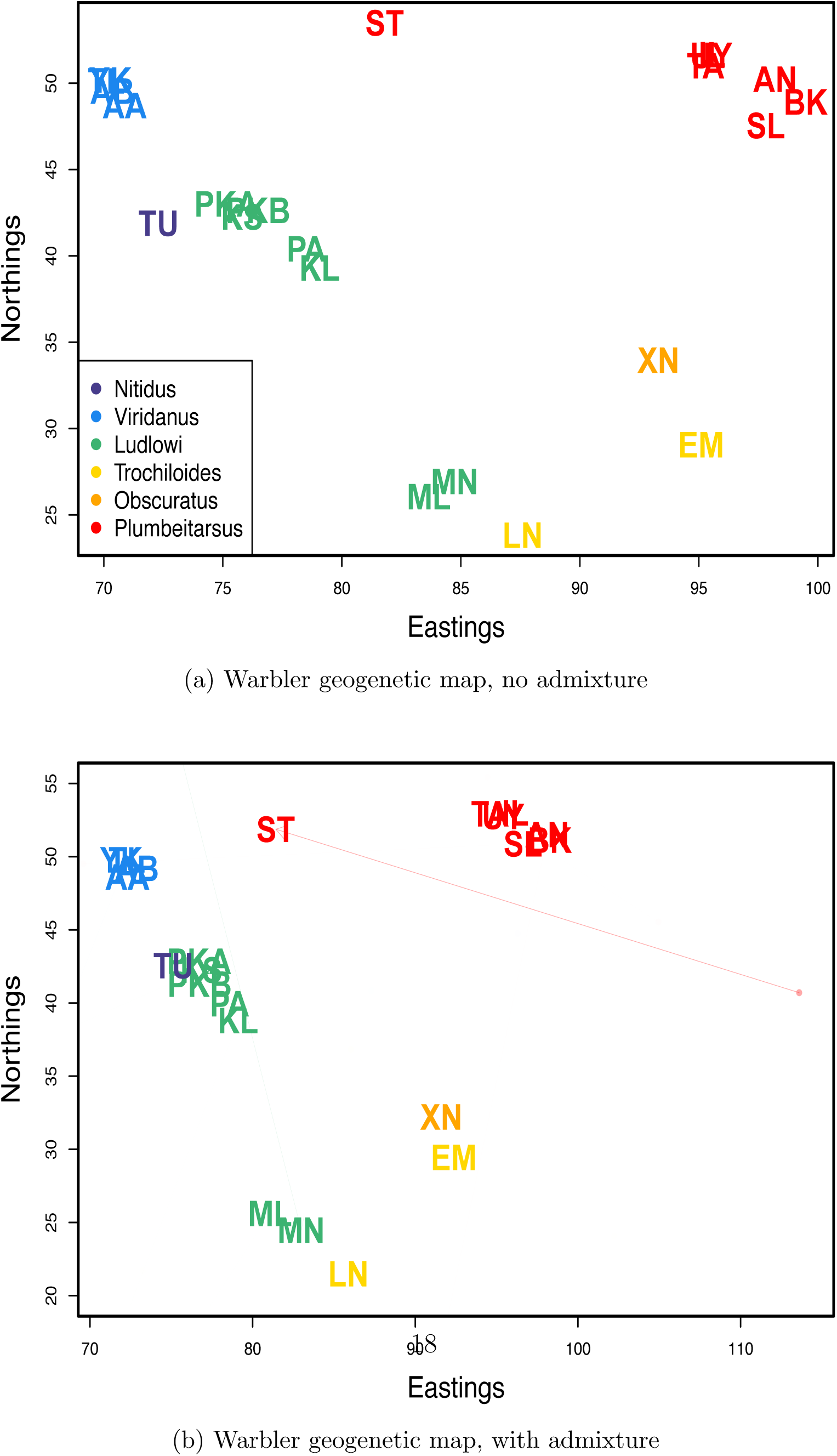
Inferred population maps with population labels colored as in Fig. 5a: a) the map inferred with no admixture inference; b) the map inferred with admixture inference.

We then ran the method allowing admixture (Fig. 6b). The only population sample with appreciable admixture is the Stolby sample (**ST**; *w* = 0.19, 95% credible interval: 0.146-0.238; Fig. S6). This sample is known to be composed of an equal mixture of eastern *plumbeitarsus* and western *viridanus* individuals [Alcaide et al., 2014]. Multiple runs agreed well on the level of admixture of the Stolby sample (see Fig. S7). What does vary across runs is whether the Stolby sample has an estimated location by the *viridanus* cluster while drawing admixture from near the *plumbeitarsus* cluster, or vise versa; however, this is to be expected given the 50/50 nature of the sample’s makeup (Fig. S7). The somewhat intermediate position of the Stolby sample, and its non-50/50 admixture proportion, likely partially reflect the influence of the priors (Fig. S8).

We repeated these analyses (with and without admixture) on an individual level (Fig. 7). No individual drew appreciable admixture (see Fig. S9 for admixture proportions), and so we discuss the results with admixture (those without admixture are nearly identical, see Figs. S10, S11, and S12). As with the analysis on multi-sample populations, the results approximately mirror the geography of the individuals.

**Figure 7:**
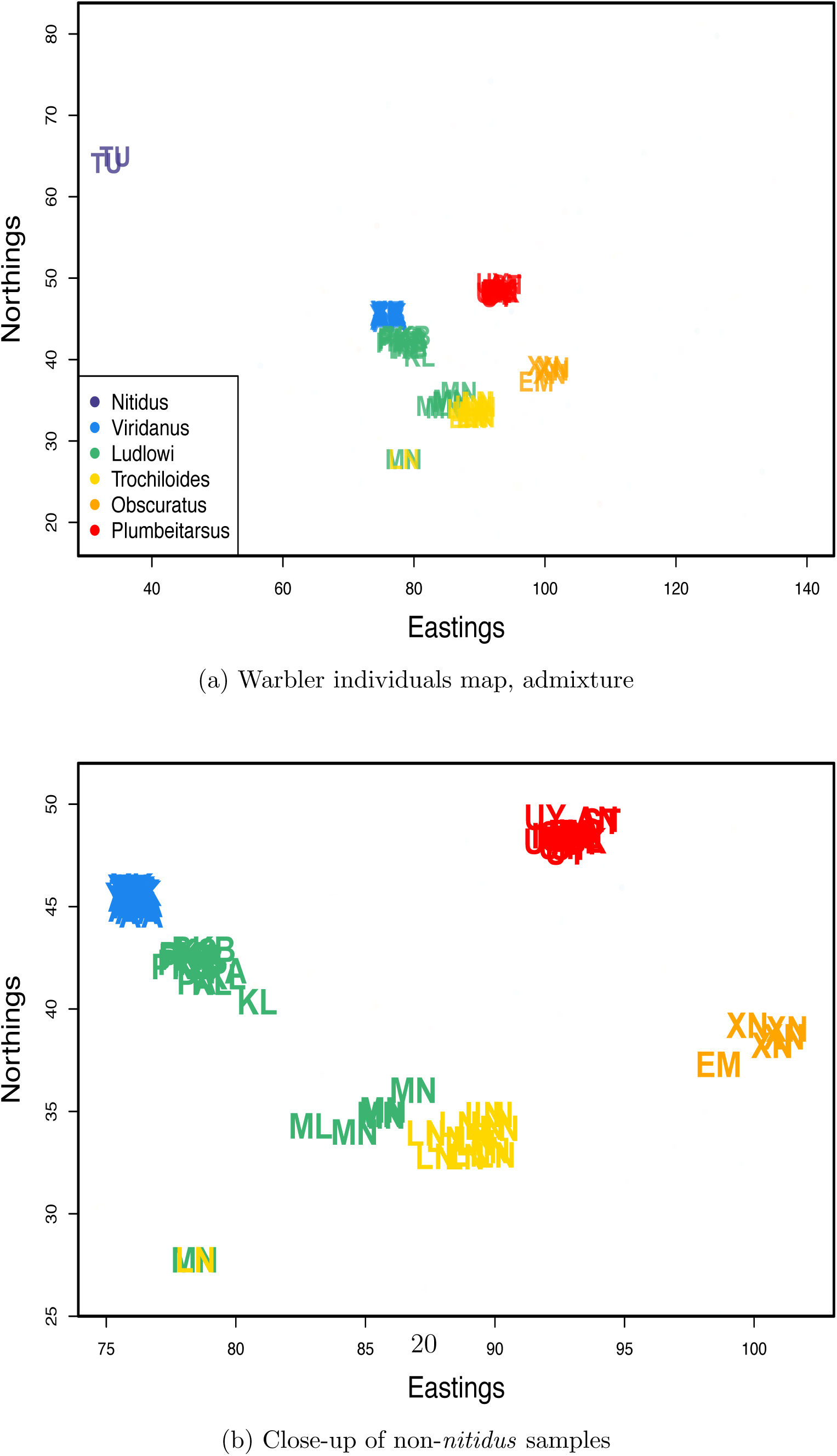
Inferred maps for warbler individuals, colored by subspecies in an analysis with admixture inference. a) map inferred with admixture; b) close-up of all non-*nitidus* samples in the admixture map.

There are, however, a number of obvious departures in the individual geogenetic map from the population map. The most obvious is that the location of a pair of *nitidus* samples (in purple) is very far from the rest of the samples. These individuals appear to be fairly close relatives: in the population-level analysis of Fig. S4a, this increase in shared ancestry was accounted for by a large nugget for the *nitidus* population. However, in the individual-level analysis, a nugget is estimated separately for each sample, so, the model must accommodate the much higher relatedness between this pair of individuals through estimated locations that are close to each other and far from the rest of the samples. The same phenomenon seems to be at work in determining the locations of a pair of individuals, one identified as *P. t. ludlowi* (Lud-MN3), one as *P. t. trochiloides* (Tro-LN11), as they also show an unusually low pairwise sequence divergence (see Fig. S13).

The split between *viridanus* and *plumbeitarsus* individuals (blue and red, respectively), in the north at the contact zone of the two waves of expansion, is clearer now than in the population-based analysis, as the estimated locations of individuals from the Stolby population are near their respective clusters. Although the geogenetic separation between the *viridanus* and *plumbeitarsus* individuals is greater than their geographic separation, they are still closer to each other than we would expect if all gene flow between the two was mediated by the southern populations, in which case we would expect the populations to form a line, with *viridanus* at one end and *plumbeitarsus* at the other. This horseshoe configuration, with *viridanus* and *plumbeitarsus* at its tips, is steady within and among runs of the MCMC and choice of position priors (see Figs. S11a-S11c).

Is this biologically meaningful? A similar horseshoe shape appears when a principal components (PC) analysis is conducted and individuals are plotted on the first two PCs (see Fig. S14 and Alcaide et al. [2014]). However, as discussed by Novembre and Stephens [2008], such patterns in PC analysis can arise for somewhat unintuitive reasons. If populations are simulated under a one dimensional stepping stone model, then plotting individuals on the first two PCs results in a horseshoe (e.g. see Fig. S15b) not because of gene flow connecting the tips, but rather because of the orthogonality requirement of PCs [see Novembre and Stephens, 2008, for more discussion]. In contrast, when SpaceMix is applied to data simulated on a one dimensional array of populations, the placement of samples is consistent with a line (see Figs. S15c, S15d). The proximity of *viridanus* and *plumbeitarsus* in geogenetic space may be due to gene flow between the tips of the horseshoe north of the Tibetan Plateau. This conclusion is in agreement with that of Alcaide et al. [2014], who observed evidence of hybridization between *viridanus* and *plumbeitarsus* using assignment methods.

The SpaceMix map also diverges from the observed map in the distribution of individuals from the subspecies *ludlowi* (in green). These samples were taken from seven sampling locations along the southwest margin of the Tibetan Plateau, but, in the SpaceMix analysis, they partition into two main clusters, one near the *trochiloides* cluster, and one near the *viridanus* cluster. This break between samples from the same subspecies, which is concordant with the findings of Alcaide et al. [2014], makes the *ludlowi* cluster unusual compared to the estimated spatial distributions of the other subspecies (see Fig. S16), and suggests a break in historic or current gene flow.

### Human Populations

Human population structure is a complex product of the forces of migration and drift acting on both local and global scales, patterned by geography [Novembre et al., 2008, Ralph and Coop, 2013], time [Skoglund et al., 2012, 2014], admixture [Hellenthal et al., 2014], landscape and environment [Beall et al., 2010, Bigham et al., 2010, Bradburd et al., 2013], and shaped by culture [Reich et al., 2009, Atzmon et al., 2010, Moorjani et al., 2011]. To visualize the patterns these processes have induced, we create a geogenetic map for a worldwide sample of modern human populations. Of course, human history at these geographic scales has many aspects that are not well captured by static maps with discrete “arrows” of admixture. Nonetheless, we talk about the locations of samples and their sources of admixture as if these are fixed, even though both reflect the compounding of drift and gene flow over many historical processes. We therefore urge caution in the interpretation our results, and view them as a simplistic but rich visualization of patterns of population structure.

We used the dataset of Hellenthal et al. [2014], comprised of 1,490 individuals from 95 population samples (see Fig. 5b for map of sampling), as well as the latitude and longitude attributed to each sample. In the analyses presented on human genotype data below, we have thinned the total dataset for LD in windows of 50 base-pairs, with a step-size of 5 base-pairs, and an upper limit of 0.2 on pairwise *r*^2^ [Purcell et al., 2007, Purcell, 2009]. We then used a random subset of 10,000 SNPs to estimate the sample covariance.

We ran two sets of SpaceMix analyses: in the first, we estimated population sample locations, and in the second, we also allowed admixture. We note that few of the putative admixture events that we report have escaped the notice of previous investigators, which is unsurprising given the depth of recent attention on human admixture studies, particularly on the subset subset of these samples that are in the HGDP dataset [see Rosenberg et al., 2002, Li et al., 2008, Loh et al., 2013, Patterson et al., 2012, Hellenthal et al., 2014, for various global analyses]. Below, when discussing a pattern we see in our analyses, we often cite other authors who have seen or suggested similar patterns. However, what is novel here is the ability to visualize these admixture events in a geographic context, and that these admixture signals stand out against a null model of migration in continuous space (rather than tree-based models).

When we only infer the location of each sample, the map roughly recapitulates the geography of the samples (Fig. 8a), a result that holds nicely when we zoom in on the more heavily sampled area of Eurasia (Fig. 8b). We see that samples both in the Americas and in Oceania lie close to the East Asian samples, but that they form two clusters on opposite sides. The proximity of these groups to the East Asians represents the fact that both groups share an ancestral population in the relatively recent past with East Eurasian populations, but the two expansions occurred independently. As in our simulations (Fig. 1f) population expansions/bottlenecks have distorted the relationship between geographic and geogenetic distance. Geogenetic distances between samples within Africa are much greater than those between any other group (see Fig. S17), and the slope of the relationship between geographic and geogenetic distances between populations on each continent decays with distance from Africa. This pattern is consistent with a history of human colonization events characterized by serial bottlenecks [Harpending and Rogers, 2000, Prugnolle et al., 2005, Ramachandran et al., 2005] following an out-of-Africa expansion, and subsequent expansions into Western Eurasia, East Asia, the Americas, and Oceania [but see Pickrell and Reich, 2014, for a discussion of other models].

**Figure 8:**
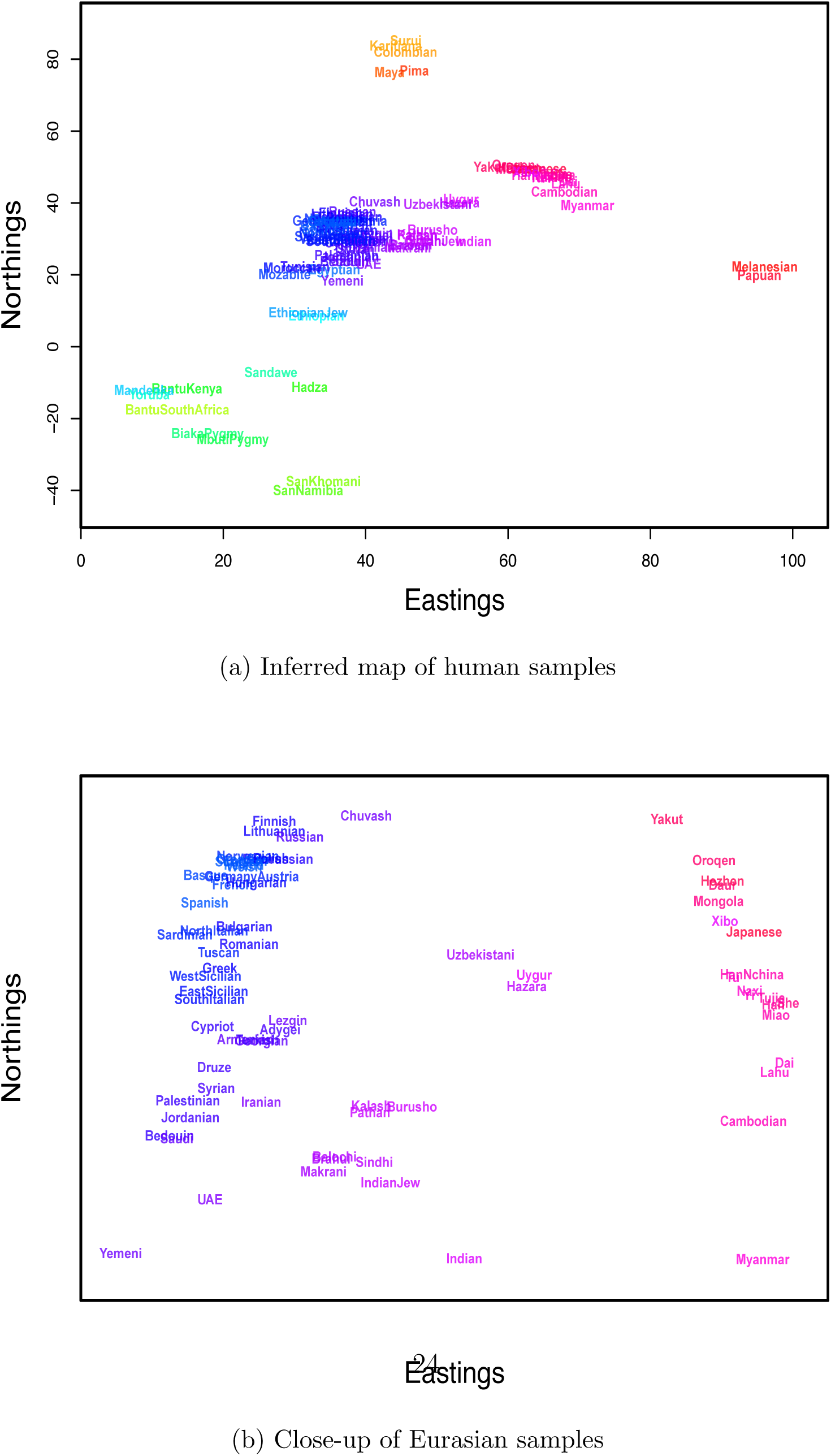
Map of human samples, inferred without admixture. (a) complete map; close-up of Eurasian samples.

To investigate possible patterns of admixture further, we ran a SpaceMix analysis with admixture (results shown in Figs. 9 and 10). The biggest change between the geogenetic map of human populations inferred with admixture and that without is the positioning of African samples with respect to the rest of the world. The relatively large geogenetic distances between these groups reflects the fact that Eurasian, North African, Oceanian, and American populations all share relatively large amounts of population history (and hence genetic drift) not shared with the Sub-Saharan African samples. Relative to the geogenetic map inferred without admixture, the inclusion of admixture shifts the estimated locations of admixed samples intermediate between Sub-Saharan Africa and North Africa/the Middle East toward one cluster or the other, which, in turn, pushes each of those major clusters to move relatively farther apart. The Ethiopian and Ethiopian Jewish samples have estimated locations closer to the Sub-Saharan samples than the of the North African samples, but draw substantial amounts of admixture (∼ 40%) from close to where the Egyptian sample has positioned itself in the the Middle East cluster, as do the Sandawe [Hodgson et al., 2014, Pickrell et al., 2012]. The SanKhomani draw admixture from near Syria, which may reflect multiple distinct geographic sources of admixture as discussed by Hellenthal et al. [2014] and Pickrell et al. [2014]. Interestingly the Bantu South African sample, though it has an estimated location near the other Bantu samples, draws admixture from close to the San populations. This is consistent with previous signals of the expansion of Bantu-speaking peoples into southern Africa [Pickrell et al., 2012, Schlebusch et al., 2012, Pickrell et al., 2014, Hellenthal et al., 2014]. The inferred sample-specific drift parameters (the ‘nuggets’) are similar between runs with and without admixture (Fig. S18).

**Figure 9:**
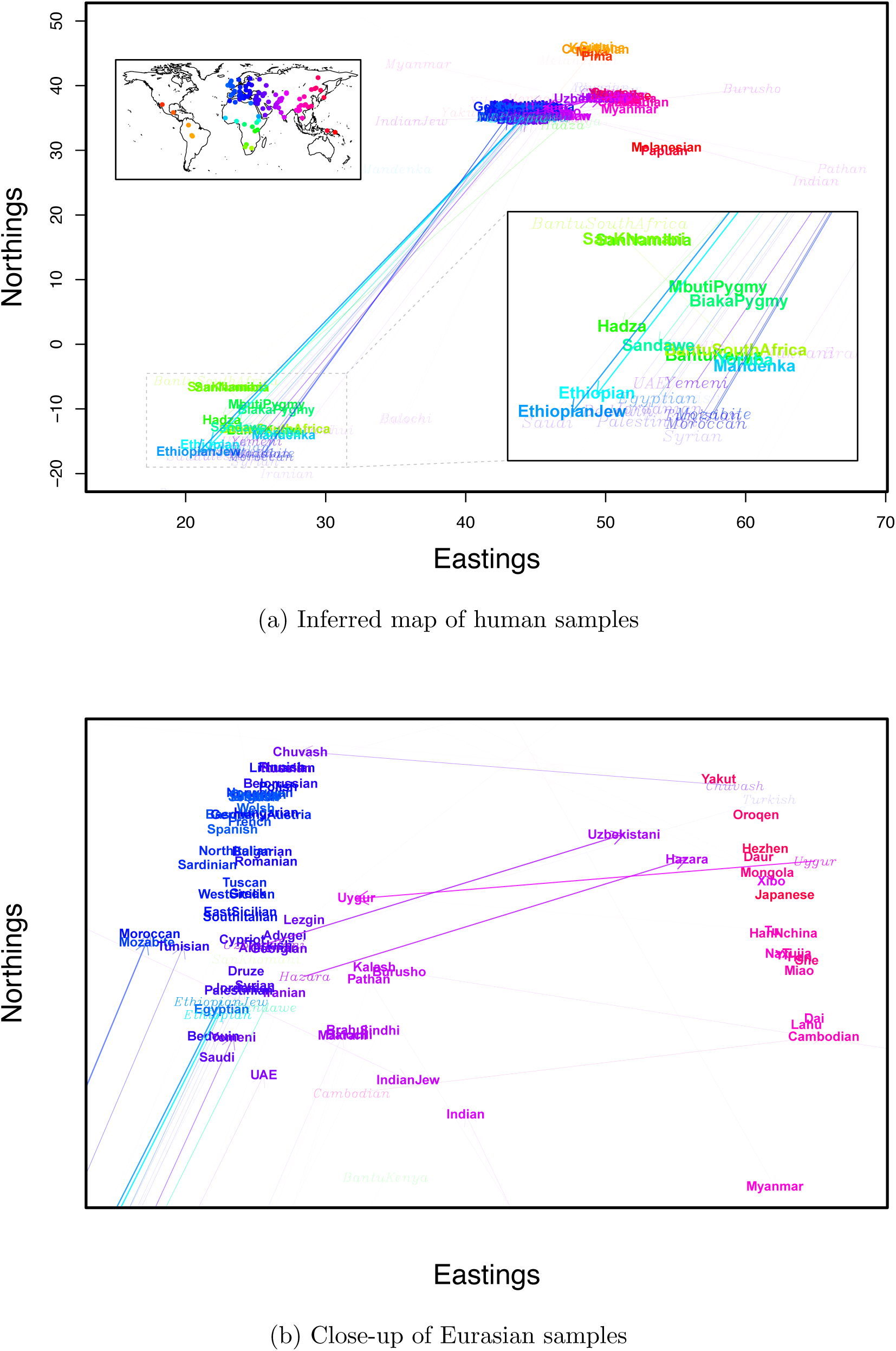
Map of human samples, inferred with admixture. (a) complete map; close-up of Eurasian samples. Italicized labels denote locations of admixture sources, with darkness proportional to the amount of admixture.

**Figure 10:**
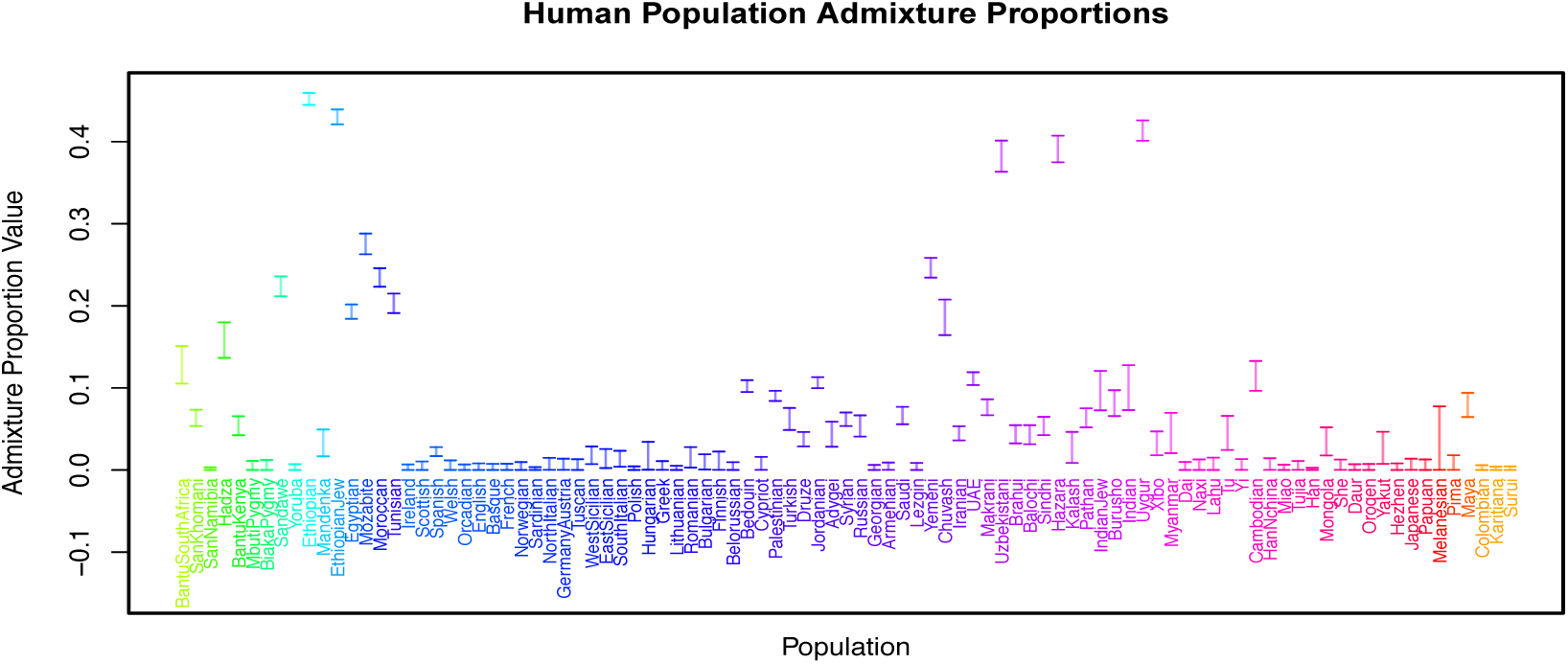
Mean admixture proportions (and 95% CIs) for each population sample.

The majority of North African samples (Egyptian, Tunisian, Morocan, Mozabite) join the Middle Eastern samples (positioned in rough accord with their sampling location along North Africa), and draw admixture from near the Ethiopian samples. All of the Middle Eastern samples draw admixture from close to the geogenetic location of the Ethiopian samples and where most of the North African samples draw admixture from, representing the complex history of North African–Middle Eastern gene flow [Henn et al., 2012, Hellenthal et al., 2014].

A number of other population samples draw admixture from Africa. The Sindhi, Makrani, and Brahui draw admixture from close to the location of the Bantu samples [Hellenthal et al., 2014], and the Balochi and Kalash draw admixture from some distance away from African population samples. Of the European samples, the Spanish and both East and West Sicilian samples draw small amounts of admixture from close to the Ethiopian samples, presumably reflecting a North African ancestry component [Moorjani et al., 2011, Botigu et al., 2013].

The other significant signal of admixture is between East and West Eurasia, a signal documented by many authors [Rosenberg et al., 2002, Li et al., 2008, Xu and Jin, 2008, Hellenthal et al., 2014]. The majority of samples maintain their relative positions within each of these groups; however, there are several samples that show admixture between eastern and western Eurasia. The Uzbekistani and Hazara samples have estimated locations close to the East Asian samples and draw a substantial admixture proportion from close to where the Georgian and Armenian samples have located themselves. Conversely, the Uygur sample has an estimated location close to the Burusho, Kalash, and Pathan samples, and draws admixture from near the Mongola and Hezhen samples. The Tu sample (with a geogenetic location in East Asia) draws a small amount of ancestry from close to the estimated location of the Uygur. The estimated location of the Chuvash sample is near the Russian and Lithuanian samples, and the Chuvash draw admixture from close to the Yakut (as do the Turkish, to a smaller extent). There are several other East-West connections: the Russian and Adygei samples have admixture from a location “north” of the East Asian samples, and the Cambodia sample draws admixture from close to the Eygptian sample [Pickrell and Pritchard, 2012, Hellenthal et al., 2014].

There are also a number of samples that draw admixture from locations that are not immediately interpretable. For example, the Hadza and Bantu Kenyan samples draw admixture from somewhat close to India, and the Xibo and Yakut from close to “northwest” of Europe. The Pathan samples draw admixture from a location far from any other samples’ locations, but close to where the India samples also draws admixture from. The Myanmar and the Burusho samples both draw admixture far from the locations estimated for other samples as well.

There are a number of possible explanations for these results. As we only allow a single admixture arrow for each sample, populations with multiple, geographically distinct sources of admixture may be have estimated admixture locations that average over those sources. This may be the case for the Hadza and Bantu Keynan samples [Hellenthal et al., 2014]. A second possibility is that the relatively steep prior on admixture proportion forces samples to draw lower proportions of admixture from locations that overshoot their true sources; this may explain the Xibo and Yakut admixture locations. A final explanation is that good proxies for the sources of admixture may not be included in our sampling, either because of of the limited geographic sampling of current day populations, or because of old admixture events from populations from which there are not other more directly descending modern populations. The admixture into the Indian and Pathan samples (whose admixture source also clusters with the Indian Jew samples in some MCMC runs) may be an example of this; Reich et al. [2009] and Moorjani et al. [2013] have hypothesized that many populations from the Indian subcontinent may be descended from an admixture event involving an ancestral Southern Indian population not otherwise represented in this dataset.

In Figs. S19 and S20, we show the results of other independent MCMC analyses on these data. The broad-scale patterns and results discussed above are consistent across these runs. However, as is to be expected, there is significant heterogeneity in the exact layout of sample and admixture locations. For example, there is some play, among MCMC runs, in the internal orientation of the African locations with respect to the east-west axis within the Eurasian cluster. For some samples that draw a significant amount of admixture, such as the central Asian populations (Uygur, Hazara and Uzbekistani), the estimated location switches with that of their source of admixture (as was also seen across MCMC runs in the warbler data analysis). Similarly the Ethiopian and Ethiopian Jew samples have estimated locations, in some MCMC runs, close to the other North African samples, and draw admixture from near the Sub-Saharan samples (as do the other North African samples).

### Discussion

In this paper we have presented a statistical framework for modeling the geography of population structure from genomic sequencing data. We have demonstrated that the method, SpaceMix, is able to accurately present patterns of population structure in a variety of simulated scenarios, which included the effects of spatially heterogeneous migration, population expansion, and population admixture. In empirical applications of SpaceMix, we have largely recovered previously estimated population relationships in a circum-Tibetan sample of greenish warblers and in a global sample of human populations, while also providing a novel way to depict these relationships. The geogenetic maps SpaceMix generates serve as simple, intuitive, and information-rich summaries of patterns of population structure. SpaceMix combines the advantages of other methods for inferring and illustrating patterns of population structure, using model-based inference to infer population relationships (like TreeMix [Pickrell and Pritchard, 2012], and MixMapper [Lipson et al., 2013]), and producing powerful visualizations of genetic structure on a map (like PCA [Patterson et al., 2006] and SPA [Yang et al., 2014]).

The patterns of genetic variation observed in modern populations are the product of a complex history of demographic processes. We choose to model those patterns as the outcome of a spatial process with geographically determined migration, and we have included statistical elements to accommodate deviations from spatial expectations. However, the true history of a sample of real individuals is vastly more complex than any low-dimensional summary, and, as with any summary of population genetic data, SpaceMix results should be interpreted with this in mind. Furthermore, our “admixture” events are shorthands for demographic relationships that occurred over possibly substantial lengths of time and regions of the globe; approximating this by a single arrow between two points on a map is certainly an oversimplification. Aspects of population history that are better described as a population phylogeny may be difficult to interpret using SpaceMix, and may be better suited to visualization with hierarchical clustering-based methods [Pritchard et al., 2000] or TreeMix/MixMapper-like methods [Pickrell and Pritchard, 2012, Lipson et al., 2013]. There is obviously no one best approach to studying and visualizing population structure; investigators should employ a range of appropriate methods to identify those that provide useful insight.

SpaceMix offers much of the flexibility of PCA – like PCA, it is well suited to describing population structure in a continuous fashion – but it also has a number of advantages over PCA. When isolation by distance holds, the first (one or) two PCs often correspond reasonably well to some simple rotation of latitude and longitude; however, these first two PCs explain a relatively small part of the total variance of the data. Furthermore, because PCs are linear functions of the genotypes, sometimes many PCs must be used to depict patterns produced by simple isolation by distance [Novembre and Stephens, 2008]. These higher order PCs can be hard to interpret in empirical data (see discussion in the warbler section). The recently introduced SPA approach [Yang et al., 2012], which also assumes allele frequencies are monotonically increasing in a given direction, may suffer from the same problem (although we note that PCA and SPA both have significant speed advantages over SpaceMix).

In comparison, if isolation by distance holds, then the two dimensions in which SpaceMix infers geogenetic positions for the samples will suffice to capture the geographic patterns of genetic differentiation (to the extent to which the parametric form of the covariance is flexible enough to capture the empirical decay of covariance with distance). The application of SpaceMix to humans nicely illustrates the utility of our approach: the first two PCs of this dataset resemble a boomerang (Fig. S21), with its arms corresponding to the Africa/Non-Africa split and the spread of populations across Eurasia. In contrast, while the SpaceMix geogenetic map is dominated by the genetic drift induced by migration out of Africa, it also captures much more detail than is contained in the first two PCs (e.g., Fig. 9b). This comparison is also nicely illustrated by the example in Fig. S15.

An advantage of PCA is that it can explain more complex patterns of population structure by allowing up to *K* different axes. Although SpaceMix can easily be extended to more than two dimensions, simply by allowing *G*_*i*_ to describe the location of a sample in *d* dimensions, the interpretation and visualization of these higher dimensions would prove difficult, and so for the moment we stick with two dimensions. On the other hand, SpaceMix can describe in two dimensions patterns that PCA, due to the constraints of linearity, would need more to describe. Our method shows the utility of representing both isolation by distance and longdistance admixture on a 2-D geogenetic map. While we perform inference of this map using likelihood-based inference relating a parametric covariance matrix to the observed empirical matrix, it would be interesting to explore other methods of generating this geogenetic map (e.g., Bookstein [1989], Sampson and Guttorp [1992], Yang et al. [2012], Petkova et al. [2014]). Such methods may offer computational speedups and also potentially help place SpaceMix within a broader statistical framework.

Another strong advantage of SpaceMix over current methods is the introduction of admixture arrows. Although PCA can be interpreted in light of simple admixture events [McVean, 2009], and recent methods can locate the recent, spatially admixed ancestry of out-of-sample individuals [Yang et al., 2012, 2014], neither approach explicitly models admixture between multiple geographically distant locations, as SpaceMix does. Assignment methods are designed to deal with many admixed samples [Pritchard et al., 2000], but they have no null spatial model for testing admixture. We feel that an isolation by distance null model is often more appropriate for testing for admixture, especially when there is geographically dense sampling. SpaceMix offers a useful tool to understand and visualize spatial patterns of genetic relatedness when many samples are admixed.

As currently implemented, SpaceMix allows each population to have only a single source of admixture, but some modern populations draw substantial proportions of their ancestry from more than two geographically distant regions. In such cases the inferred source of admixture may fall between the true locations of the parental populations. Although it is statistically and computationally feasible to allow each population to choose more than one source of admixture, we were concerned about both the identifiability and the interpretability of such a model, and have not implemented it. However, there may be empirical datasets in which such a modeling scheme is required to effectively map patterns of population structure. In addition, we have assumed that only single populations are admixed, when in fact it is likely that particular admixture events may affect multiple samples.

One concern is that the multiple admixed samples (from a single admixture event) may simply have clustered estimated locations, and not need to draw admixture from elsewhere due to the fact that their frequencies are well described by their proximity to other admixed populations. Along these lines, it is noticeable that many of our European samples draw little admixture from elsewhere (also noted by [Hellenthal et al., 2014] using a different approach), despite evidence of substantial ancient admixture [Lazaridis et al., 2014]. This may reflect the fact that all of the European samples are affected by the admixture events, and are relatively over-represented in our sample. However, this may also simply reflect the fact that the admixture is ancient, and that the ancient populations that took part in these events are not well represented by our extant sampling. Reassuringly, we see multiple cases where similarly admixed populations (Central Asians, Middle Eastern, and North African) populations are separately identified as admixed. This suggests that geogenetic clustering (in lieu of drawing admixture) of populations that share similar histories of admixture is not a huge concern (at least in some cases). The method could in theory be modified to allow geogenetically proximal populations to draw from the same admixture event; however, this may be difficult to make fully automated.

In this paper, we have treated the loci in the dataset as independent, and, where necessary, we have thinned empirical datasets to decrease LD between loci. One possible approach that avoids the necessity of thinning the data would be to calculate the sample covariance in large (e.g., megabase), non-overlapping windows along the genome, then average those sample covariances across all windows. Another approach is to use empirical LD between loci to estimate the effective number of independent loci in the dataset, and use this quantity as the number of degrees of freedom in the Wishart likelihood calculation. Additionally, although we have focused on the covariance among alleles at the same locus, linkage disequilibrium (covariance of alleles among loci) holds rich information about the timing and source of admixture events [Chakraborty and Weiss, 1988, Moorjani et al., 2013, Hellenthal et al., 2014, Gravel, 2012] as well as information about isolation by distance [Ralph and Coop, 2013]. Just as population graph approaches have been extended to incorporate information from LD [Loh et al., 2013], a spatial covariance approach could be informed by LD. A null model inspired by models of LD under isolation by distance models [De and Durrett, 2007, Barton et al., 2013] could be fitted, allowing the covariance among alleles to decay with their geographic distance and the recombination distance between the loci. In such a framework, sources and time-scales of admixture could be identified through unusually long-distance LD between geographically separated populations.

The landscape of allele frequencies on which the location of populations that were the source of population’s admixture are estimated is entirely informed by the placement of other modern samples, even though the admixture events may have occurred many generations ago. This immediately leads to the caveat that, instead of “location of the parental population,” we should refer to the “location of the closest descendants of the parental population.” The increased sequencing of ancient DNA [see Pickrell and Reich, 2014, for a recent review] promises an interesting way forward on that front, and it will also be exciting to learn where ancient individuals fall on modern maps, as well as how the inclusion of ancient individuals changes the configuration of those maps [Skoglund et al., 2014]. The inclusion of ancient DNA samples in the analyzed sample offers a way to get better representation of the ancestral populations from which the ancestors of modern samples received their admixture. However, it is also possible to model genetic drift as a spatiotemporal process, in which covariance in allele frequencies decays with distance in both space and in time. We are currently exploring using ancient DNA samples as ‘fossil calibrations’ on allele frequency landscapes at points in the past, so that modern day samples may draw admixture from coordinates estimated in spacetime.

## Methods

Here we describe in more detail the algorithm we use to estimate the posterior distribution defined by (7) of the population locations, *G*, their sources of admixture, *G*^*^, their admixture proportions, *w*, their independent drift parameters, *η*, and the parameters of the model of isolation by distance, 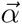. First, we give the exact form of the covariance matrix we use, and then describe the Markov chain Monte Carlo algorithm that samples parameter values from the posterior distribution.

### The standardized sample covariance

As motivation, consider several randomly mating (Wright-Fisher) populations that all split from an ancestral population in which a neutral allele is present at frequency *E*_*f*_, and then subsequently exchange migrants. Since the allele is neutral, the mean change in its frequency in each population after *t* generations is zero, and if *t* is much smaller than the population size (so the frequencies remain close to 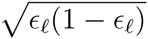, the variance is proportional to 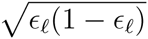. Conveniently, additional variance introduced by binomial sampling of alleles is also proportional to would then be natural to consider the covariance matrix of

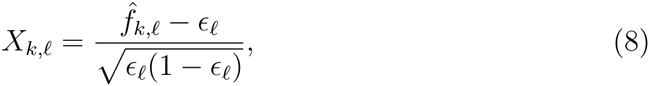

since these standardized allele frequencies would be independent if the loci are unlinked, and would have mean zero and variance independent of the sample sizes or allele frequencies. The central limit theorem would then imply that in the limit of a large number of loci, the sample covariance matrix *X*^*T*^ *X* is Wishart with degrees of freedom equal to the number of loci and mean determined by the pattern of migration.

Although the conditions are not strictly met, these theoretical considerations indicate that such a normalization may be a reasonable thing to do, even after substituting the empirical mean allele frequency 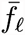 in place of *ϵ*_*f*_, which is what we do to define 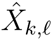 in equation (1). Recall that the sample allele frequency at locus *l* in population *k* is given by 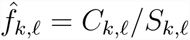, where *C*_*k,l*_ is the number of (arbitrarily chosen) counted alleles, and *S_k,l_* is the total number of sampled alleles. As sample size may vary across loci, we first calculate 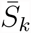, the mean sample size in population *k*, as 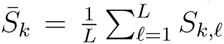. We then compute the global mean allele frequency at locus *l* as

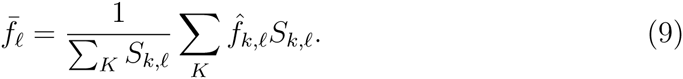

If sample size were constant across all loci in each population, this would be equivalent to defining the variance-normalized sample frequencies

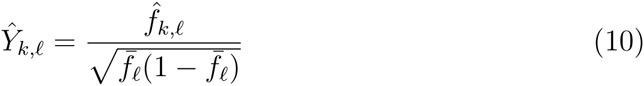

and writing 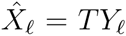 where *T* is the mean centering matrix whose elements are given by

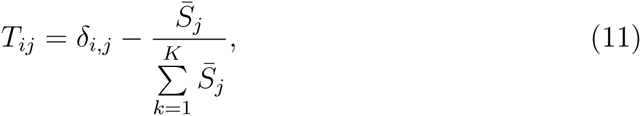

where *δ*_*i,j*_ = 1 if *i* = *j* and is 0 otherwise. If the covariance matrix of *Y* is Ω^*^, then the covariance matrix of 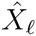 would be *T* ^*T*^ Ω^*^*T*. Since allowing *T* to vary by locus would be computationally infeasible, we make one final assumption, that the covariance matrix of the standardized frequencies 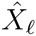 at each locus is given by *T* ^*T*^ Ω^*^*T*. This makes it inadvisable to include loci for which there are large differences in sample sizes across populations. This mean centering acts to to reduce the covariance among populations in 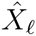 compared to 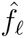 and can induce negative covariance between more unrelated populations (as, across loci, they are often on opposite sides of the mean).

Additionally, the covariance matrix of the standardized frequencies has rank *K–*1 rather than *K*, and so the corresponding Wishart distribution is singular. To circumvent this problem we compute the likelihood of a (*K–*1)-dimensional projection of the data. Any projection would do; we choose a projection matrix ψ by dropping the last column of the orthogonal matrix in the QR decomposition of *T*, and compute the likelihood of the empirical covariance matrix of allele frequencies 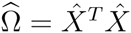 as

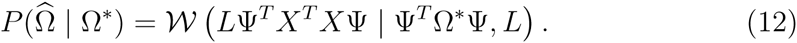

### Markov chain Monte Carlo Inference Procedure

The inference algorithm described here may be used to estimate the parameters with any of these held fixed, for instance: (1) population locations are fixed, and they do not draw any admixture; (2) population locations are estimated, but not admixture; (3) populations may draw admixture, but their own locations are fixed; or (4) population locations and admixture are both estimated. The free parameters for each of options are given in Table 1.

**Table 1:** List of models that may be specified using SpaceMix, along with the number and identity of free parameters in each.

| Model | # of Free Parameters | Parameters |
| --- | --- | --- |
| (1) stationary population locations, no admixture | $K + 3$ | $\alpha_0, \alpha_1, \alpha_2, \eta$ |
| (2) inferred population locations, no admixture | $2K + 3$ | $\alpha_0, \alpha_1, \alpha_2, \eta, G$ |
| (3) stationary population locations, inferred admixture | $2K + 3$ | $\alpha_0, \alpha_1, \alpha_2, \eta, G^*, w$ |
| (4) inferred population locations, inferred admixture | $3K + 3$ | $\alpha_0, \alpha_1, \alpha_2, \eta, G, G^*, w$ |

Although we anticipate most empirical researchers will be interested in the joint inference of a geogenetic map with admixture (Model 4), we have presented these models separately, as we believe each have their own utility. Model 1 and Model 3 can each be used to infer landscapes of allele frequencies, upon which genotyped individuals can be probabilistically placed (following [Wasser et al., 2004]). This application may be useful to determine the geographic origin of potentially contraband biological samples (e.g., ivory), or the most likely source of museum specimens missing sampling metadata. Model 3 has the potential to improve the performance of these spatial assignment methods over Model 1, as the inclusion of admixture in the model may allow for more accurate inference of allele frequency surfaces. Model 2 directly parallels Principal Component Analysis, and can also be used in a model comparison framework with Model 4 to formally test for the presence of admixture in the sampled dataset.

Below, we outline the inference procedure for the most parameter-rich model (inference on both population locations, their sources of admixture, and the proportions in which they draw admixture, in addition to inference of the parameters of the spatial covariance function). A table of all parameters, their descriptions, and their priors is given in Table 2.

**Table 2:** List of parameters used in the SpaceMix models, along with their descriptions and priors. locations *G*^(*obs*)^. 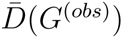 is the mean of the pairwise distances between observed

| Parameter | Description | Prior |
| --- | --- | --- |
| $\alpha_0$ | controls the sill of the covariance matrix | $\alpha_0 \sim \text{Exp}(\lambda = 1/100)$ |
| $\alpha_1$ | controls the rate of the decay of covariance with distance | $\alpha_1 \sim \text{Exp}(\lambda = 1)$ |
| $\alpha_2$ | controls the shape of the decay of covariance with distance | $\alpha_2 \sim U(0.1, 2)$ |
| $\eta_k$ | the nugget in population $k$ (population specific drift parameter) | $\eta_k \sim \text{Exp}(\lambda = 1)$ |
| $G_k$ | the geogenetic location of population $k$ | $G_k \sim \mathcal{N}(\mu = G_k^{(obs)}, \sigma = \frac{1}{2}\bar{D}(G^{(obs)}))$ |
| $w_k$ | the proportion of admixture in population $k$ | $2w_k \sim \beta(\alpha = 1, \beta = 100)$ |
| $G_k^*$ | the geogenetic location of the source of admixture in population $k$ | $G_k^* \sim \mathcal{N}(\mu = \bar{G}^{(obs)}, \sigma = 2\bar{D}(G^{(obs)}))$ |

We now specify in detail the Markov chain Monte Carlo algorithm we use to sample from the posterior distribution on the parameters, for Bayesian inference.

We assume that the user has specified the following data:

- the allelic count data, *C*, from *K* population over *L* variant loci, where *C*_*k,l*_ gives the number of observations of a given allele at locus *l* in population *k*.
- the sample size data, *S*, from *K* population over *L* variant loci, where *S*_*k,l*_ gives the total number of alleles typed at locus *l* in population *k*.

It is not necessary, but a user may also specify

- the geographic sampling locations, *G*^(*obs*)^, from each of the *K* populations, where 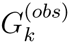 gives the longitude and latitude of the *k*^th^ sampled individual(s).

The geographic location data may be missing, or generated randomly, for some or all of the samples; if so, the spatial priors on estimated population locations, *G*, and their sources of admixture, *G*^*^ will not be tethered to the true map.

**Initiating the MCMC** We then calculate the standardized sample covariance matrix 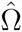 as described in the section “The standardized sample covariance” above, as well as 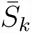, the mean sample size across loci for each population. Armed with the standardized sample covariance, the geographic sampling locations, and the inverse mean sample sizes across samples (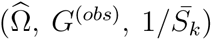), we embark upon the analysis.

To initiate the chain, we specify starting values for each parameter. We draw initial values for *α*_0_, *α*_1_, *α*_2_, *η*, and *w* randomly from their priors. We initiate *G* at user-specified geographic sampling locations and *G*^*^ at randomly drawn, uniformly distributed values of latitude and longitude in the observed range of both axes.

**Overview of MCMC procedure** We use a Metropolis-Hastings update algorithm. In each iteration of the MCMC, one of our current set of parameters Θ = *α*_0_, *α*_1_, *α*_2_, *η*, *w*, *G*, *G*^*^ is randomly chosen to be updated by proposing a new value. In the cases of *η*, *w*, *G*, *G*^*^, where each population has its own parameter, a single population, *k* is randomly selected and only its parameter value (e.g. *η*_*k*_) is chosen to be updated. Below we outline the proposal distributions for each parameter. This gives us a proposed update to our set of parameters Θ^*t*^, which differs from Θ at only one entry.

The set of locations of populations and their sources of admixture specify a pairwise geographic distance matrix *D*, which, given the current 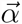 and *η* parameters, gives the admixed covariance matrix described in (6), Ω^*^. The likelihood of the two sets of parameters Θ and Θ ′, calculated with (12) and the priors of Table 2, combine to give the Metropolis-Hastings ratio, *R*, the probability of accepting the proposed parameter values Θ ′:

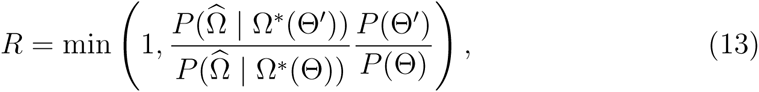

Note that all of our moves, described below, are symmetric, so the Hastings ratio is always 1. If we accept our proposed move, Θ is replaced by Θ′ and this is recorded, otherwise Θ′ is discarded and we remain at Θ.

**Updates for 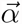** *η***, and** *w* We propose updates to the values of the 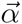, *η*, and *w* parameters via a symmetric normal density with mean zero and variance given by a tuning parameter specific to that parameter. For example, 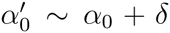, where 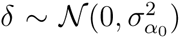 and *σ*_*α0*_ is the tuning parameter for *α*_0_. For *η* and *w*, each of which consists of *K* parameters, each parameter receives its own independent tuning parameter. If the proposed move takes the parameter outside the range of its prior, the move is rejected and we do not move in that iteration of the MCMC.

**Updates for geographic coordinates** *G* **and** *G*^*^ Updates to the location parameters, *G* and *G*^*^, are somewhat more complicated due to the curvature of the Earth. Implementing updates via a symmetric normal density on estimated latitude and longitude directly would have the drawback of a) being naive to the continuity of the spherical manifold and b) vary the actual distance of the proposed move as a function of the current lat/long parameter values (e.g., a 1° change in longitude at the equator is a larger distance than at the North Pole).

Instead, we propose a bearing and a distance traveled, and, given these two quantities and a starting position, calculate the latitude and longitude of the proposed update to the location. For example, in an update to the location of population *i*, *G*_*i*_, we propose a distance traveled Δ_*G*_*i*__ where, e.g., Δ_*G*_*i*__ ∼ |𝒩(0, *σ*_*G*_*i*__)| and a bearing, *γ*, where *γ ~ U* (0, 2*p*). Then we use the following equations to calculate the latitude and longitude of the proposed location:

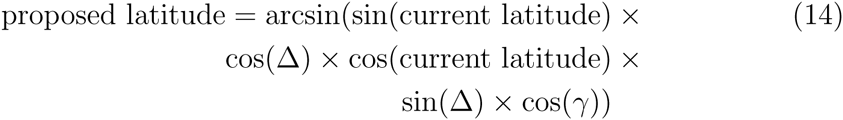

and

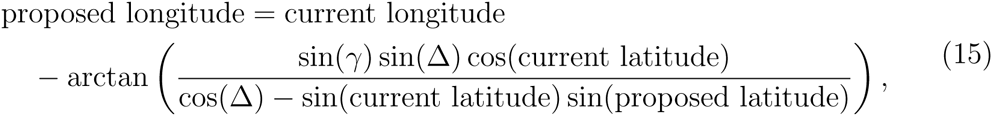

where latitude and longitude are given in radians and are taken mod 2*p*. As with *η* and *w*, each population’s location and admixture source location parameters have their own tuning parameters.

**Adaptive Metropolis-within-Gibbs proposal mechanism** We use an adaptive Metropolis-within-Gibbs proposal mechanism on each parameter [Roberts and Rosenthal, 2009, Rosenthal, 2011]. This mechanism attempts to maintain an acceptance proportion for each parameter as close as possible to 0.44 [optimal for one-dimensional proposal mechanisms; Roberts et al., 1997, Roberts and Rosenthal, 2001]. We implement this mechanism by creating, for each variable *i*, an associated variable *ζ*_*i*_, which gives the log of the standard deviation of the normal distribution from which parameter value updates are proposed. As outlined above, in the cases of {*η*, *w*, *G*, *G*^*^}, for which each population receives a free parameter, each population gets its own value of *ζ*.

When we start our MCMC, *ζ*_*i*_ for all parameters is initiated at a value of 0, which gives a proposal distribution variance of 1. We then proceed to track the acceptance rate, *r*_*i*_ for each parameter in windows of 50 MCMC iterations, and, after the *n*th set of 50 iterations, we adjust *ζ*_*i*_ by an “adaption amount”, which is added to *ζ*_*i*_ if the acceptance proportion in the *n*th set of 50 iterations 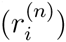 has been above 0.44, and subtracted from *ζ*_*i*_ if not. The magnitude of the adaption amount is a decreasing function of the index *n*, so that updates to *ζ*_*i*_ proceed as follows:

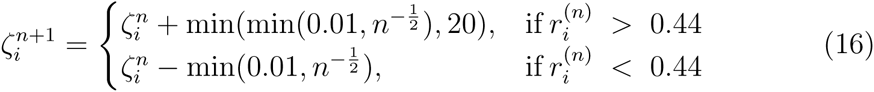

We choose to cap the maximum adaption amount at 20 (which is the equivalent of capping the variance of the proposal distribution at 4.85 × 10^8^) to avoid proposal distributions that offer absurdly large or small updates. This procedure, also referred to as “auto-tuning”, results in acceptance rate plots like those shown in Fig. S22, and in more efficient mixing and decreased autocorrelation time of parameter estimates in the MCMC.

### Simulations

We ran our simulations using a coalescent framework in the program ms [Hudson, 2002]. Briefly, we simulated populations on a lattice, with nearest neighbor migration rate *m*_*i,j*_, as well as migration on the diagonal of the unit square at rate 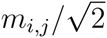. For each locus in the dataset, we used the **-s** option to specify a single segregating site, and then we simulated 10,000 loci independently, which were subsequently conglomerated into a single dataset for each scenario. For all simulations, except the “Populations on a line” scenario (Fig. S15), we sampled only every other population, and, from each population, we sampled 10 haplotypes (corresponding to 5 diploid individuals). In the “Populations on a line” scenario, we simulated no intervening populations, such that every population was sampled.

To simulate a barrier event, we divided the migration rate between neighbors separated by the longitudinal barrier by a factor of 5. To simulate an expansion event, we used the **-ej** option to move all lineages from each daughter population to its parent population at a very recent point in the past. For admixture events, we used the **-es** and **-ej** options to first (again, going backward in time) split the admixed population into itself and a new subpopulation of index *k* + 1, and second, to move all lineages in the (*k*^th^ + 1) into the source of admixture. Forward in time, this procedure corresponds to cloning the population that is the source of admixture, then merging it, in some admixture proportion, with the (now) admixed population. The command line arguments used to call 

~~~
ms
~~~

 for a single locus for each simulation are included in the Appendix.

### Intuition on identifiability of admixture parameters

A natural concern is whether all of the parameters we infer are separately identifiable, most notably whether population locations, admixture locations, and proportions can be estimated. That is, if a population has received some admixture from another population, what is to stop it from having an estimated location near that population in geogenetic space to satisfy its increased resemblance to that population, rather than drawing admixture from that location? We do not provide a formal proof, but here build and illustrate some relevant intuition.

Admixture is identifiable in our model because there are covariance relationships among populations that cannot simply be satisfied by shifting population locations around (as demonstrated by the tortured nature of Fig. 2b). Consider the simple spatial admixture scenario shown in Fig. 11. Populations *A*–*D* are arrayed along a line, but there is recent admixture from *D* into *B* (such that 40% of the alleles assigned to B are sampled from location *D*). The lines show the expected covariance under isolation by distance that each population (*A*, *C*, or *D*, as indicated by line color) has with a putative population at a given distance. The dots show the admixed covariance between *B* and the three other populations, as well as *B*’s variance with itself (*B*-*B*) as specified by equation (6), with no nugget or sampling effect.

**Figure 11:**
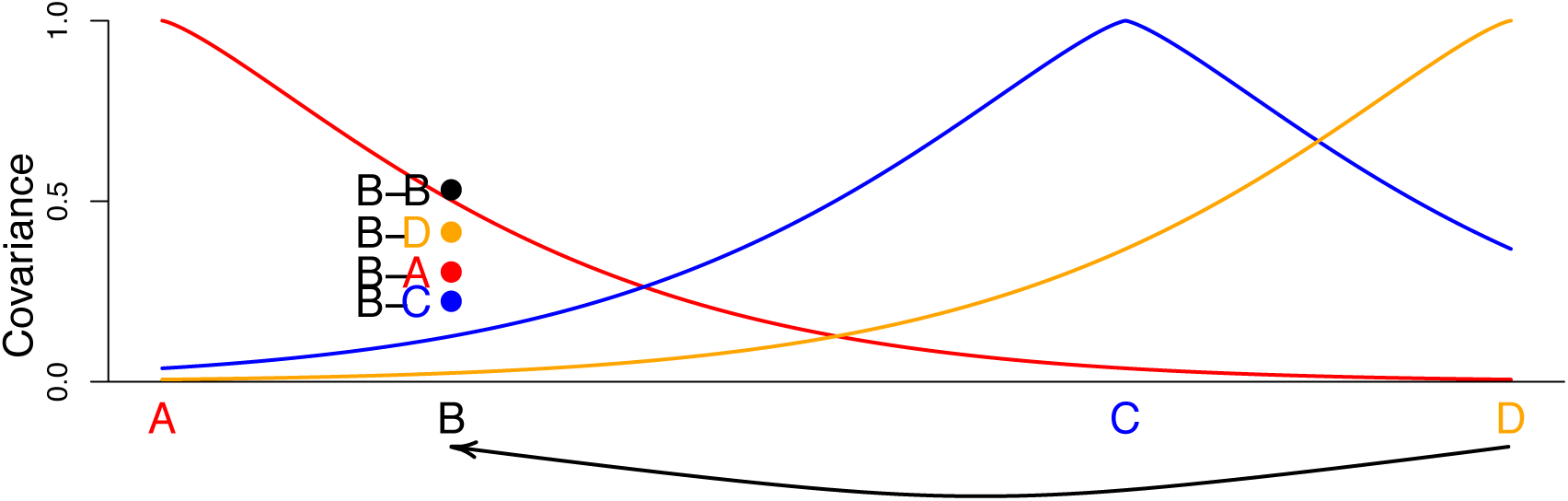
Lines show the covariances populations *A*, *C*, and *D* would have with population *B* as a function of *B*’s location, with no admixture, under the parametric form of equation (2). The colored dots above ‘*B*’ show the covariances observed with *B* at that location if *B* has 40% admixture from *D*. There is no single spatial location with unadmixed covariances remotely similar to these.

Due to its admixture from *D*, *B* has lower covariance with *A* than expected given its distance, somewhat higher covariance with *C*, and much higher covariance with In addition, the variance of *B* is lower than that of the other three populations, which each have variance 1: the value of the covariance when the distance is zero. This lower variance results from the fact that the frequencies at *B* represent a mixture of the frequency at *D* and the frequency at *B* before the admixture.

Now, using this example scenario, let us return to the concern posed above: that admixture location and population location are not identifiable. For the sake of simplicity, assume that we hold the locations of *A*, *C*, and *D* constant, as well as the decay of covariance with distance (as could be the case if *A*–*D* are part of a larger analysis). The covariance relationships of *B* to the other populations cannot be simply satisfied if *B* had an estimated location near *D*, as *B* would then have a covariance with *C* that is higher, and a covariance with *A* that is lower, than that we actually observe.

Introducing admixture into the model allows it to satisfy all of these conditions: it can draw ancestry from *D* but keep part of its resemblance to *A*, it avoids *B* having an estimated location too close to *C*, and it explains *B*’s low variance. Even in the absence of a sample from population *C*, *B* is better described as a linear mixture of a population close to *A* and *D*. However, there are specific scenarios in which a limited sampling scheme (both in size and location), can lead to tradeoffs in the likelihood between estimated population location and that of its source of admixture.

The analyses depicted in Figs. 2c, 4b, and 4c, give examples of these tradeoffs. In each, the inferred admixture proportions in the admixed populations are less than those used to simulate the data, and the model is able to explain the high covariance the admixed populations have with their sources of admixture via their inferred location, rather than just via their inferred source of admixture and admixture proportion. The reason the model explains these admixed populations’ anomalous covariance with their inferred location, rather than with their admixture source, is that we place a very harsh prior against admixture inference (Table 2). The prior is designed to make inference conservative with respect to admixture, but it has the side effect of skewing the posterior probability toward lower admixture proportions.

### Empirical Applications

Below, we describe the specifics of our analyses of the greenish warbler dataset and the global human dataset. The analysis procedure for each dataset is given here:

For each analysis,

1. Five independent chains were run for 5 × 10^6^ MCMC iterations each in which population locations were estimated (but no admixture). Population locations were initiated at the origin (i.e. at iteration 1 of the MCMC, *G*_*i*_ = (0, 0)), or at uniformly distributed coordinates between the minimum and maximum observed range of latitude and longitude, and all other parameters were drawn randomly from their priors at the start of each chain.
2. The chain with the highest posterior probability at the end of the analysis was selected and identified as the “Best Short Run”.
3. A chain was initiated from the parameter values in the last iteration of the Best Short Run. Because inference of admixture proportion and location was not allowed in the five initial runs, admixture proportions were initiated at 0 and admixture locations, *G*^*^ were initiated at the origin. This chain (the “Long Run”) was run for 10^8^ iterations, and sampled every 10^5^ iterations for a total of 1000 draws from the posterior.

For each dataset, we ran two analyses using the observed population locations as the prior on *G*. Then, to assess the potential influence of the spatial prior on population locations, we ran one analysis in which the observed locations were replaced with random, uniformly distributed locations between the observed minima and maxima of latitude and longitude. For the warbler dataset, we repeated this analysis procedure, treating each sequenced individual as its own population. For clarity and ease of interpretation, we present a full Procrustes superimposition of the inferred population locations (*G*) and their sources of admixture (*G*^*^), using the observed latitude and longitude of the populations/individuals (*G*) to give a reference position and orientation. As results were generally consistent across multiple runs for each dataset regardless of the prior employed, we (unless stated otherwise) present only the results from the ‘random’ prior analyses.

Finally, we compared the SpaceMix map to a map derived from a Principal Components Analysis [Patterson et al., 2006]. For this analysis, we calculated the eigendecomposition of the mean-centered allelic covariance matrix, then plotted individuals’ coordinates on the first two eigenvectors [e.g. Novembre et al., 2008]. For consistency of presentation, we show the full Procrustes superimposition of the PC coordinate space around the geographic sampling locations.

### Treemix Comparisons

To contrast our approach to tree-based approaches to admixture we applied 

~~~
TreeMix
~~~

 [Pickrell and Pritchard, 2012] to our spatial simulations. We took the ms output on which we had also run Spacemix (Figs. 1,2,4), and converted it into 

~~~
TreeMix
~~~

 format. We also included an outgroup sequence for each dataset, which consisted of a single haploid individual who carried the 0 (ancestral allele) at every locus. We ran TreeMix without migration edges on the processed 

~~~
ms.treemix.file.gz
~~~

 file to construct the initial tree, which was rooted using the myoutgroup sequence using the following command:

~~~
treemix -i ms.treemix.file.gz -root myoutgroup -o treemix output
~~~

We then sequentially added admixture migration edges, using the following command to add another edge to the existing tree (“ 

~~~
prev.treemix
~~~

”):

~~~
treemix -i ms.treemix.file.gz -root myoutgroup -g prev.treemix.vertices.gz prev.treemix.edges.gz -m 1 -o treemix output
~~~

The 

~~~
TreeMix
~~~

 graphs and residual covariance matrices were visualized using the scripts provided with 

~~~
TreeMix
~~~

. The code to process and plot the data is provided in the Appendix.

In Fig. S23 we show the tree and admixture graphs produced when TreeMix is run on a lattice stepping stone model (a smaller scale version of this exercise was previously done by Pickrell and Pritchard [2012]). The tree produced by running TreeMix is rake-like, showing the lack of deep shared sub-division. However, while the tree captures some features of isolation by distance, (e.g., neighboring samples are often sister to each other), the tree structure forces many unnatural splits of geographically neighboring populations [as was previously found; Pickrell and Pritchard, 2012, their Figure S14]. The admixture migration arrows act to mitigate the strongest departures from the tree, such as geographically neighboring samples that were forced into separate places on the tree, but are unable to fully accommodate the spatial relationships between samples. Different runs of 

~~~
TreeMix
~~~

 on the same dataset result in quite different trees and orders of migration events, reflecting both the high degree of symmetry in our simulated samples on a grid, and also the poor fit of the tree model to spatial data. In Figs. S24 and S25, we also present the results of 

~~~
TreeMix
~~~

 run on our expansion and barrier simulations. In Figs. S26, S27, and S28, we show the application of 

~~~
TreeMix
~~~

 to the scenarios simulated with admixture events. For none of our scenarios was the true admixture the first migration edge added; in fact, only for the corner admixture scenario was the true admixture event in the first three edges added. This reflects the fact that 

~~~
TreeMix
~~~

 has to add migration edges to cope with the residual covariance induced simply due to the poor fit of a tree to spatially simulated data, and so misses more subtle (but real) admixture events. The poor performance of 

~~~
TreeMix
~~~

 here is the result of the spatially simulated data not conforming to the underlying assumption of the 

~~~
TreeMix
~~~

 tree-like model.

## Acknowledgements

We would like to gratefully acknowledge the Coop Lab, Yaniv Brandvain, Razib Khan, Dan Runcie, Brad Shaffer, Marjorie Weber, and Will Wetzel for helpful discussions. We also thank Miguel Alcaide, Cristian Capelli, Garrett Hellenthal, and Darren Irwin for sharing data. This work was supported in part by the National Science Foundation under award number NSF #1262645 (DBI) to PR and GC, the National Institute of General Medical Sciences of the National Institutes of Health under award numbers NIH RO1GM83098 and RO1GM107374 to GC, and the National Science Foundation under award numbers NSF # 1148897 and # 1402725 to GB.

## Supplementary Materials

**Figure S1:**
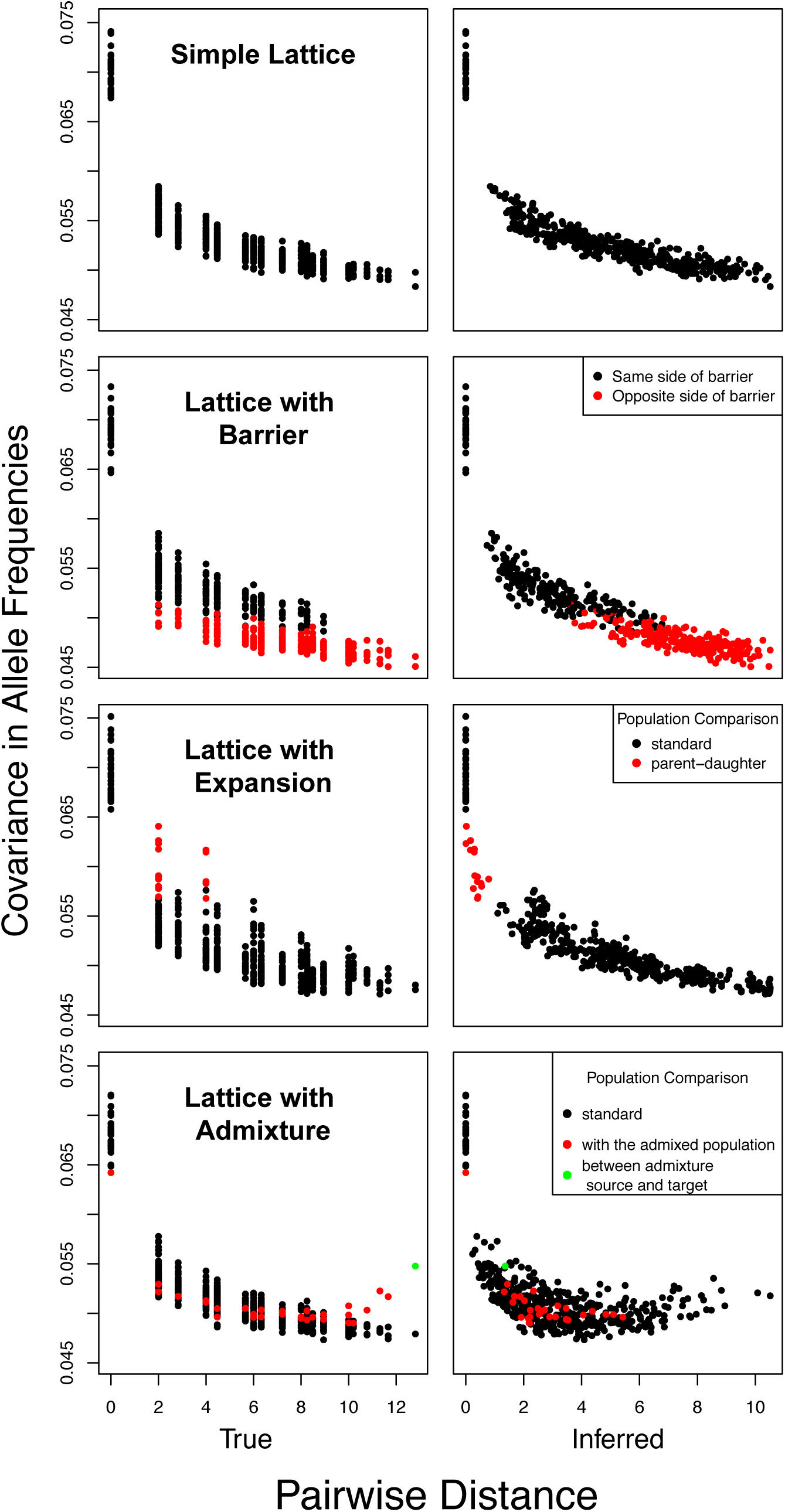
Decays in covariance for four different simulation scenarios (from top to bottom: simple lattice; lattice with barrier; lattice with expansion; lattice with admixture). Left column: sample covariance plotted against observed pairwise distance. Right column: sample covariance plotted against inferred geogenetic distance.

**Figure S2:**
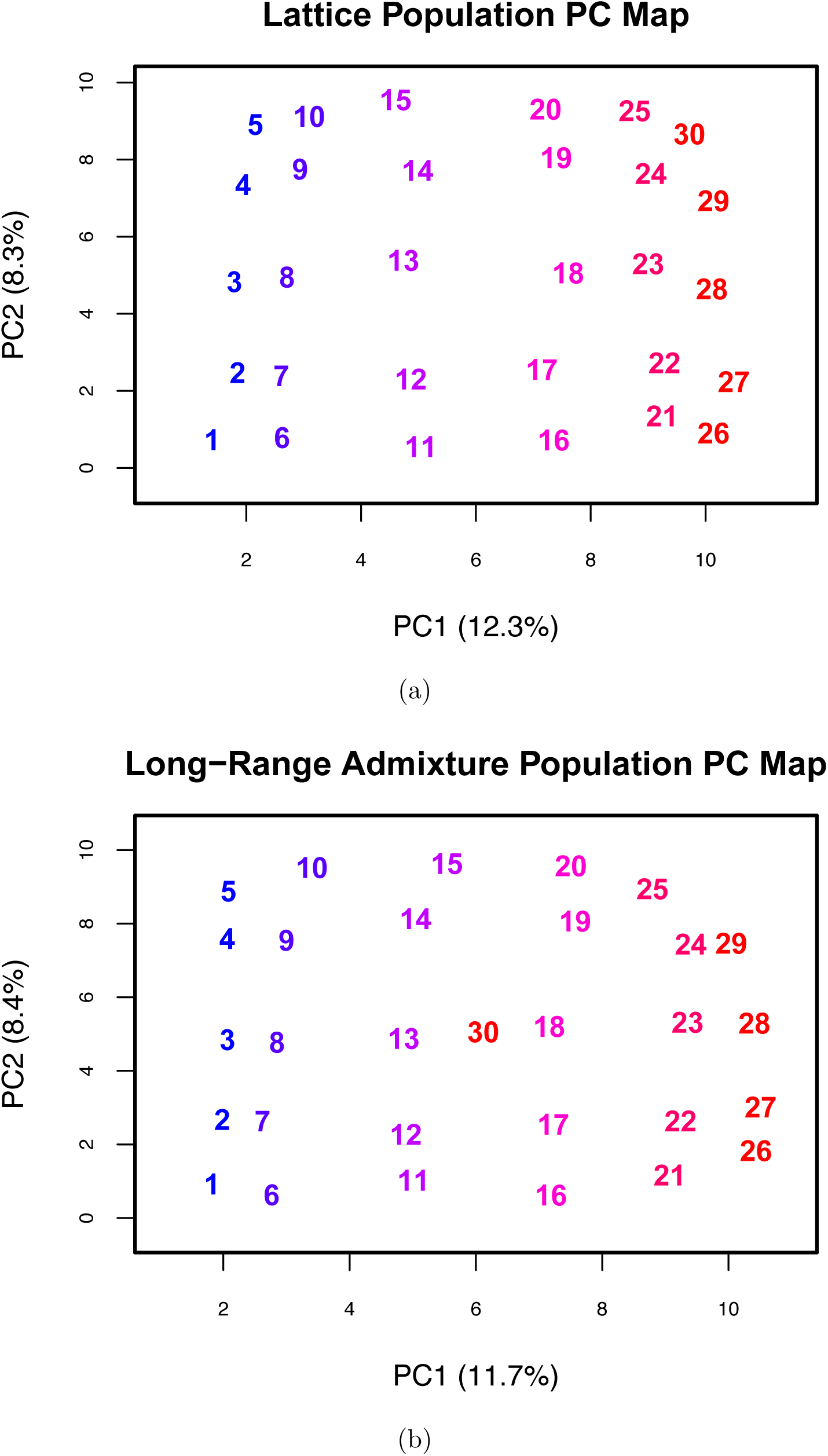
Plots of the first two Principal Component axes (with variance explained labeled on the relevant axes) of the mean-centered covariance matrix from simulated spatial scenarios. a) the basic lattice scenario shown in 1a. b) the lattice scenario with admixture from Population 1 into Population 30, shown in 2a

**Figure S3:**
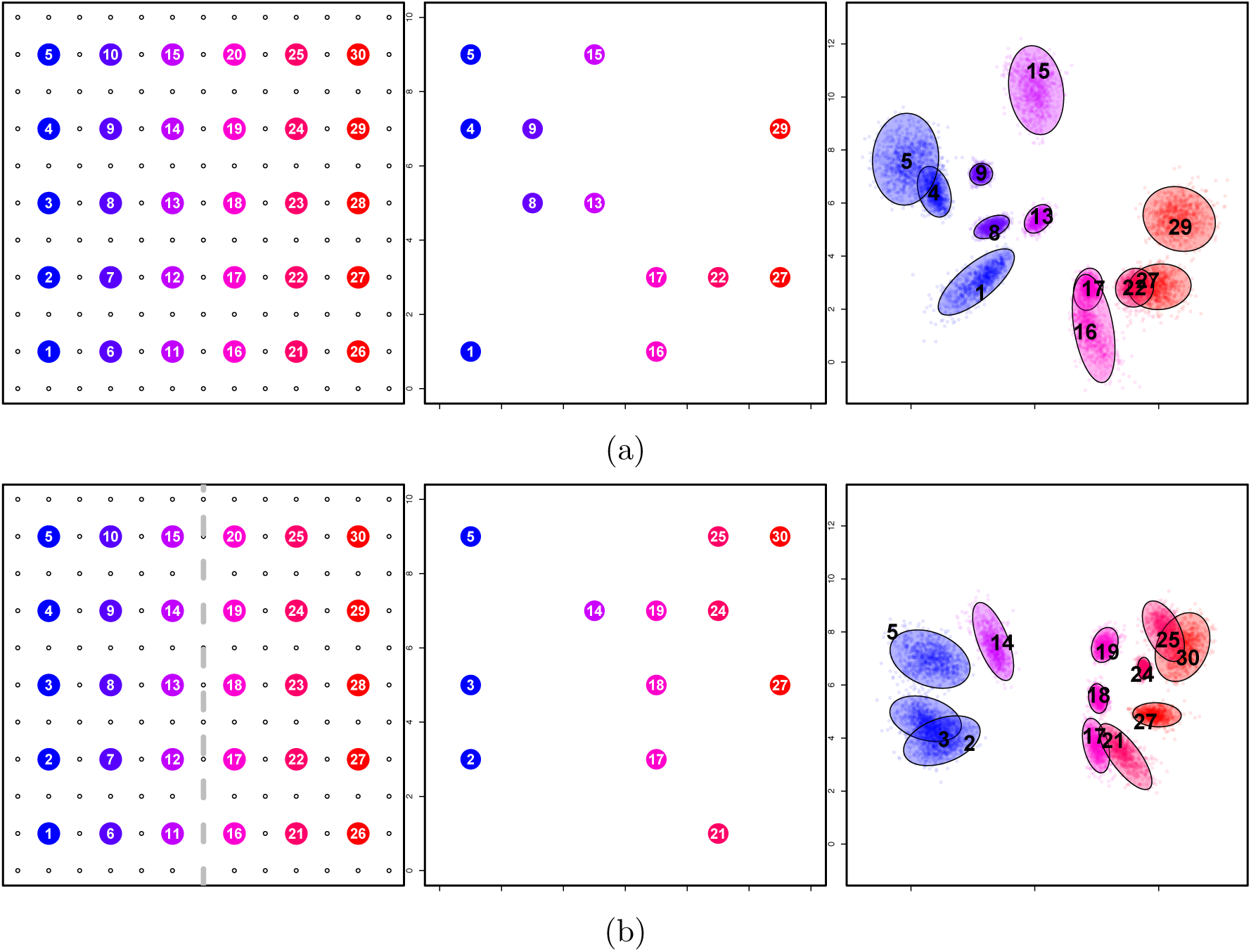
The effect of uneven sampling on inference of geogenetic maps. a) inference using a dataset with a random subsample of 12 populations from the simple lattice scenario with nearest neighbor migration; b) inference using a dataset with a random subsample of 12 populations from the barrier scenario with nearest neighbor migration and a barrier down the center line of longitude.

**Figure S4:**
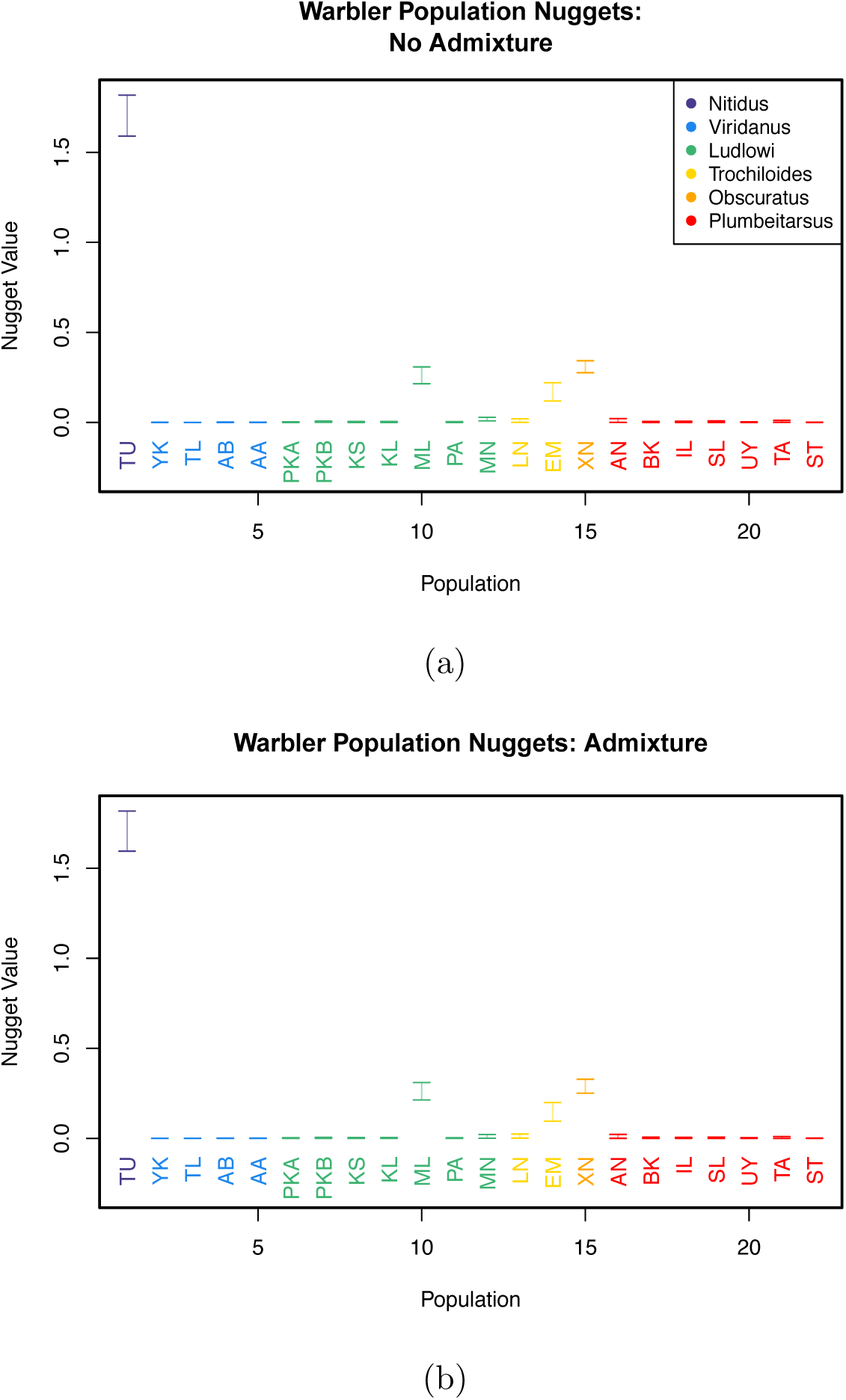
Credible intervals on estimated warbler population nugget parameters. a) analysis without admixture; b) analysis with admixture.

**Figure S5:**
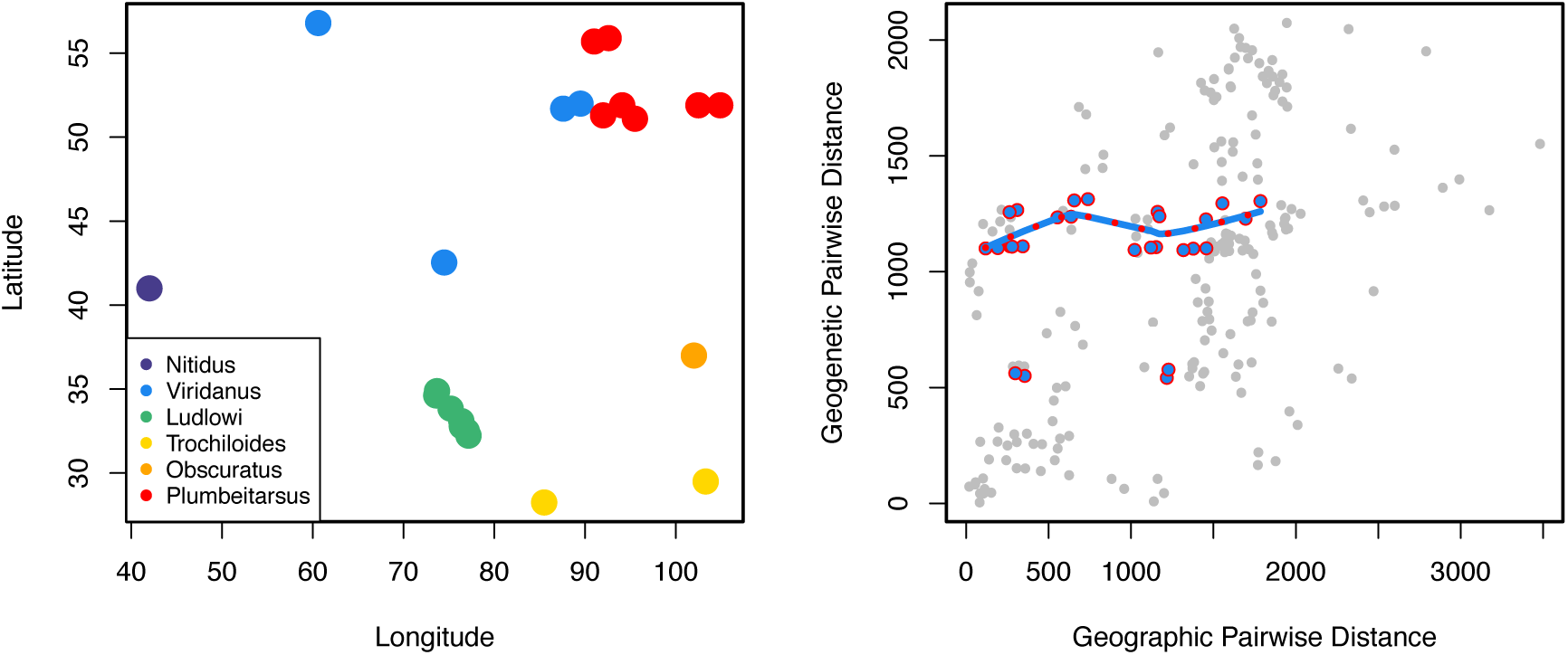
Comparing geographic to geogenetic pairwise distance between warbler populations: a) observed population coordinates; b) pairwise geographic (great-circle) distance between populations compared to that between their geogenetic locations. The highlighted points show distances between populations from the *plumbeitarsus* and *viridanus* subspecies. Notice that, regardless of their observed distance, their geogenetic separations are roughly constant, and much larger than the geographic distance between them.

**Figure S6:**
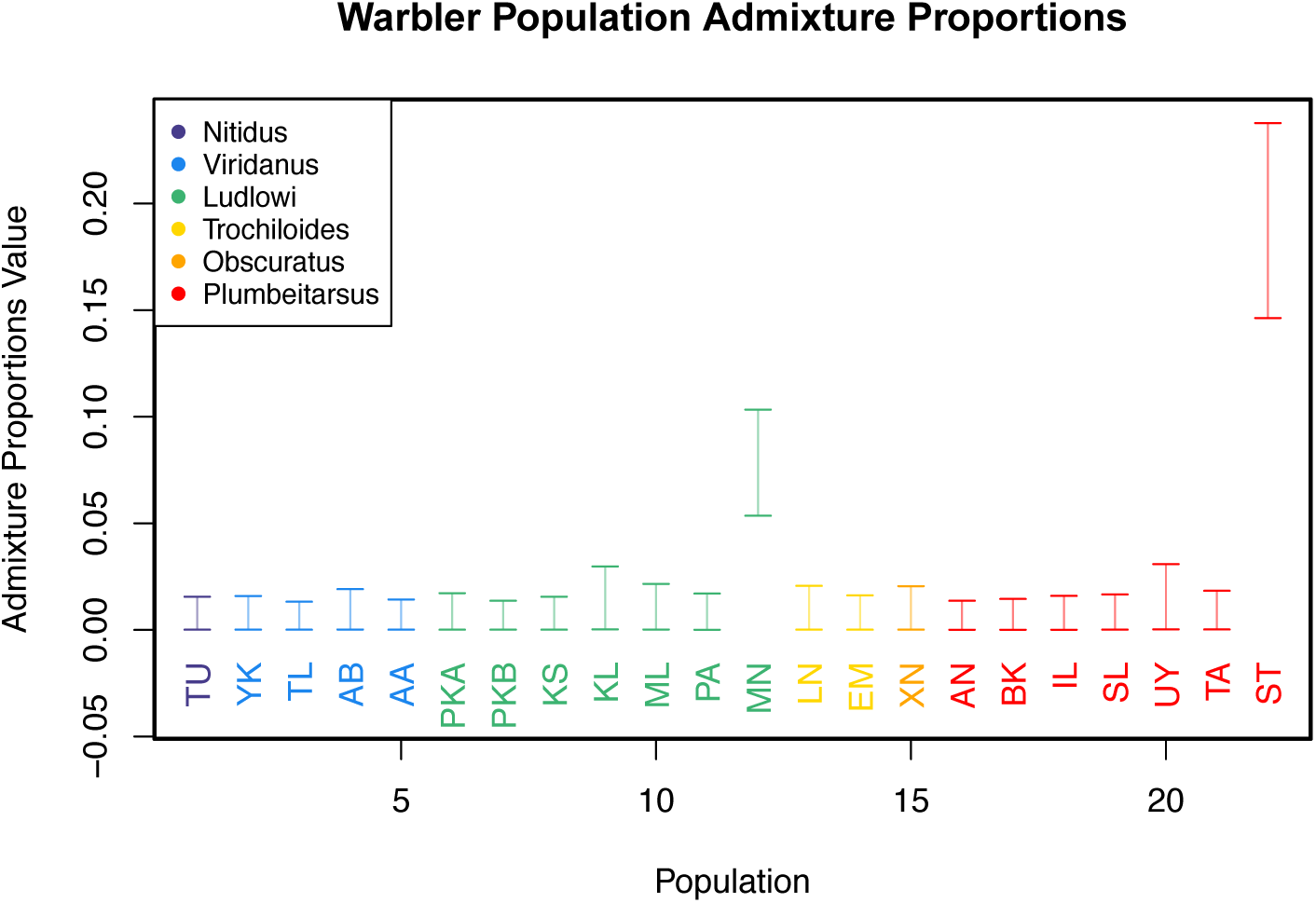
Credible intervals on estimated warbler population admixture proportion parameters.

**Figure S7:**
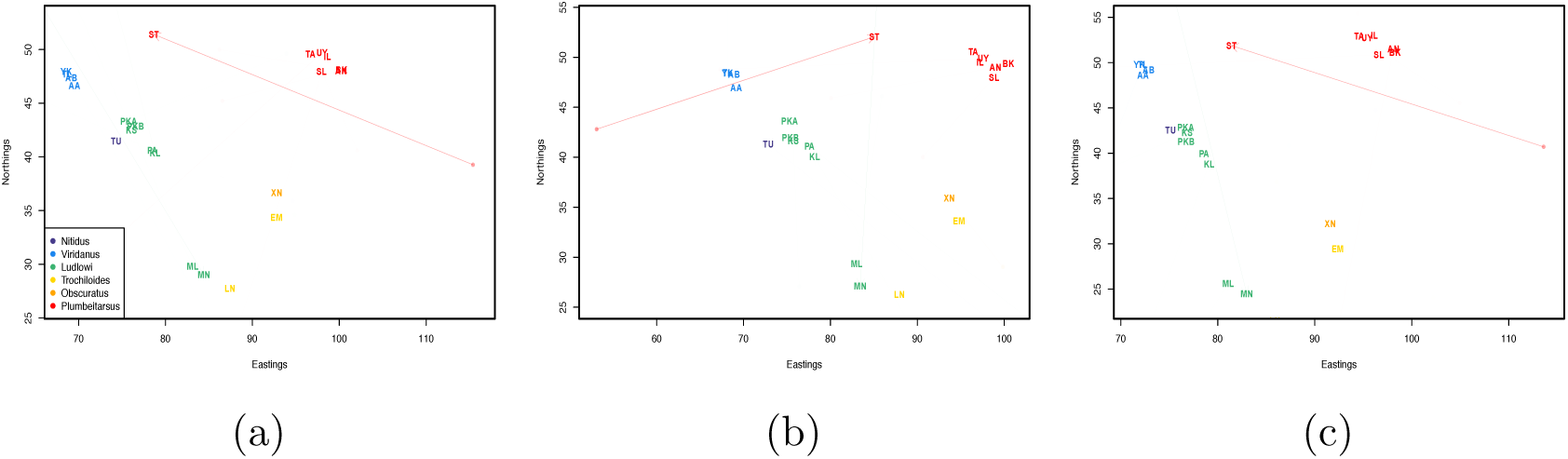
Comparison of inferred maps from three independent analyses. (a,b) Results from analysis using observed locations as priors on population locations. (c) Results from analysis using random, uniformly distributed locations within the observed range of latitude and longitude as priors on population locations.

**Figure S8:**
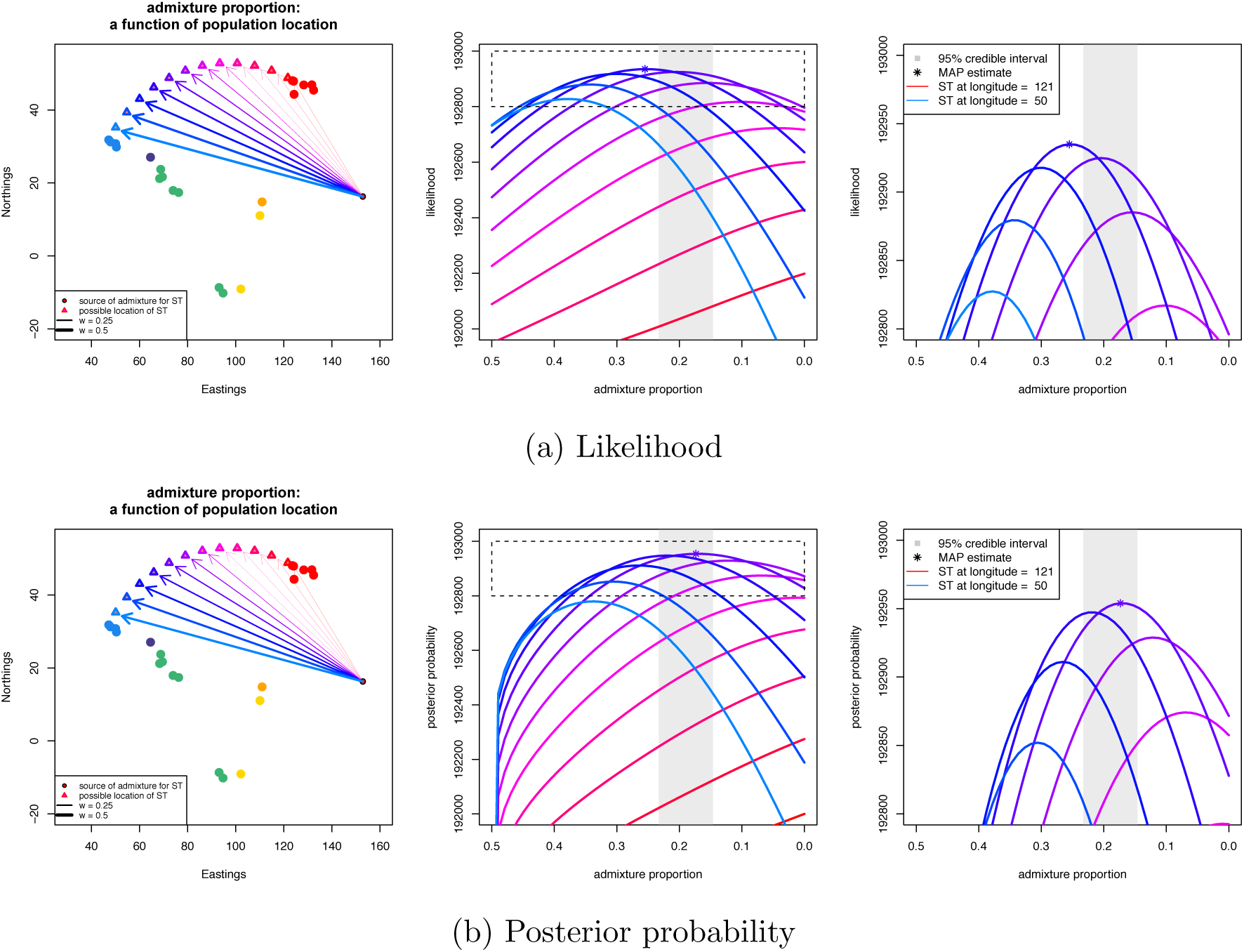
Likelihood surfaces for different placements of population ST between *plumbeitarsus* and *viridanus* clusters: (a) log likelihood surface; (b) posterior probability surface, incorporating the priors. The maximum a posteriori estimate (MAP) is shown as a star.

**Figure S9:**
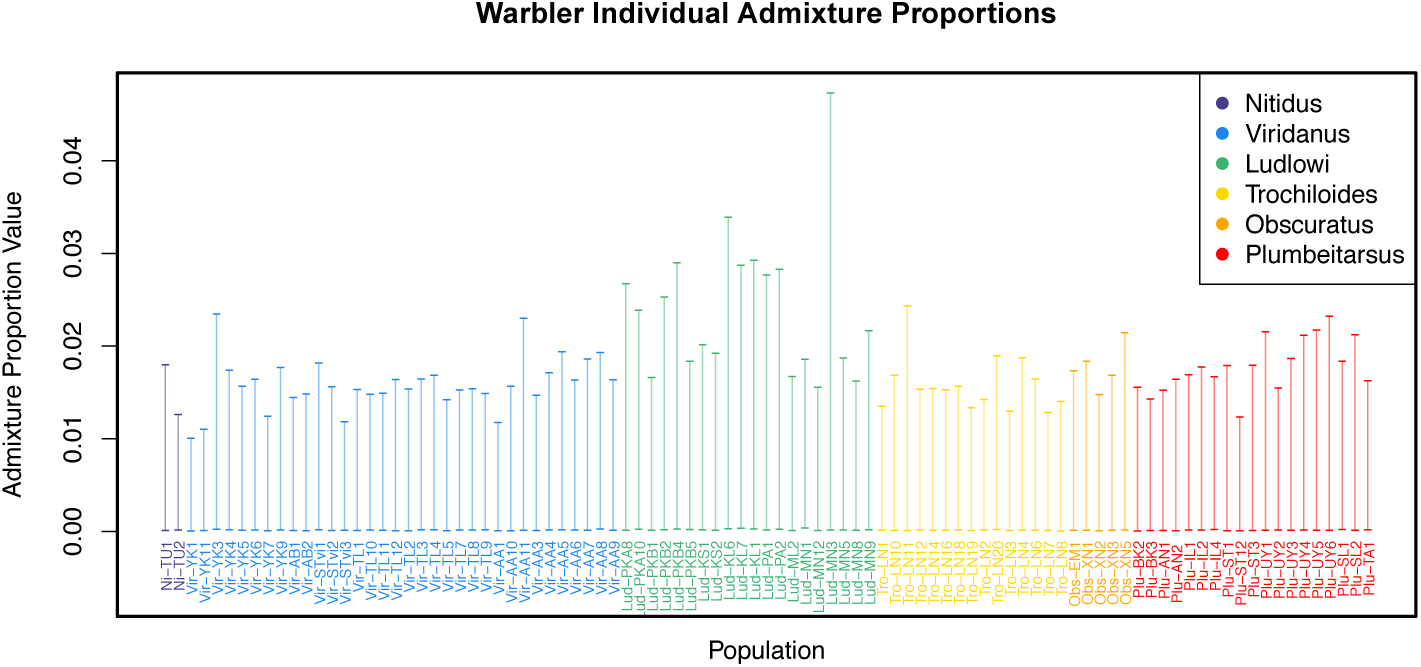
Credible intervals on estimated warbler individual admixture propor-tion parameters.

**Figure S10:**
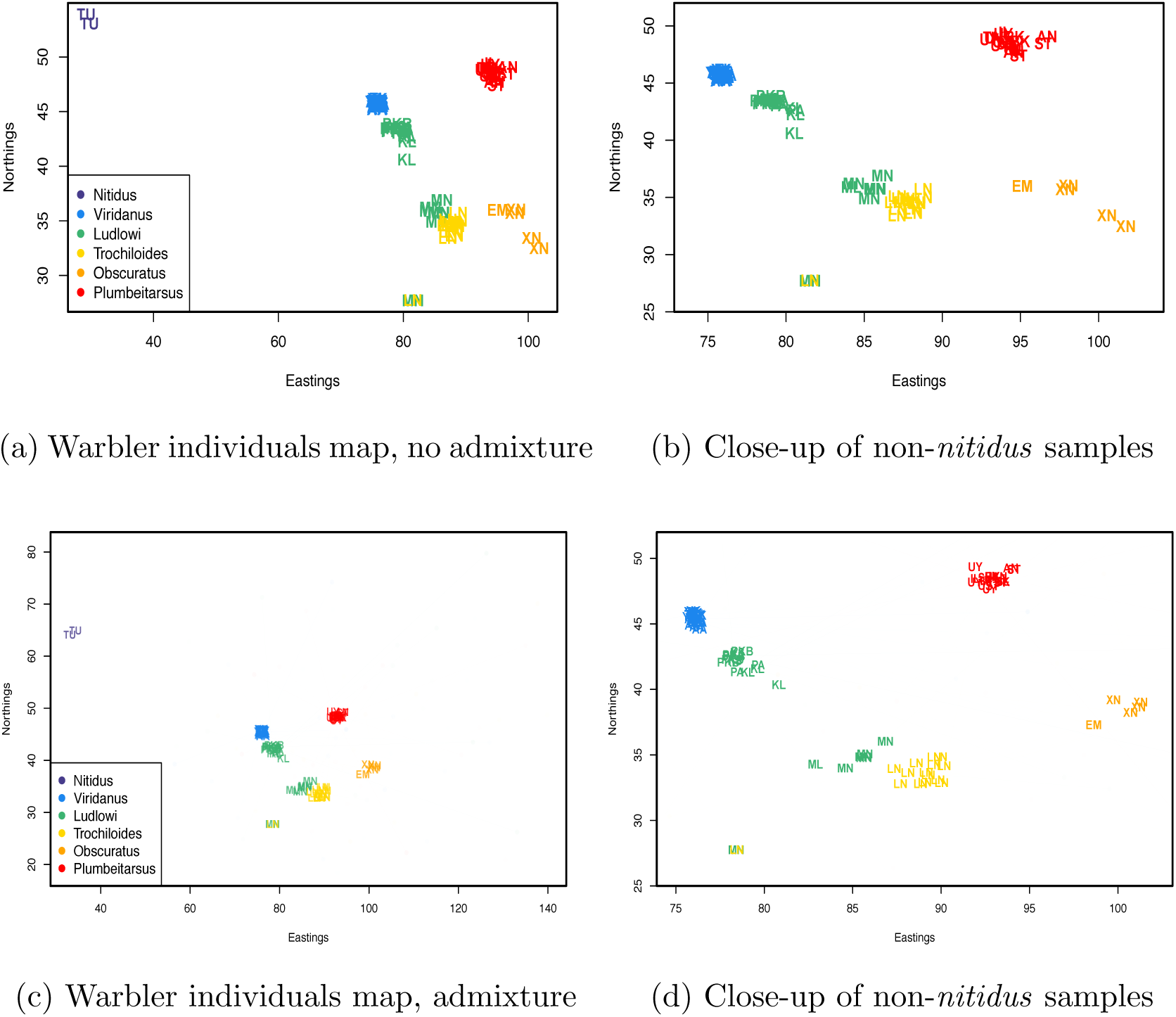
Inferred maps for warbler individuals, colored by subspecies under analyses with and without admixture inference. a) map inferred without admixture; (b) close-up of all non-*nitidus* samples in non-admixture map; c) map inferred with admixture; d) close-up of all non-*nitidus* samples in the admixture map.

**Figure S11:**
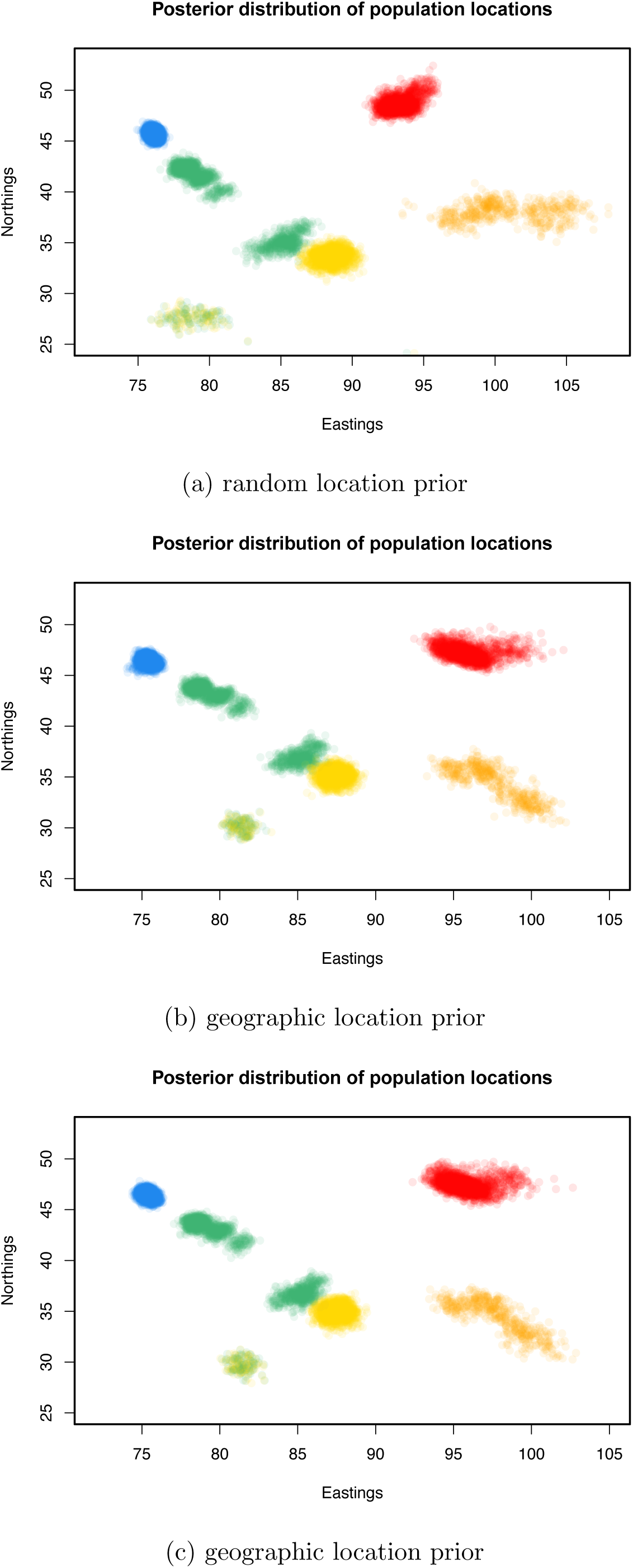
Maps of the posterior distributions on population locations in three separate SpaceMix analyses on the warbler individual dataset.

**Figure S12:**
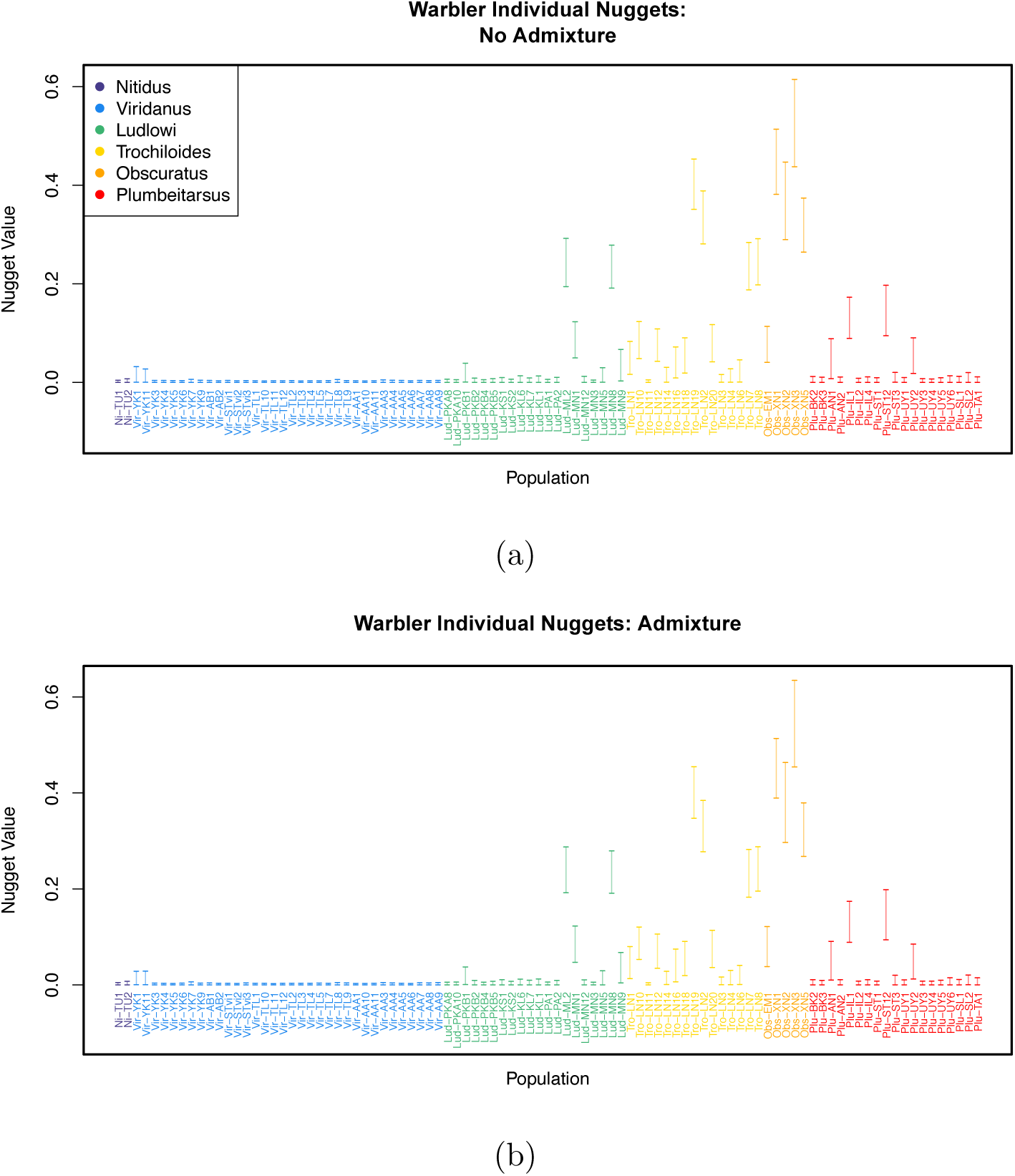
Credible intervals on estimated warbler individual nugget parameters. a) analysis without admixture; b) analysis with admixture.

**Figure S13:**
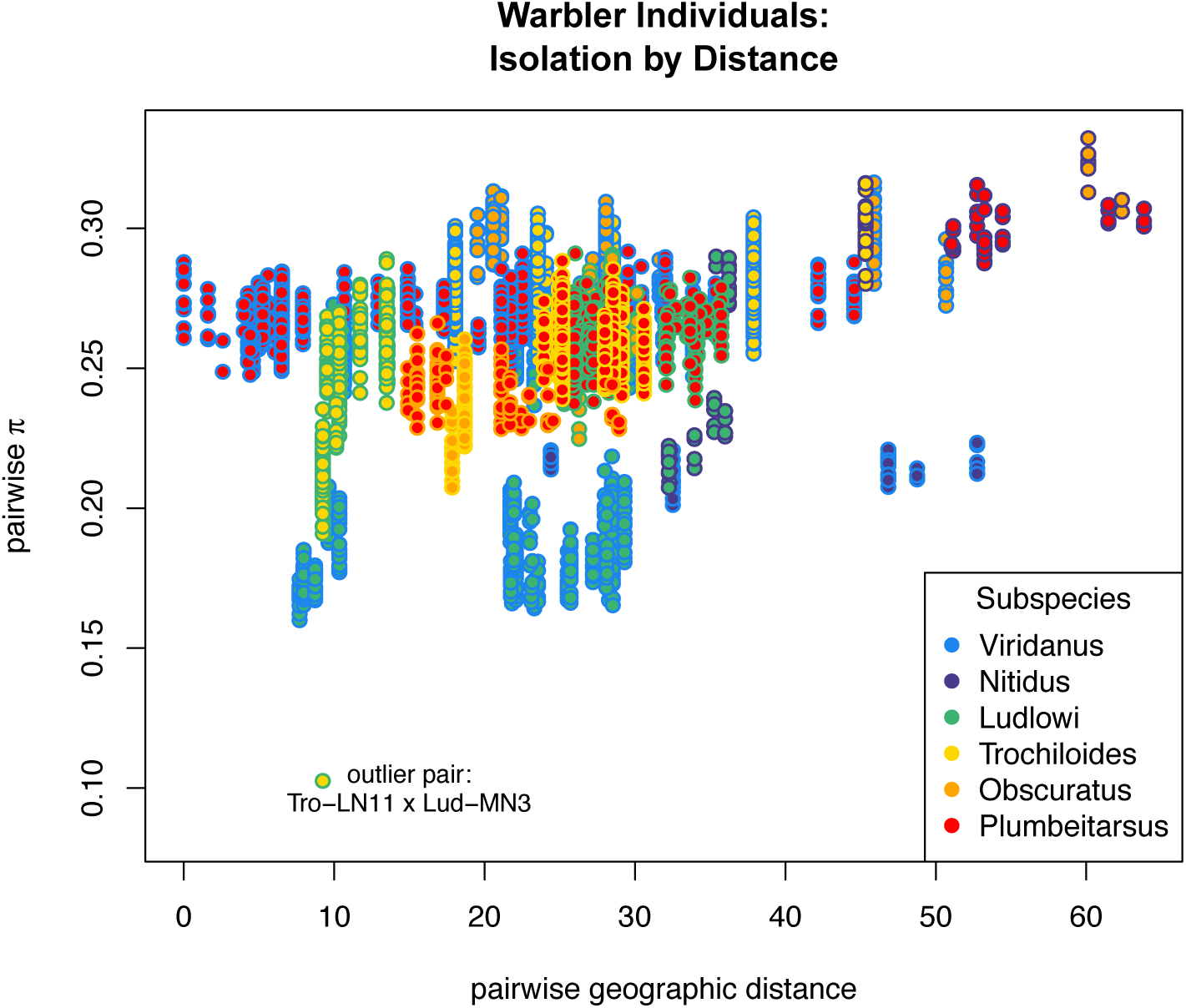
Mean pairwise sequence divergence at polymorphic sites calculated between all pairs of individuals from different subspecies, and colored by the subspecies to which each individual in the comparison is drawn. Note that individuals Tro-LN11 and Lud-MN3 have sequence divergence that is unusually low relative to that of other comparisons between individuals from the same two subspecies.

**Figure S14:**
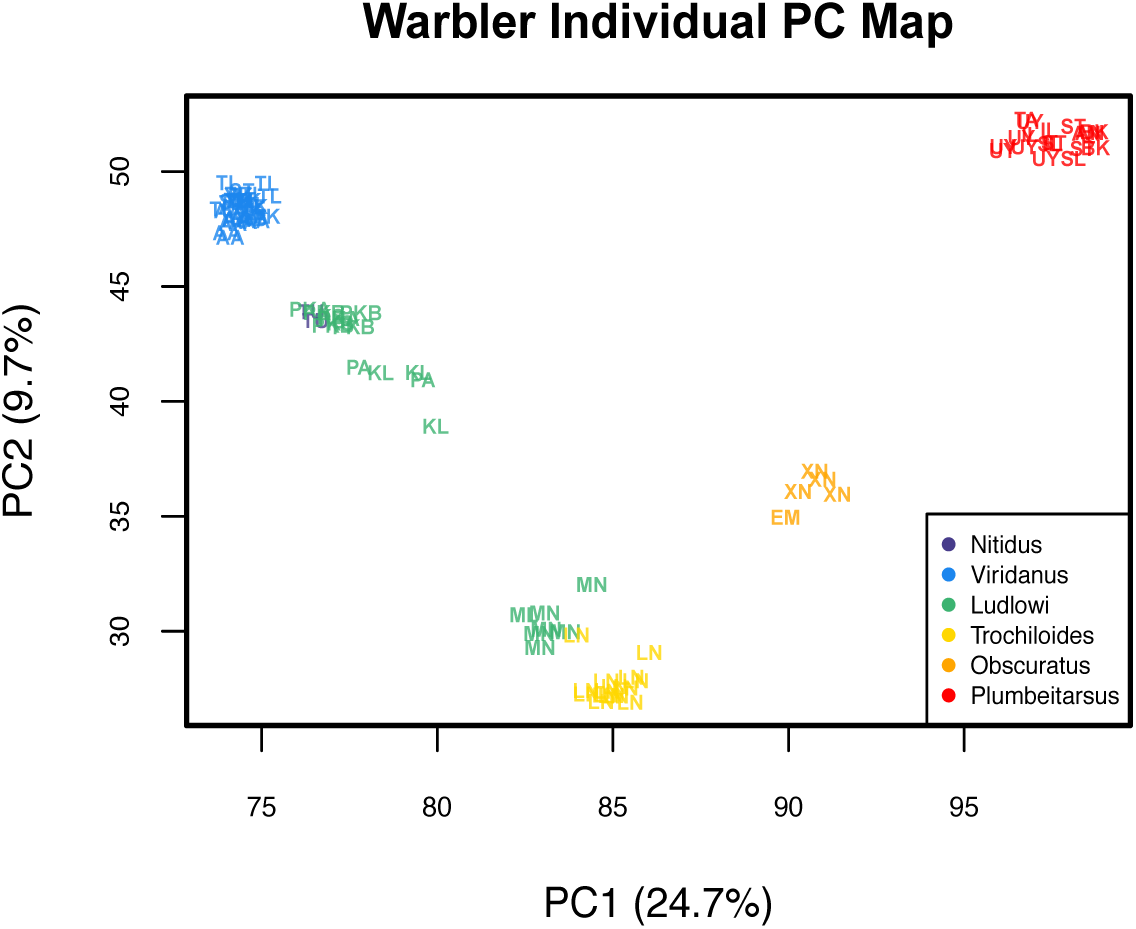
The map of warbler individuals derived from a Principal Components analysis. The PC coordinates have undergone a full Procrustes transformation around the actual sampling coordinates.

**Figure S15:**
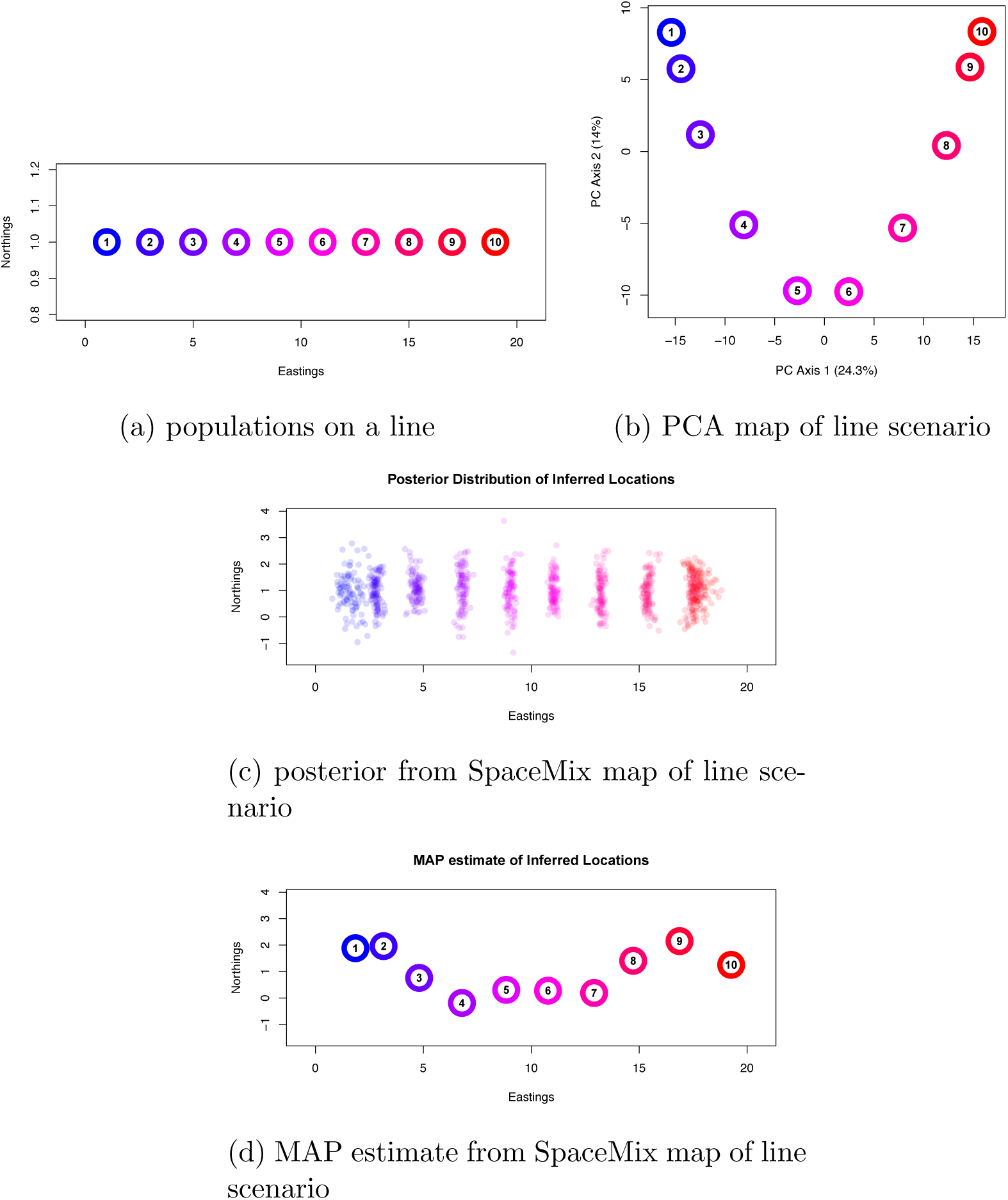
Simulation scenario of populations on a line, contrasting PCA-based inference and SpaceMix inference. a) Scenario used to simulate data in a spatial coalescent framework with nearest-neighbor migration; b) PCA map of allele frequencies, plotting PC axis 1 against PC axis 2, forming a ‘U’ shape; c) Posterior distribution of SpaceMix location inference, forming a rough line; d) snapshot of the MAP draw from the posterior, again showing a rough line.

**Figure S16:**
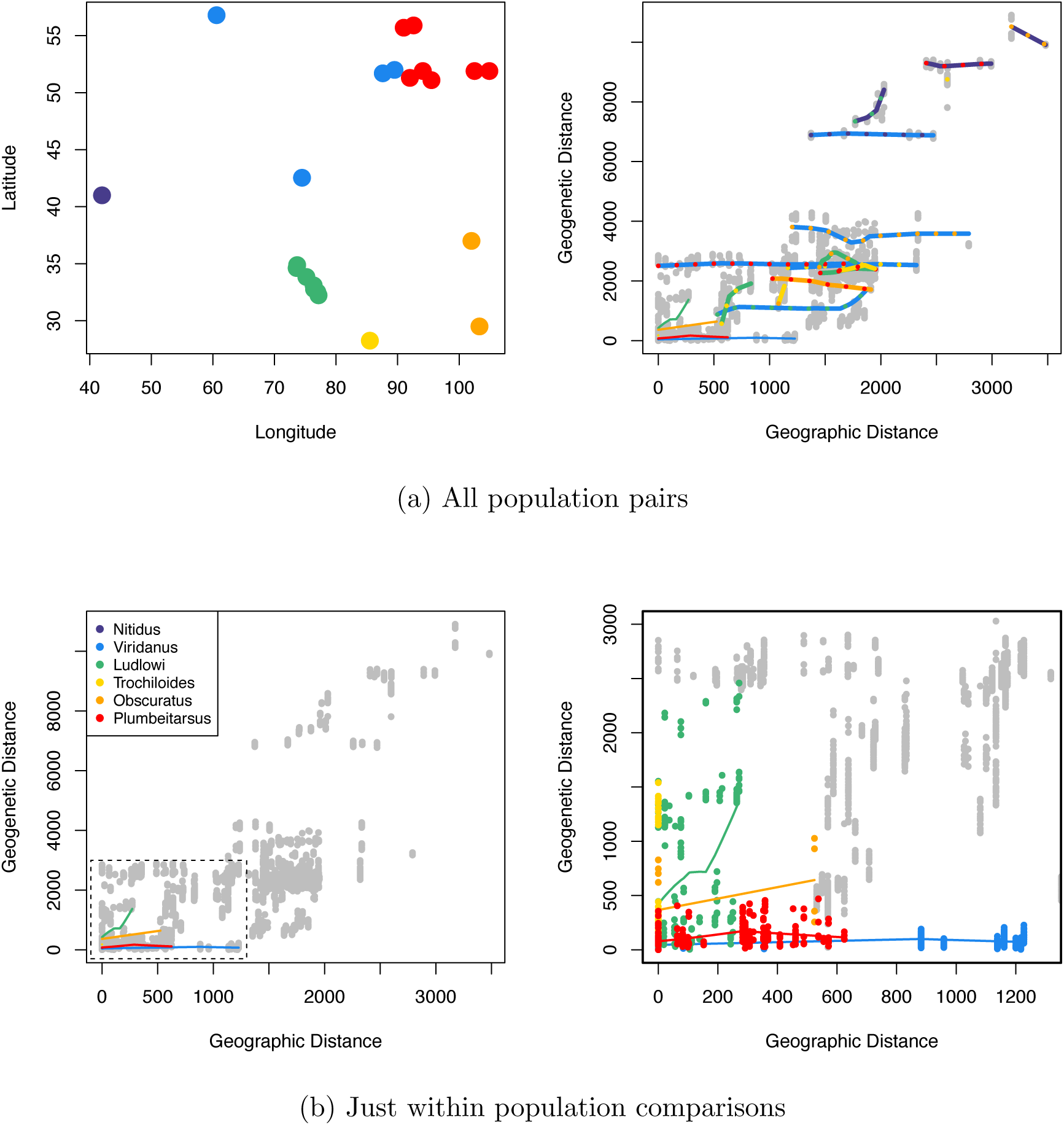
Comparing geographic pairwise distance to geogenetic pairwise distance between warbler individuals, (a) between and (b) within subspecies populations.

**Figure S17:**
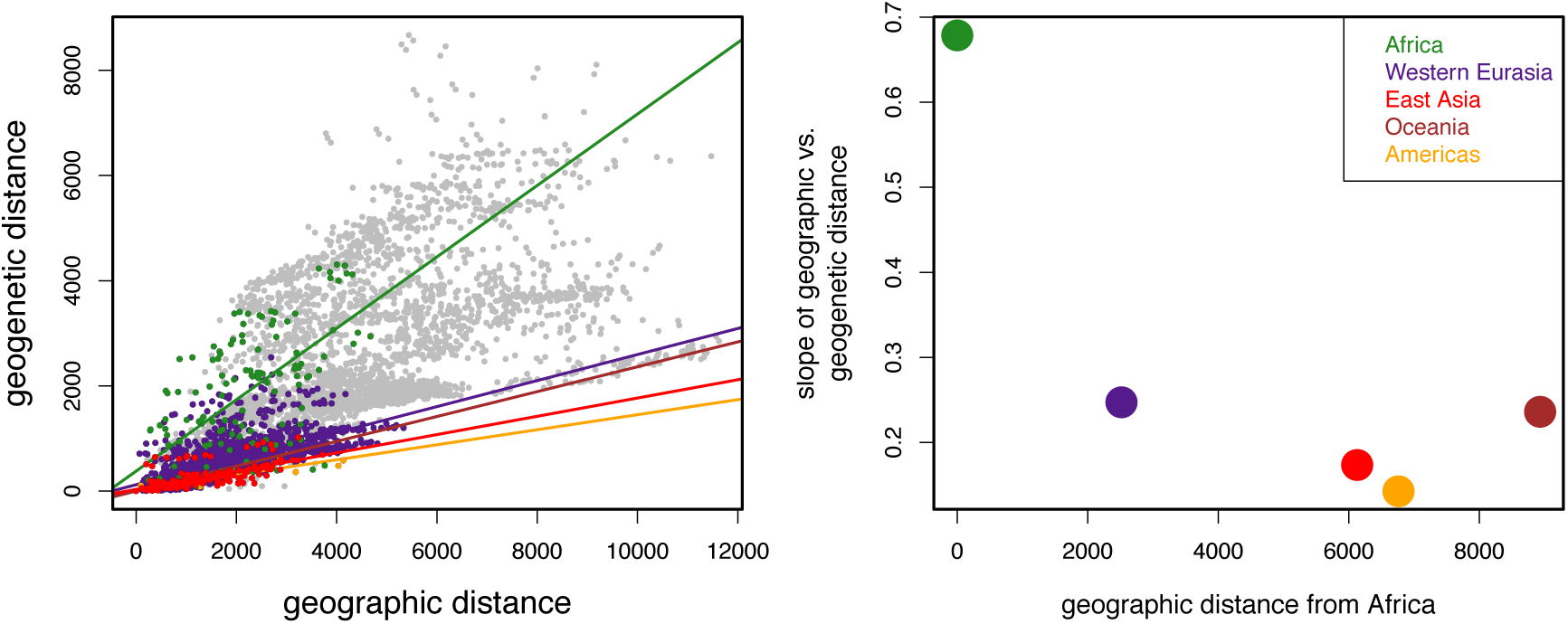
Comparison of geographic pairwise distance to geogenetic pairwise distance between human populations, colored by continent from which populations were sampled (i.e., two populations sampled from Africa are green). Eurasia is divided into Western Eurasia and East Asia.

**Figure S18:**
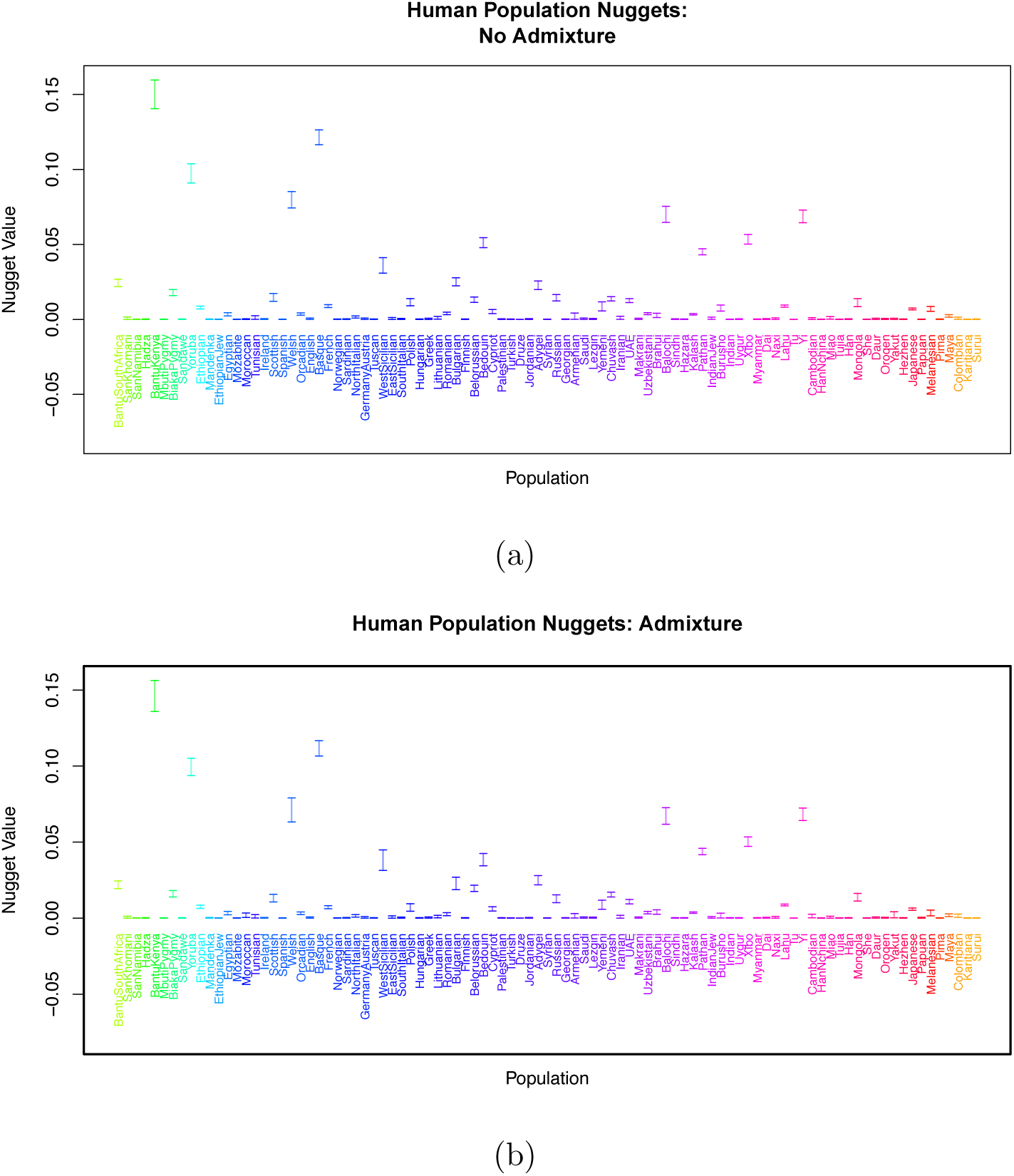
Credible intervals on estimated human sample nugget parameters. analysis without admixture; a) analysis with admixture.

**Figure S19:**
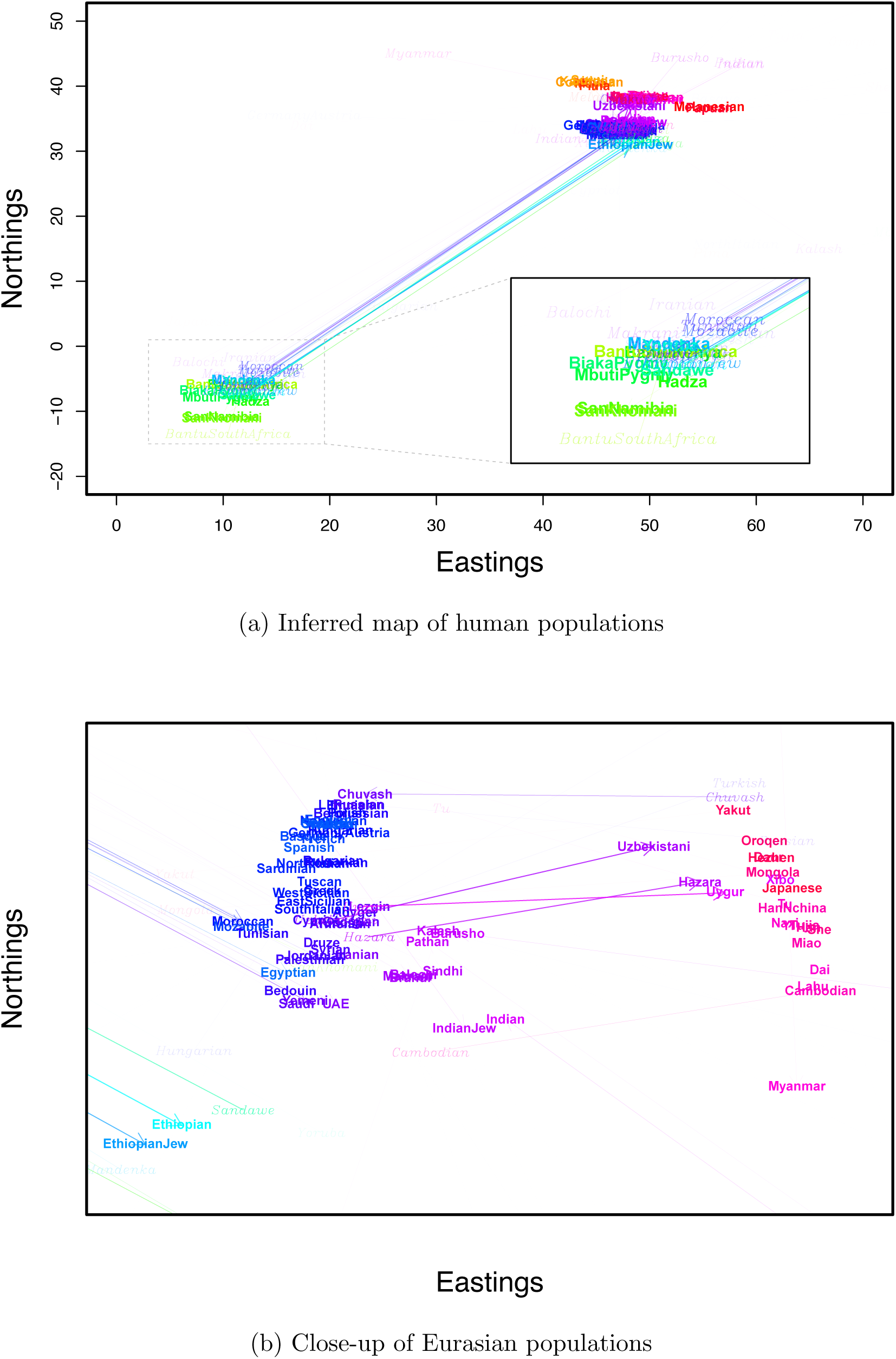
Map of human populations from a different SpaceMix analysis

**Figure S20:**
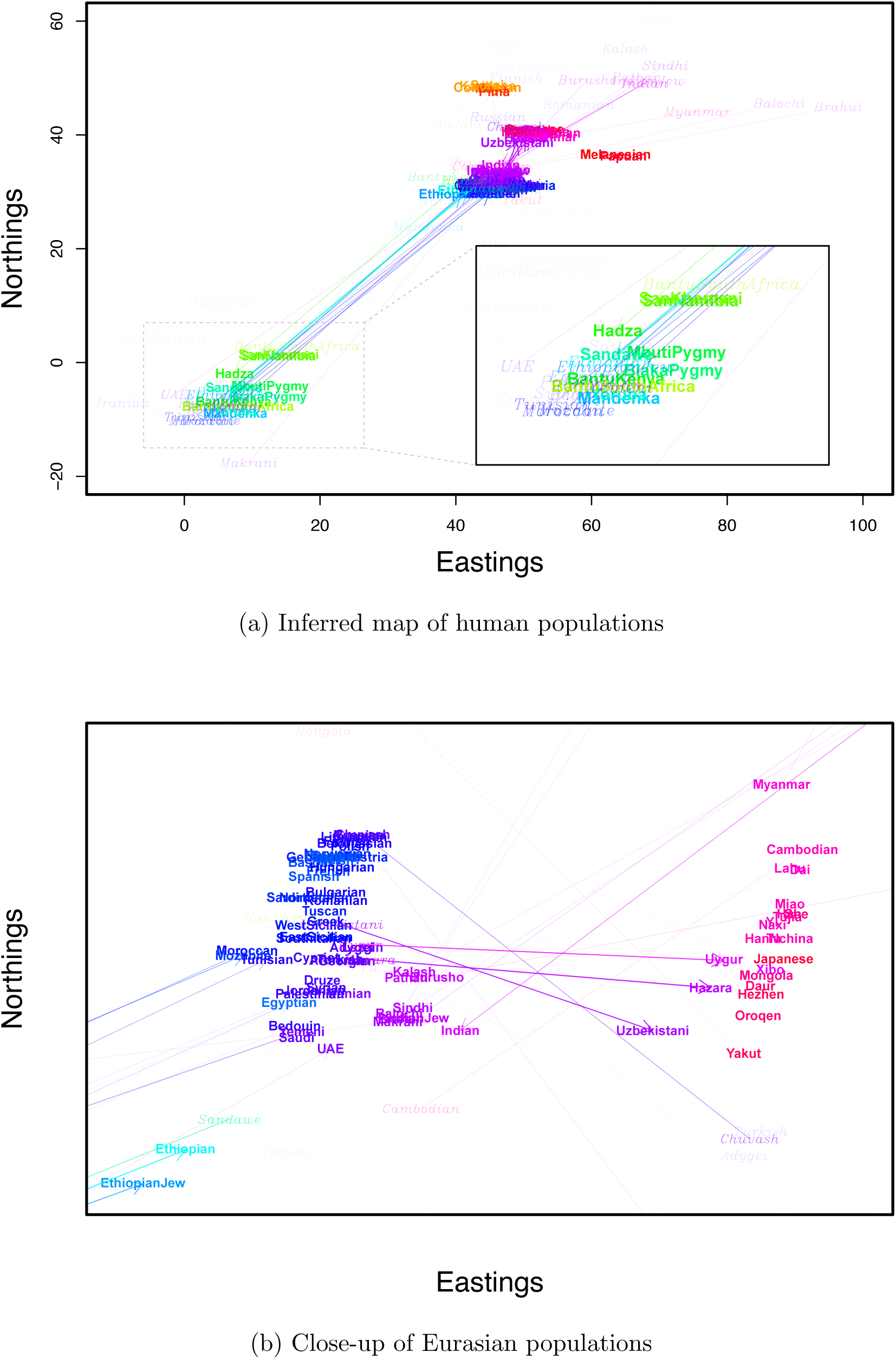
Map of human populations from another SpaceMix analysis

**Figure S21:**
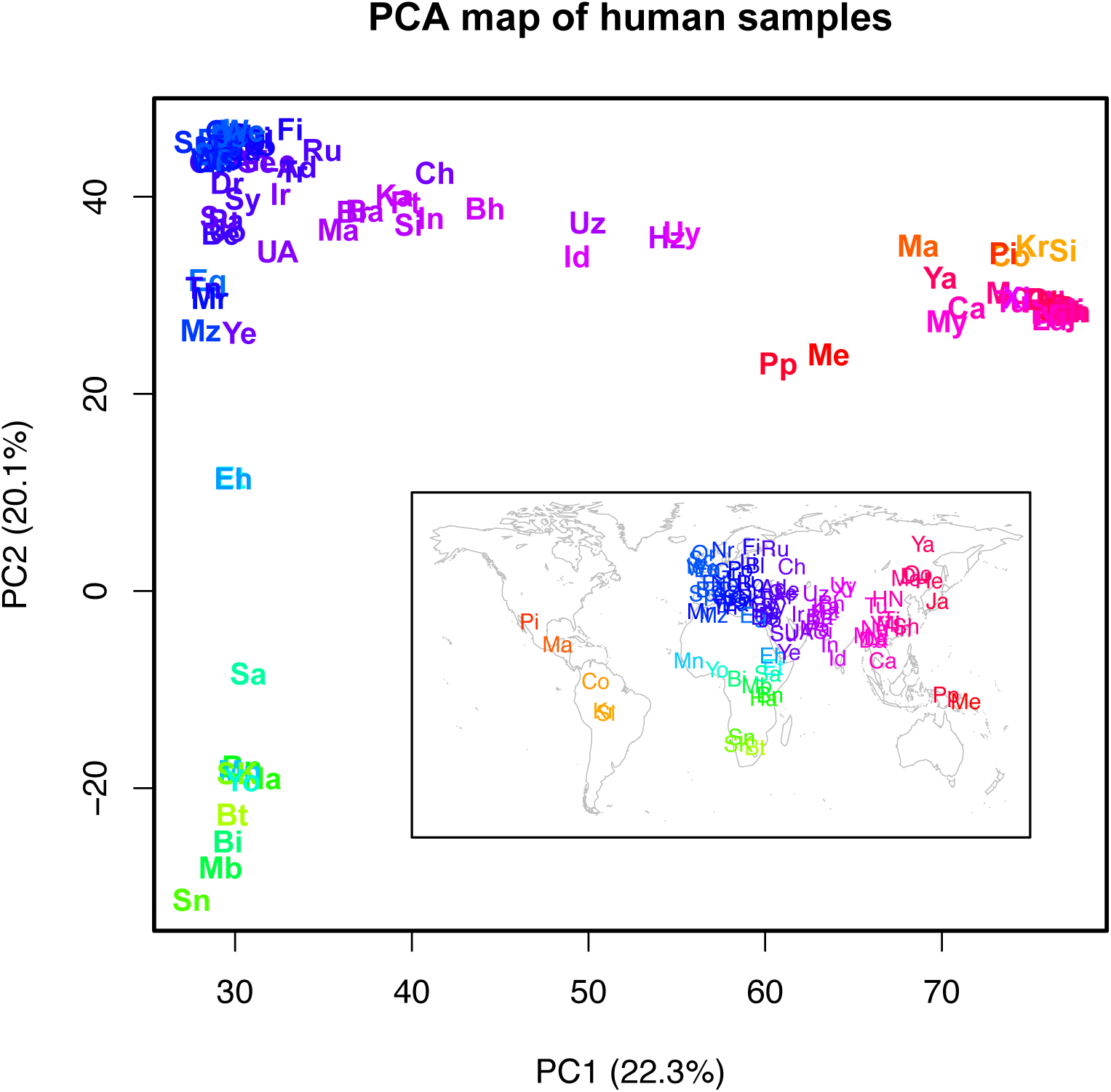
PCA map of human samples used in SpaceMix analyses. The PC coordinates have undergone a full Procrustes transformation around the actual sampling coordinates (shown in the inset map).

**Figure S22:**
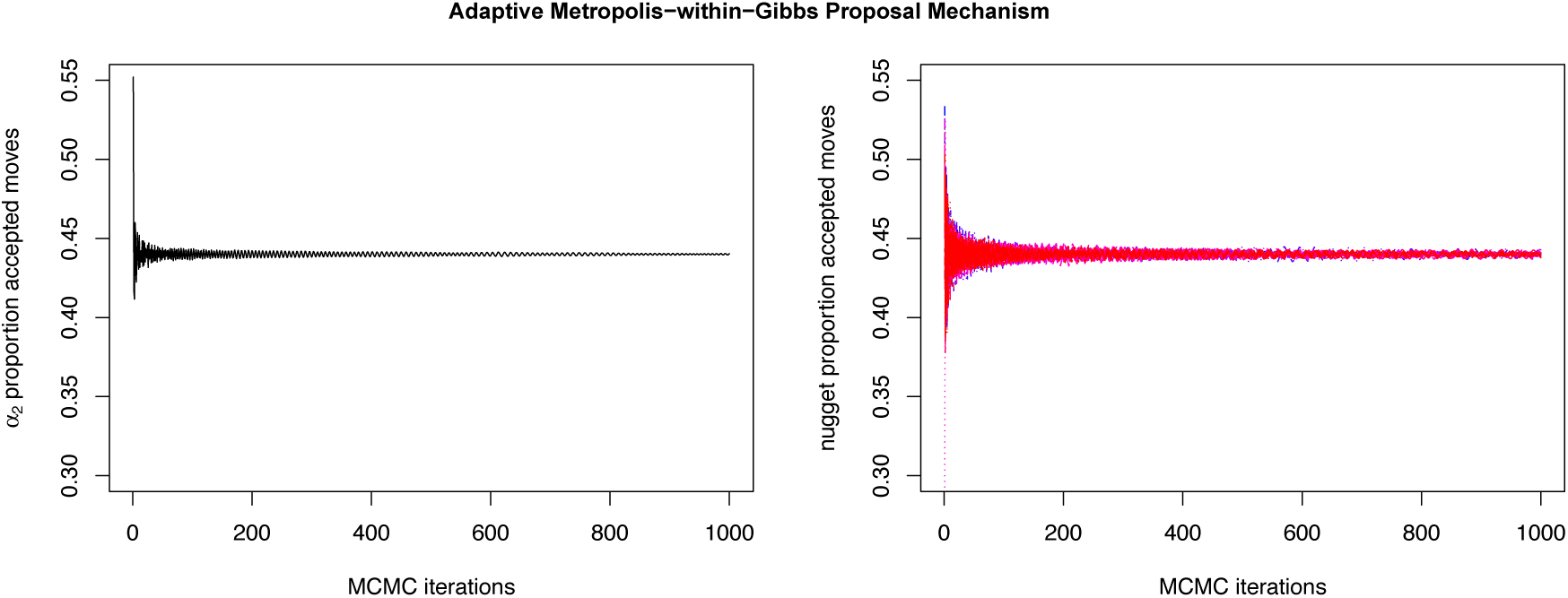
Example parameter acceptance proportions for the *α*_2_ parameter and the nugget parameter, *η*, using the adaptive Metropolis-within-Gibbs proposal mechanism.

**Figure S23:**
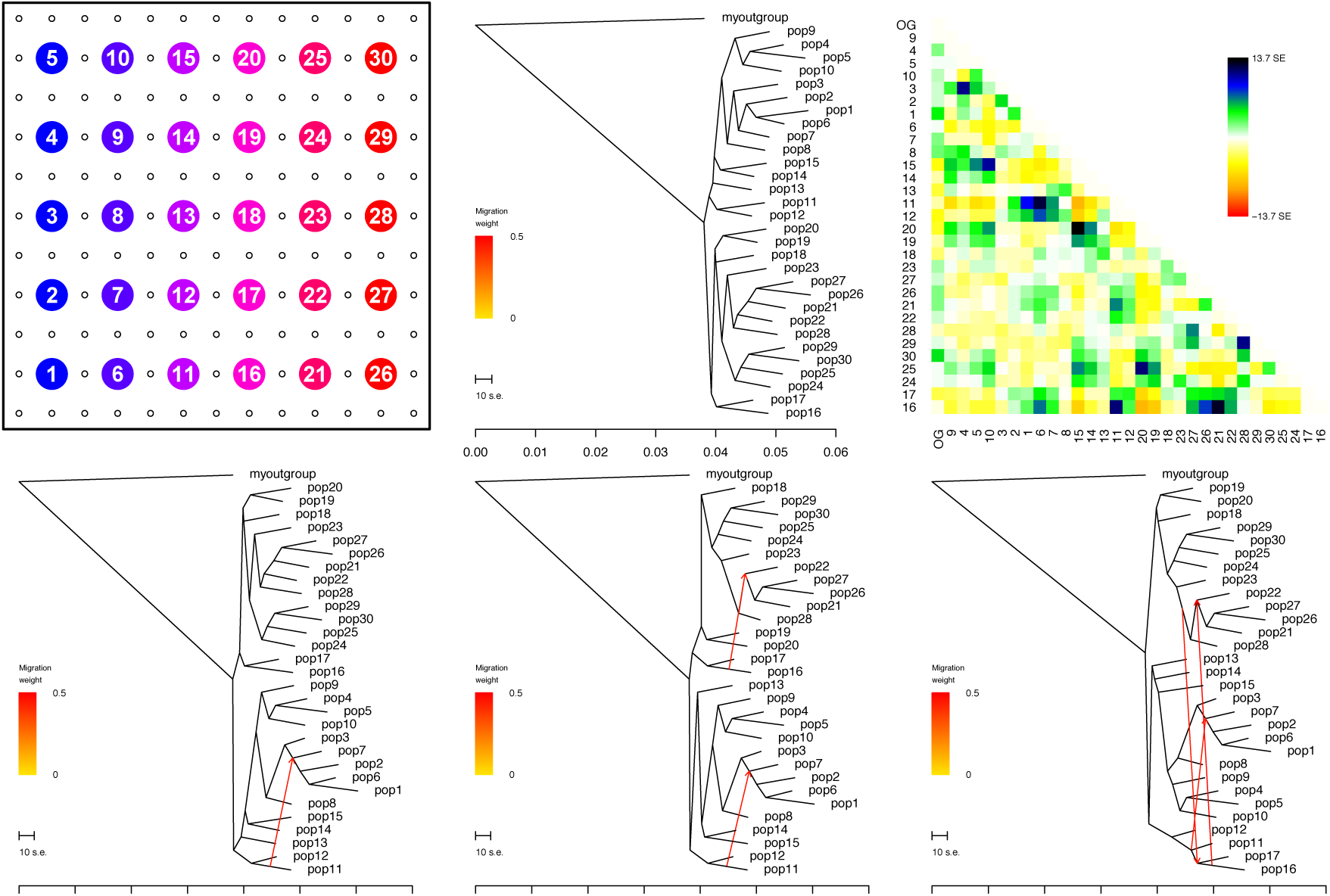
TreeMix analysis of lattice spatial coalescent simulation. The tree, residual covariance matrix, and first three migration admixture arrows are shown

**Figure S24:**
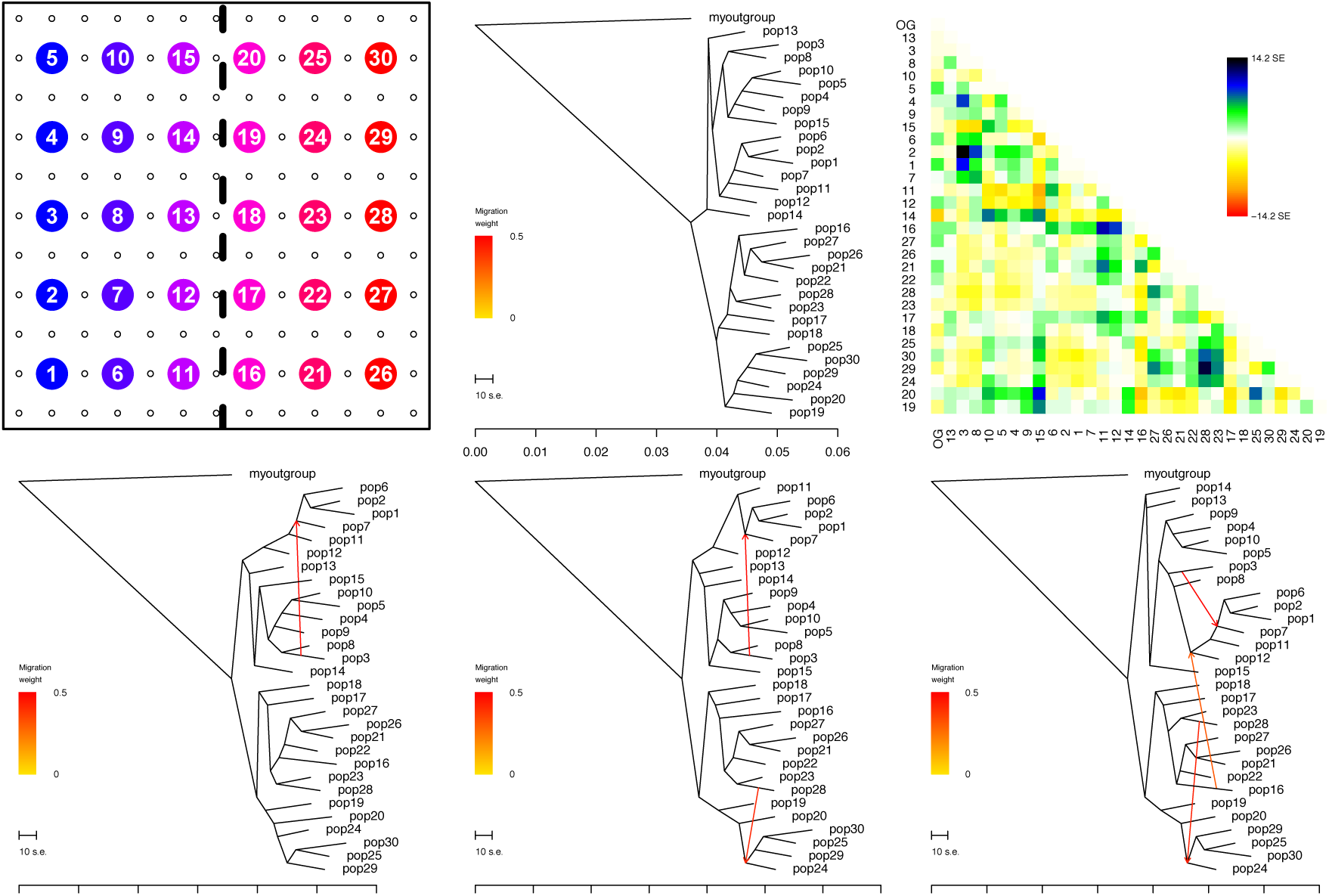
TreeMix analysis of lattice spatial coalescent simulation with a barrier.. The tree, residual covariance matrix, and first three migration admixture arrows are shown

**Figure S25:**
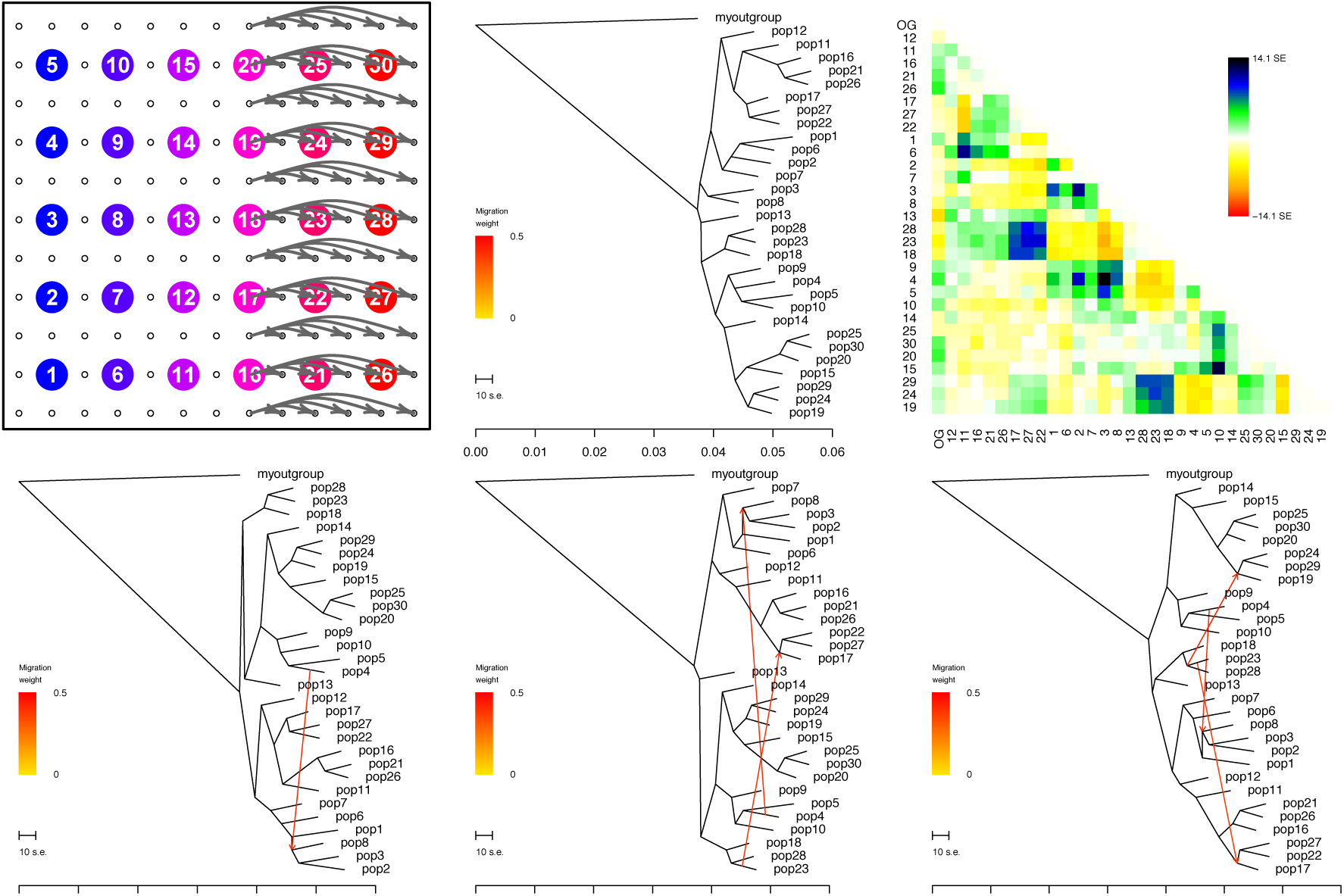
TreeMix analysis of lattice spatial coalescent simulation with an expansion event. The tree, residual covariance matrix, and first three migration admixture arrows are shown

**Figure S26:**
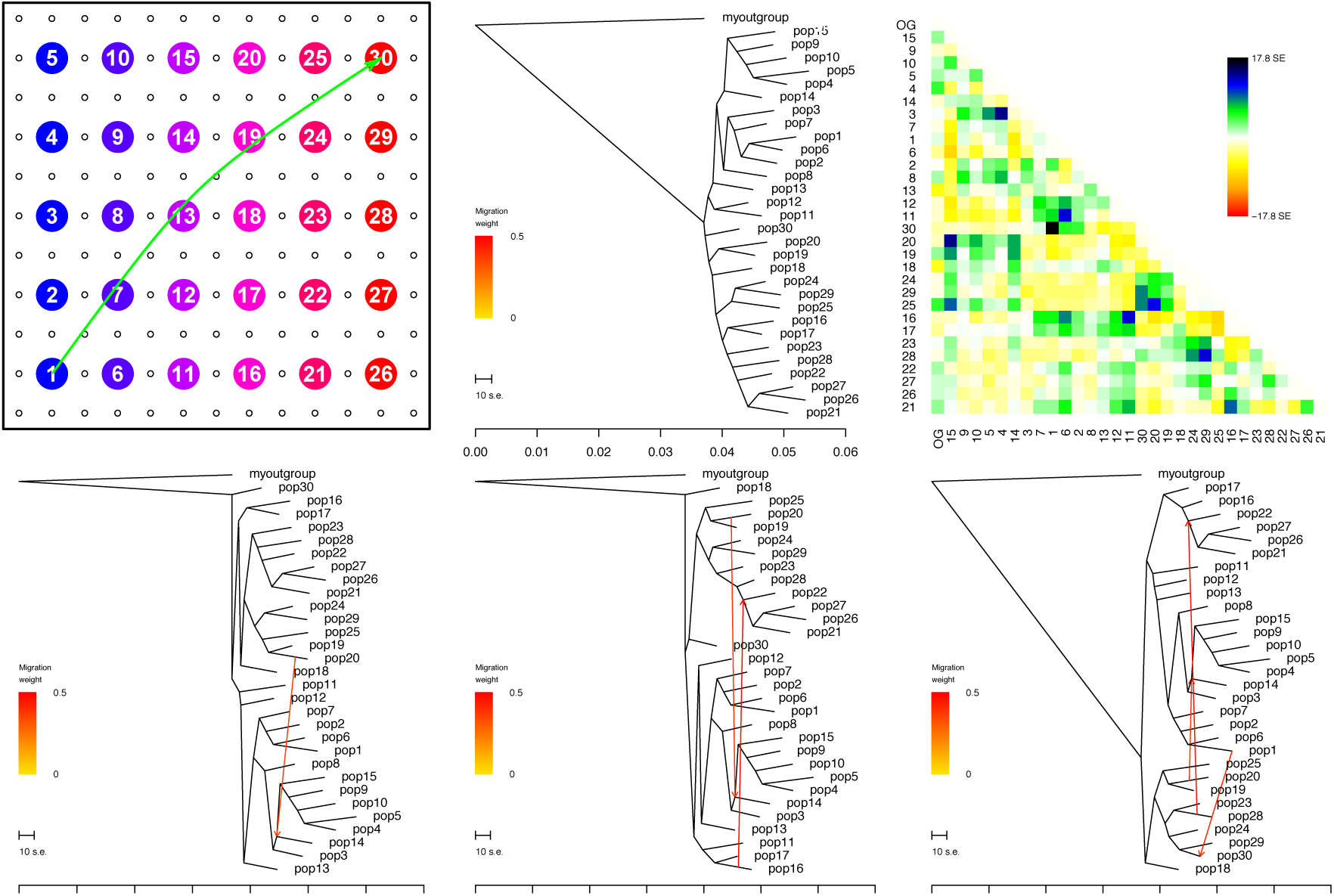
TreeMix analysis of lattice spatial coalescent simulation with a longrange admixture event. The tree, residual covariance matrix, and first three migration admixture arrows are shown

**Figure S27:**
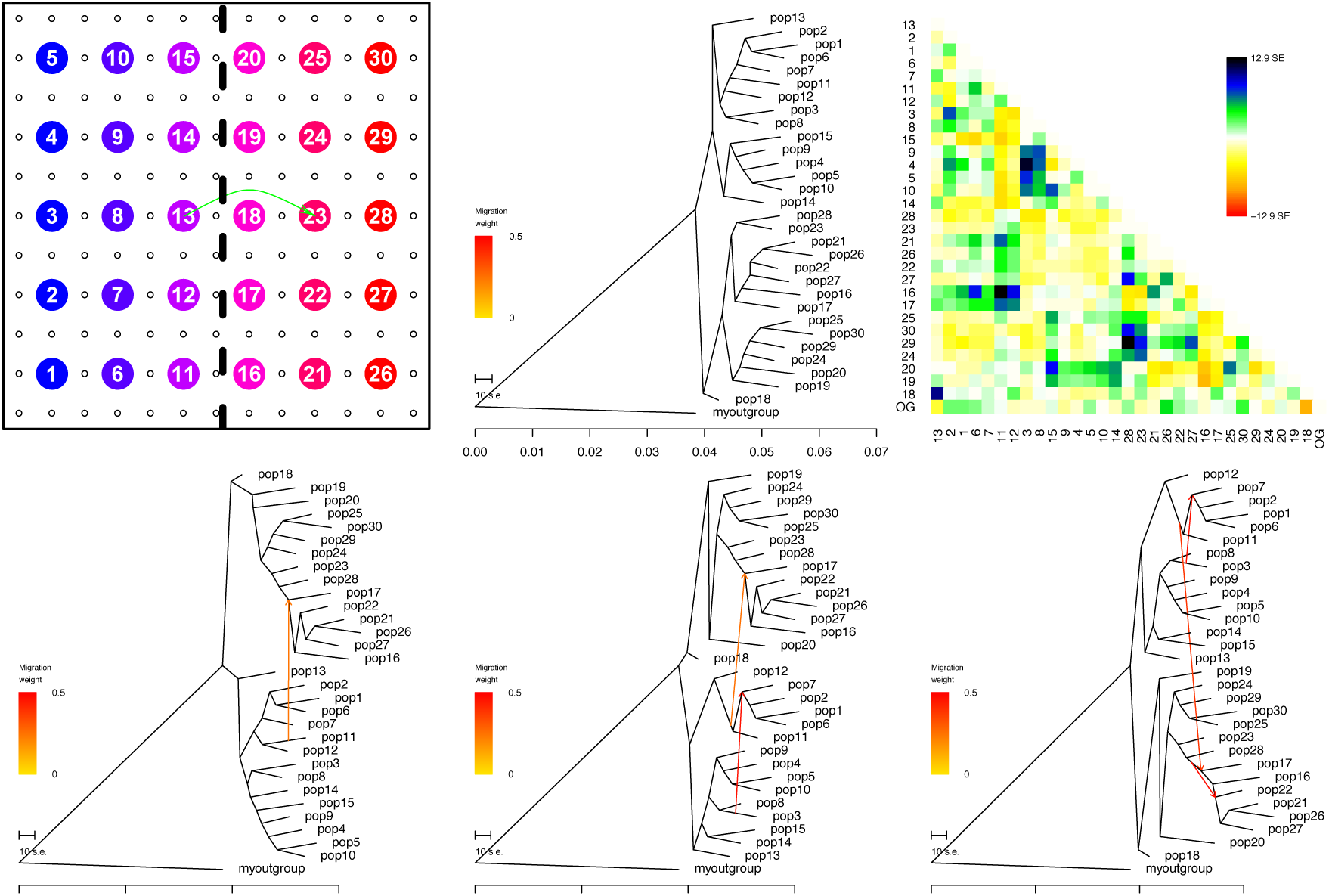
TreeMix analysis of lattice spatial coalescent simulation with a barrier and short-range admixture event. The tree, residual covariance matrix, and first three migration admixture arrows are shown

**Figure S28:**
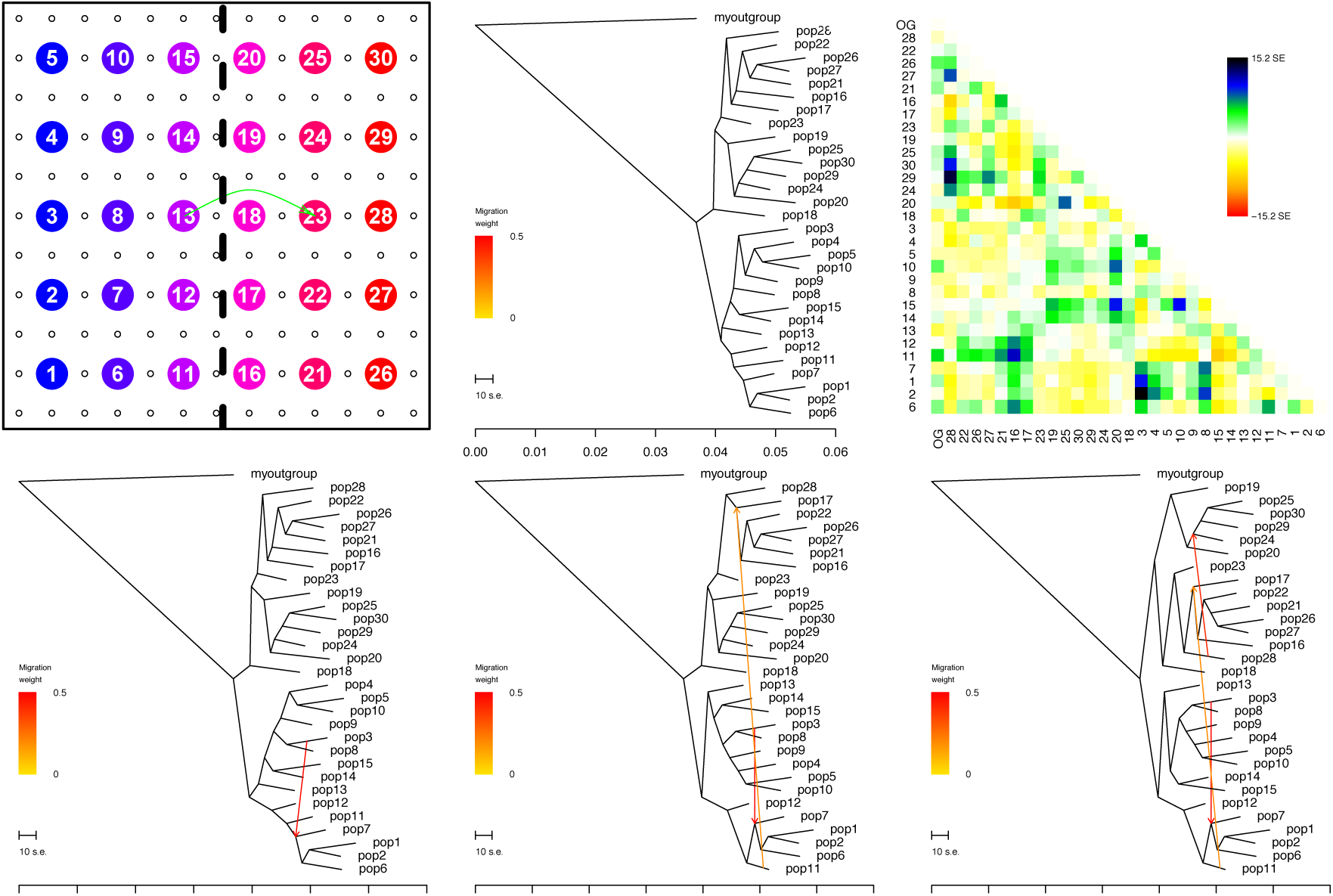
TreeMix analysis of lattice spatial coalescent simulation with a barrier and mid-range admixture event. The tree, residual covariance matrix, and first three migration admixture arrows are shown

**Table S1:** Subspecies and geographic meta-data for greenish warbler individuals included in analysis

| Sample | Subspecies | Longitude | Latitude |
| --- | --- | --- | --- |
| Vir-YK1 | Viridanus | 60.60 | 60.60 |
| Vir-YK11 | Viridanus | 60.60 | 60.60 |
| Vir-YK3 | Viridanus | 60.60 | 60.60 |
| Vir-YK4 | Viridanus | 60.60 | 60.60 |
| Vir-YK5 | Viridanus | 60.60 | 60.60 |
| Vir-YK6 | Viridanus | 60.60 | 60.60 |
| Vir-YK7 | Viridanus | 60.60 | 60.60 |
| Vir-YK9 | Viridanus | 60.60 | 60.60 |
| Vir-AB1 | Viridanus | 89.50 | 89.50 |
| Vir-AB2 | Viridanus | 89.50 | 89.50 |
| Vir-STvi1 | Viridanus | 92.60 | 92.60 |
| Vir-STvi2 | Viridanus | 92.60 | 92.60 |
| Vir-STvi3 | Viridanus | 92.60 | 92.60 |
| Vir-TL1 | Viridanus | 87.60 | 87.60 |
| Vir-TL10 | Viridanus | 87.60 | 87.60 |
| Vir-TL11 | Viridanus | 87.60 | 87.60 |
| Vir-TL12 | Viridanus | 87.60 | 87.60 |
| Vir-TL2 | Viridanus | 87.60 | 87.60 |
| Vir-TL3 | Viridanus | 87.60 | 87.60 |
| Vir-TL4 | Viridanus | 87.60 | 87.60 |
| Vir-TL5 | Viridanus | 87.60 | 87.60 |
| Vir-TL7 | Viridanus | 87.60 | 87.60 |
| Vir-TL8 | Viridanus | 87.60 | 87.60 |
| Vir-TL9 | Viridanus | 87.60 | 87.60 |
| Vir-AA1 | Viridanus | 74.48 | 74.48 |
| Vir-AA10 | Viridanus | 74.48 | 74.48 |
| Vir-AA11 | Viridanus | 74.48 | 74.48 |
| Vir-AA3 | Viridanus | 74.48 | 74.48 |
| Vir-AA4 | Viridanus | 74.48 | 74.48 |
| Vir-AA5 | Viridanus | 74.48 | 74.48 |
| Vir-AA6 | Viridanus | 74.48 | 74.48 |
| Vir-AA7 | Viridanus | 74.48 | 74.48 |
| Vir-AA8 | Viridanus | 74.48 | 74.48 |
| Vir-AA9 | Viridanus | 74.48 | 74.48 |
| Ni-TU1 | Nitidus | 42.00 | 42.00 |
| Ni-TU2 | Nitidus | 42.00 | 42.00 |
| Lud-PKA8 | Ludlowi | 73.69 | 73.69 |
| Lud-PKA10 | Ludlowi | 73.69 | 73.69 |
| Lud-PKB1 | Ludlowi | 73.61 | 73.61 |
| Lud-PKB2 | Ludlowi | 73.61 | 73.61 |
| Lud-PKB4 | Ludlowi | 73.61 | 73.61 |
| Lud-PKB5 | Ludlowi | 73.61 | 73.61 |
| Lud-KS1 | Ludlowi | 75.19 | 75.19 |
| Lud-KS2 | Ludlowi | 75.19 | 75.19 |
| Lud-KL6 | Ludlowi | 76.37 | 76.37 |
| Lud-KL7 | Ludlowi | 76.37 | 76.37 |
| Lud-KL1 | Ludlowi | 76.37 | 76.37 |
| Lud-PA1 | Ludlowi | 76.97 | 76.97 |
| Lud-PA2 | Ludlowi | 76.97 | 76.97 |
| Lud-ML2 | Ludlowi | 76.43 | 76.43 |
| Lud-MN1 | Ludlowi | 77.16 | 77.16 |
| Lud-MN12 | Ludlowi | 77.16 | 77.16 |
| Lud-MN3 | Ludlowi | 77.16 | 77.16 |
| Lud-MN5 | Ludlowi | 77.16 | 77.16 |
| Lud-MN8 | Ludlowi | 77.16 | 77.16 |
| Lud-MN9 | Ludlowi | 77.16 | 77.16 |
| Tro-LN1 | Trochiloides | 85.50 | 85.50 |
| Tro-LN10 | Trochiloides | 85.50 | 85.50 |
| Tro-LN11 | Trochiloides | 85.50 | 85.50 |
| Tro-LN12 | Trochiloides | 85.50 | 85.50 |
| Tro-LN14 | Trochiloides | 85.50 | 85.50 |
| Tro-LN16 | Trochiloides | 85.50 | 85.50 |
| Tro-LN18 | Trochiloides | 85.50 | 85.50 |
| Tro-LN19 | Trochiloides | 85.50 | 85.50 |
| Tro-LN2 | Trochiloides | 85.50 | 85.50 |
| Tro-LN20 | Trochiloides | 85.50 | 85.50 |
| Tro-LN3 | Trochiloides | 85.50 | 85.50 |
| Tro-LN4 | Trochiloides | 85.50 | 85.50 |
| Tro-LN6 | Trochiloides | 85.50 | 85.50 |
| Tro-LN7 | Trochiloides | 85.50 | 85.50 |
| Tro-LN8 | Trochiloides | 85.50 | 85.50 |
| Obs-EM1 | Obscuratus | 103.30 | 103.30 |
| Obs-XN1 | Obscuratus | 102.00 | 102.00 |
| Obs-XN2 | Obscuratus | 102.00 | 102.00 |
| Obs-XN3 | Obscuratus | 102.00 | 102.00 |
| Obs-XN5 | Obscuratus | 102.00 | 102.00 |
| Plu-BK2 | Plumbeitarsus | 104.90 | 104.90 |
| Plu-BK3 | Plumbeitarsus | 104.90 | 104.90 |
| Plu-AN1 | Plumbeitarsus | 102.50 | 102.50 |
| Plu-AN2 | Plumbeitarsus | 102.50 | 102.50 |
| Plu-IL1 | Plumbeitarsus | 95.50 | 95.50 |
| Plu-IL2 | Plumbeitarsus | 95.50 | 95.50 |
| Plu-IL4 | Plumbeitarsus | 95.50 | 95.50 |
| Plu-ST1 | Plumbeitarsus | 92.60 | 92.60 |
| Plu-ST12 | Plumbeitarsus | 92.60 | 92.60 |
| Plu-ST3 | Plumbeitarsus | 92.60 | 92.60 |
| Plu-UY1 | Plumbeitarsus | 94.10 | 94.10 |
| Plu-UY2 | Plumbeitarsus | 94.10 | 94.10 |
| Plu-UY3 | Plumbeitarsus | 94.10 | 94.10 |
| Plu-UY4 | Plumbeitarsus | 94.10 | 94.10 |
| Plu-UY5 | Plumbeitarsus | 94.10 | 94.10 |
| Plu-UY6 | Plumbeitarsus | 94.10 | 94.10 |
| Plu-SL1 | Plumbeitarsus | 91.00 | 91.00 |
| Plu-SL2 | Plumbeitarsus | 91.00 | 91.00 |
| Plu-TA1 | Plumbeitarsus | 92.00 | 92.00 |

**Table S2:** sample size and geographic meta-data for human samples included in analysis

|  | Population | Longitude | Latitude | Mean Sample Size |
| --- | --- | --- | --- | --- |
| 1 | BantuSouthAfrica | 28.00 | -26.00 | 15.99 |
| 2 | SanKhomani | 18.10 | -24.60 | 59.96 |
| 3 | SanNamibia | 20.00 | -21.50 | 9.99 |
| 4 | Hadza | 33.10 | -4.50 | 5.93 |
| 5 | BantuKenya | 37.00 | -3.00 | 21.99 |
| 6 | MbutiPygmy | 29.00 | 1.00 | 25.98 |
| 7 | BiakaPygmy | 17.00 | 4.00 | 41.97 |
| 8 | Sandawe | 35.70 | 6.20 | 55.94 |
| 9 | Yoruba | 5.00 | 8.00 | 41.98 |
| 10 | Ethiopian | 38.70 | 9.00 | 37.70 |
| 11 | Mandenka | -12.00 | 12.00 | 43.98 |
| 12 | EthiopianJew | 38.70 | 14.10 | 22.00 |
| 13 | Egyptian | 26.80 | 30.80 | 24.00 |
| 14 | Mozabite | 3.00 | 32.00 | 57.98 |
| 15 | Moroccan | -5.50 | 33.60 | 49.97 |
| 16 | Tunisian | 9.80 | 35.60 | 24.00 |
| 17 | Ireland | -8.20 | 53.40 | 14.00 |
| 18 | Scottish | -4.20 | 56.50 | 12.00 |
| 19 | Spanish | -3.70 | 40.50 | 67.95 |
| 20 | Welsh | -3.70 | 52.60 | 8.00 |
| 21 | Orcadian | -3.00 | 59.00 | 29.99 |
| 22 | English | -0.80 | 52.00 | 12.00 |
| 23 | Basque | 0.00 | 43.00 | 47.99 |
| 24 | French | 2.00 | 46.00 | 55.97 |
| 25 | Norwegian | 8.50 | 60.50 | 35.99 |
| 26 | Sardinian | 9.00 | 40.00 | 55.98 |
| 27 | NorthItalian | 9.70 | 45.70 | 23.99 |
| 28 | GermanyAustria | 10.50 | 51.20 | 8.00 |
| 29 | Tuscan | 11.00 | 43.00 | 16.00 |
| 30 | WestSicilian | 12.50 | 38.00 | 20.00 |
| 31 | EastSicilian | 16.10 | 37.00 | 20.00 |
| 32 | SouthItalian | 16.90 | 39.50 | 35.96 |
| 33 | Polish | 19.10 | 51.90 | 31.99 |
| 34 | Hungarian | 19.50 | 47.20 | 40.00 |
| 35 | Greek | 21.80 | 39.10 | 39.99 |
| 36 | Lithuanian | 23.90 | 55.20 | 20.00 |
| 37 | Romanian | 25.00 | 45.90 | 28.00 |
| 38 | Bulgarian | 25.50 | 42.70 | 35.99 |
| 39 | Finnish | 25.70 | 61.90 | 4.00 |
| 40 | Belorussian | 28.00 | 53.70 | 16.00 |
| 41 | Bedouin | 33.50 | 31.00 | 89.98 |
| 42 | Cypriot | 33.50 | 35.50 | 24.00 |
| 43 | Palestinian | 35.00 | 33.50 | 91.95 |
| 44 | Turkish | 35.20 | 39.00 | 34.00 |
| 45 | Druze | 37.00 | 32.00 | 83.96 |
| 46 | Jordanian | 37.00 | 30.00 | 40.00 |
| 47 | Adygei | 39.00 | 44.00 | 33.99 |
| 48 | Syrian | 39.00 | 34.80 | 32.00 |
| 49 | Russian | 40.00 | 61.00 | 49.98 |
| 50 | Georgian | 44.60 | 41.80 | 39.99 |
| 51 | Armenian | 45.00 | 40.10 | 31.99 |
| 52 | Saudi | 45.10 | 23.90 | 20.00 |
| 53 | Lezgin | 47.50 | 43.00 | 35.96 |
| 54 | Yemeni | 48.50 | 15.60 | 13.99 |
| 55 | Chuvash | 50.20 | 53.20 | 34.00 |
| 56 | Iranian | 53.70 | 32.40 | 39.99 |
| 57 | UAE | 54.40 | 24.50 | 27.98 |
| 58 | Makrani | 64.00 | 26.00 | 49.99 |
| 59 | Uzbekistani | 64.60 | 41.40 | 29.99 |
| 60 | Brahui | 65.00 | 29.00 | 49.98 |
| 61 | Balochi | 67.00 | 31.00 | 47.99 |
| 62 | Sindhi | 69.00 | 25.00 | 47.99 |
| 63 | Hazara | 69.50 | 33.00 | 43.98 |
| 64 | Kalash | 71.00 | 36.00 | 45.99 |
| 65 | Pathan | 72.50 | 34.00 | 43.99 |
| 66 | IndianJew | 72.90 | 19.00 | 16.00 |
| 67 | Burusho | 74.00 | 37.00 | 49.98 |
| 68 | Indian | 77.60 | 13.00 | 25.97 |
| 69 | Uygur | 81.00 | 44.00 | 20.00 |
| 70 | Xibo | 81.00 | 43.00 | 17.99 |
| 71 | Myanmar | 96.00 | 21.90 | 5.99 |
| 72 | Dai | 99.00 | 21.00 | 19.98 |
| 73 | Naxi | 100.00 | 26.00 | 15.99 |
| 74 | Lahu | 101.00 | 22.00 | 16.00 |
| 75 | Tu | 101.00 | 36.00 | 20.00 |
| 76 | Yi | 103.00 | 28.00 | 20.00 |
| 77 | Cambodian | 105.00 | 12.00 | 19.98 |
| 78 | HanNchina | 108.00 | 39.00 | 20.00 |
| 79 | Miao | 108.00 | 28.00 | 19.99 |
| 80 | Tujia | 110.00 | 29.00 | 20.00 |
| 81 | Han | 114.00 | 26.00 | 67.96 |
| 82 | Mongola | 119.00 | 48.00 | 20.00 |
| 83 | She | 119.00 | 27.00 | 19.99 |
| 84 | Daur | 124.00 | 49.00 | 17.99 |
| 85 | Oroqen | 126.00 | 50.00 | 18.00 |
| 86 | Yakut | 129.00 | 63.00 | 49.98 |
| 87 | Hezhen | 133.00 | 47.00 | 16.00 |
| 88 | Japanese | 138.00 | 38.00 | 55.97 |
| 89 | Papuan | 143.00 | -4.00 | 33.97 |
| 90 | Melanesian | 155.00 | -6.00 | 19.99 |
| 91 | Pima | -108.00 | 29.00 | 27.99 |
| 92 | Maya | -91.00 | 19.00 | 41.97 |
| 93 | Colombian | -68.00 | 3.00 | 13.99 |
| 94 | Karitiana | -63.00 | -10.00 | 27.99 |
| 95 | Surui | -62.00 | -11.00 | 16.00 |

